# Photoswitchable microtubule stabilisers optically control tubulin cytoskeleton structure and function

**DOI:** 10.1101/778993

**Authors:** Adrian Müller-Deku, Kristina Loy, Yvonne Kraus, Constanze Heise, Rebekkah Bingham, Julia Ahlfeld, Dirk Trauner, Oliver Thorn-Seshold

## Abstract

Small molecule inhibitors provide a versatile method for studies in microtubule cytoskeleton research, since tubulin is not readily amenable to functional control using genetics. However, traditional chemical inhibitors do not allow spatiotemporally precise applications on the length and time scales appropriate for selectively modulating microtubule-dependent processes. We have synthesised a panel of taxane-based light-responsive microtubule stabilisers, whose tubulin hyperpolymerisation activity can be induced by photoisomerisation to their thermodynamically metastable state. These reagents can be isomerised in live cells, optically controlling microtubule network integrity, cell cycle repartition, and cell survival, and offering biological response on the timescale of seconds and spatial precision to the level of individual cells. These azobenzene-based microtubule stabilisers offer the possibility of noninvasive, highly spatiotemporally precise modulation of the microtubule cytoskeleton in live cells, and can prove powerful reagents for studies of intracellular transport, cell motility, and neurodegeneration.

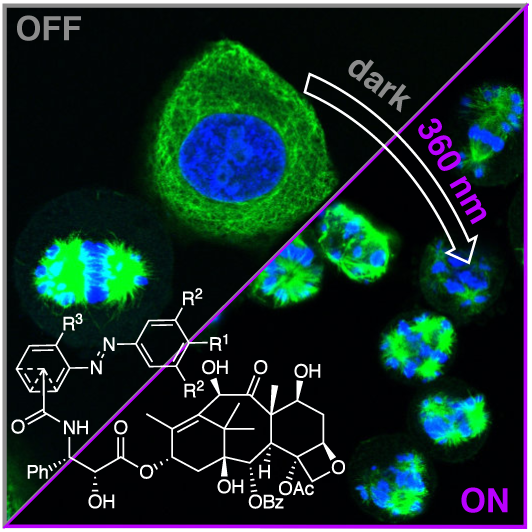

## Introduction

The cellular cytoskeleton, built from F-actin, microtubules and intermediate filaments, is a scaffold for critical biological processes from signaling and cargo trafficking, to cell shape maintenance and cell division. Most of the cytoskeleton’s myriad biological roles are inherently spatially and temporally differentiated, although all rely on the same three protein scaffold structures. Aiming to study these cytoskeleton-dependent processes with the spatiotemporal resolution appropriate to their biological function, a range of recent approaches have used optogenetic methods to photocontrol selected cytoskeleton-associated proteins, so enabling their spatiotemporally precise manipulation.^1–3^ More broadly however, direct spatiotemporal control over the structure and dynamics of the cytoskeleton scaffold proteins themselves, would offer a general approach to modulate any of their structure-dependent functions - although no genetic methods have yet been shown to deliver this.

We here focus on the microtubule (MT) cytoskeleton. MTs play particularly important roles in intracellular transport, cell motility, and structural plasticity, and there is an eminent need to achieve a better understanding of how these many functions are implemented and regulated.^4^ The role of MT dynamics during cell proliferation has also made them a major anticancer target, for which several outstanding drugs (taxanes, epothilones, vinca alkaloids) have been developed.^5–7^ In biological research, these drugs and other small molecule modulators (e.g. nocodazole, combretastatin, peloruside) remain the most general tools for MT cytoskeleton research. However, these inhibitors suppress all contemporaneous MT-dependent functions spatially indiscriminately: so they too do not allow spatiotemporally precise applications on the length and time scales appropriate for selectively studying MT-dependent processes. This restricts their scope of applications and their utility for selective research into MT cytoskeleton biology.^8^

Deeper insights could be gained from inhibitors that allow spatiotemporally specific MT manipulation. Photopharmaceuticals – photoswitchable (exogenous) small molecule inhibitors that act as an optically-controlled interface between a researcher and a protein of interest – have developed greatly in recent years.^9,10^ Photopharmaceuticals conceptually enable studies not otherwise accessible to biology, marrying the spatiotemporal precision of light application known from optogenetics, to the flexibility and system-independence of exogenous small molecule inhibitors, in a way that particularly favours noninvasive studies of temporally-regulated, spatially-anisotropic biological systems, such as the MT cytoskeleton.^9,11,12^ Photopharmaceuticals have succeeded in delivering a measure of optical control over a broad range of biochemical and biological phenomena, with early cell-free studies now supplanted by applications *in cellulo* and recently *in vivo*.^13–16^

In the cytoskeleton field, photopharmaceutical analogues of the MT destabiliser colchicine were recently developed, to begin addressing the need for spatiotemporally precise MT cytoskeleton studies. The azobenzene-based Photostatins (**PST**s), which can be reversibly photoswitched by low-intensity visible light between their biologically inactive *E*-isomers and their MT-inhibiting, colchicine-like *Z* isomers, were first reported in 2014.^9,17–19^ MT destabilising photopharmaceuticals based on two different families of molecular photoswitch – styrylbenzothiazoles (**SBTubs**)^20^ and hemithioindigos (bi-active **HOTubs**^21^ and dark-active **HITubs**)^22^ – have since been developed, delivering increased metabolic robustness in the intracellular environment as well as alternative optical switching profiles (all-visible switching with hemithioindigos, and GFP-orthogonal switching with **SBTubs**). All three reagent families have enabled highly spatiotemporally precise optical control over endogenous MT network integrity, MT polymerisation dynamics, cell division and cell death; and the **PST**s have already been used in animals to resolve outstanding questions e.g. in mammalian development^13,14^ and neuroscience^23^. These applications illustrate the power of photopharmacology to enable previously inaccessible studies of spatiotemporally anisotropic cytoskeletal processes without genetic engineering.^12–14,24^

With the optically precise blockage of MT polymerisation addressable by a range of photopharmaceuticals, we now desired to develop photopharmaceutical MT stabilisers (hyperpolymerisers) as conceptually novel tools with an alternative spectrum of biological research applications. The biological functions of the MT cytoskeleton are primarily driven by the localisation of stable or growing MTs themselves. Therefore, achieving optically-specific promotion of MT network integrity should allow spatiotemporally precise stimulation of MT-dependent functions, rather than localised blockade.

Such reagents could be applied to many biological questions, including further exploring recent discoveries of important roles of MTs in developing and regenerating neurons. While during development, MT stabilization has been shown to determine axonal identity and remodeling, MT stabilization in mature neurons seems to promote axonal regeneration by reducing the formation of retraction bulbs and modulating glial scar formation after spinal cord injury.^25–29^ However, the temporal characteristics of these phenomena are unclear, and the roles of MT stabilization in surrounding glia and immune cells rather than the damaged neurons themselves have not yet been resolved. Temporally-specific studies of the roles of dynamic and stabilised MTs would be highly desirable for studies in a range of other fields, such as in mitotic progression and immune cell response, particularly if stabilisation can be spatially targeted to selected cells within complex environments. Photopharmaceutical stabilisers could be used to trigger and study these phenomena with high temporal resolution and cell-level specificity, and would promise to shed new light on such roles of MTs in development, neuronal regeneration and repair.^25,30^

As far as we are aware, Liu and coworkers^31^ recently disclosed the only work in the direction of photopharmaceutical MT stabilisation, applying the photoswitchability of host-guest interactions between beta-cyclodextran and arylazopyrazoles (up to binding constant 2300 M^-1^)^32^ to aim at photoswitchably reversible, noncovalent heterodimerisation of two paclitaxel conjugates that should crosslink two MTs. The authors report that the heterodimeriser isomer-dependently alters the proportion of subG1-phase (dying) cells from 8% to 12% (*cis* and *trans* respectively, ± 2%) although *in cellulo* photoswitchability was not demonstrated. We considered that this work does not solve the need for a robust optical MT reagent, and features conceptual drawbacks. Crosslinking by the heterodimer (maximum taxol-to-taxol distance ca. 2 nm) of two MTs seems geometrically unlikely since taxol’s binding site is on the luminal (inner) face of the MTs^33^, giving a minimum taxol-to-taxol distance ca. 7 nm and requiring the linker to penetrate directly through both protein walls, as well as stretching substantially and reorienting the usual taxol binding pose. The lack of cellular potency of each monomer half is also striking (perhaps since the critical^7^ 2’-hydroxyl is capped with the photoswitch albeit in a way that may be enzymatically labile), and the additional possibility of isomer-dependent transmembrane trafficking of this ∼ 4 kDa construct complicates results. Such considerations (see discussion in Supporting Information) encouraged us instead to pursue lower-molecular-weight, druglike reagents that could offer robust, structurally rationalisable performance. We therefore chose to develop monomeric paclitaxel analogues incorporating azobenzenes that directly give photoswitchability of tubulin binding potency, as optically controlled MT stabilisers for *in situ* spatiotemporal photocontrol of MT network structure and function. We now report our development of these reagents.

## Results

### Design and Synthesis

We chose the azobenzene photoswitch for installing photoswitchable binding potency to the taxane core. This photoswitch offers a substantial geometric change upon isomerisation, which we hoped would differentiate the isomers’ binding constants, and allows reliable, high-quantum-yield, near-UV/visible-light-mediated, highly robust *E*↔*Z* photoisomerisability, which allows repeated *in cellulo* photoswitching. Taxanes feature a number of chemically modifiable positions; we chose to focus on sites where substituents can be tolerated, but where their geometric changes might impact binding potency through steric interactions. Potent taxanes feature a side-chain 3’-amine substituted with mid-size hydrophobic groups (e.g. Boc group in docetaxel, Bz in paclitaxel)^7,34^ which abut the tubulin protein surface yet are projected away from the protein interior (Fig 1a, highlighted in pink); the other sidechain positions (eg. the 3’-phenyl or 2’-hydroxyl) offer less spatial tolerance for substitution as they project into the protein^7^. The 3’-amine also tolerates the attachment of polar cargos such as the large silarhodamine fluorophore, as long as they are attached via a long spacer,^35^ making it desirable for photopharmaceutical tuning as it might tolerate azobenzenes with a range of structural characteristics.

**Figure 1.**
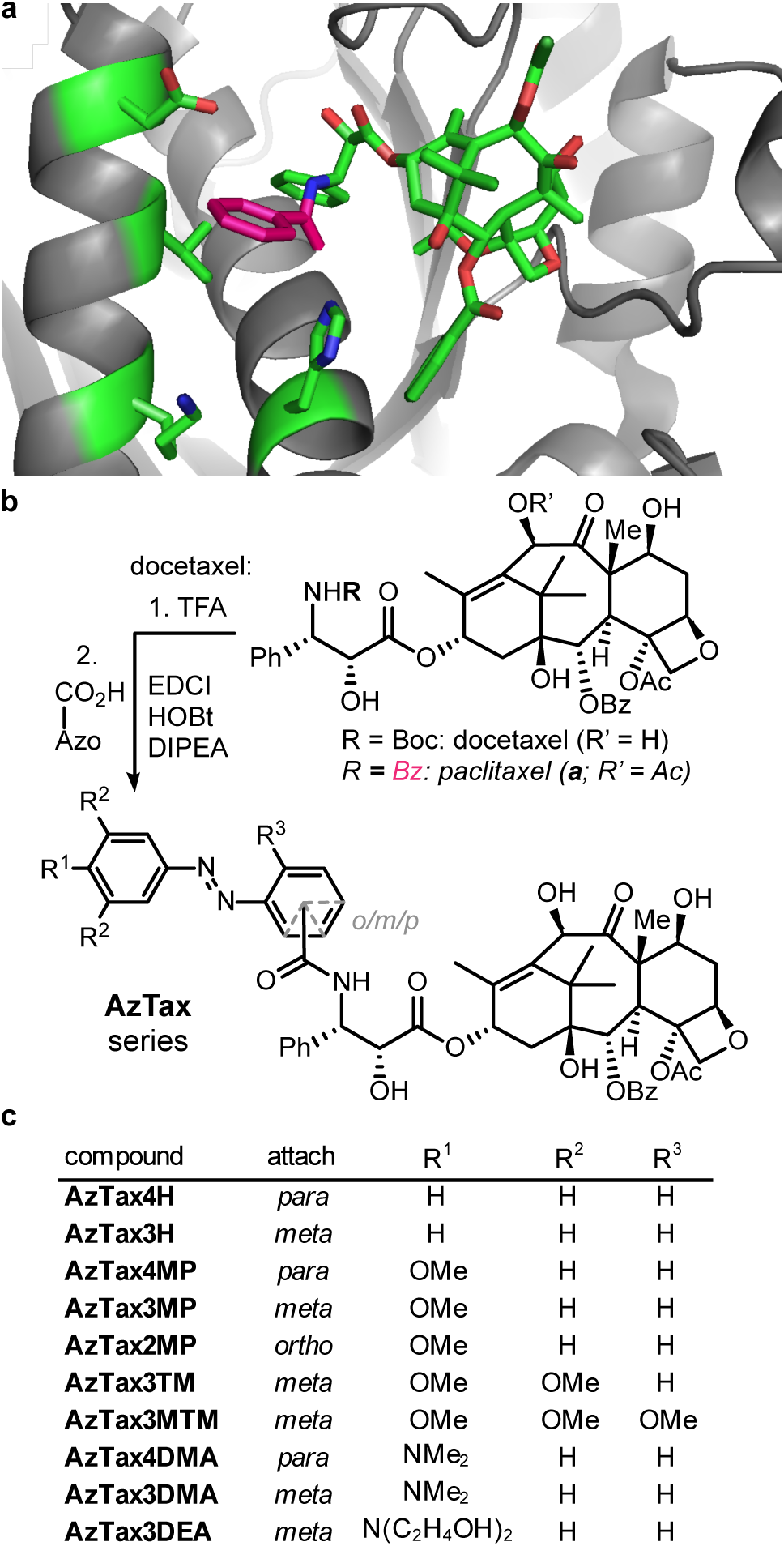
Design and synthesis of **AzTaxs. (a)** Paclitaxel:tubulin structure (PDB: 3J6G^33^) with the benzamide indicated in pink. **(b)** Synthesis of **AzTaxs** from docetaxel. **(c)** Panel of **AzTaxs** examined in this study.

We accordingly designed a panel of 3’-azoben-zamide-taxanes (**AzTaxs**) for biological testing. As taxanes have famously poor aqueous solubility (still worsened by attaching an azobenzene), we initially determined to focus on compounds displaying satisfactory potency at concentrations substantially below their solubility limit. This avoids the case that the compounds’ apparent potencies would be dictated by solubility effects, and so should enable robust use as reagents across a variety of systems and settings. Theorising that the sterics around the azobenzene phenyl ring proximal to the taxane core would be the greatest potency-affecting factor, we first focussed on testing which orientions of photoswitch would be best tolerated. We therefore scanned orientations of the diazene in *ortho, meta* and *para* relative to the amide (**AzTax2**/**3**/**4** compound sets), and when early testing showed that the **AzTax2** set had the lowest potency, we abandoned it at this stage. Next, examination of the published tubulin:paclitaxel cryo-EM structures^33,36^ indicated that the azobenzene’s distal ring can project freely away from the protein. Therefore we hypothesised that steric variation to the distal ring would not greatly impact binding potency of either isomer, but could be used orthogonally to tune their photochemical properties, by substitutions in *para* to the diazene that can mesomerically affect the properties of the N=N double bond. We accordingly synthesised unsubstituted (“H”), *para-*methoxy (“MP”), and *para-*dimethylamino (“DMA”) derivatives of the **AzTax3**/**4** sets. These were chosen to vary the photochemical properties of most relevance to photopharmacology: the completeness of the *E*→*Z* and the *Z*→*E* photoisomerisations at fixed wavelengths, which dictate the dynamic range of isomer photoswitchability, and τ (the halflife of the spontaneous unidirectional *Z*→*E* relaxation). Lastly, when the **AzTax3** set proved promising in early studies, we also examined installing an electron-donating 3,4,5-trimethoxy motif on the distal ring (**AzTax3TM**) as well as an additional R^3^ methoxy group to reduce the rotatability of the proximal ring in case this could amplify the difference between isomer potencies (**AzTax3MTM**), and we controlled for solubility effects by exchanging the dimethylamino substituent for a more soluble diethanolamino (**AzTax3DEA**). The target **AzTaxs** were synthesised by degradation of commercial docetaxel followed by amide couplings to various azobenzenecarboxylic acids in moderate yields (Fig 1b-c).

### Photochemical characterisation

The **AzTaxs** all displayed robust and repeatable *E*↔*Z* photoswitching under near-UV/visible illuminations, as expected from the literature^11^ (Fig S1). The photochemical properties within each substituent set were similar. The “H” compounds displayed a 3-fold dynamic range of *Z-*isomer photoswitchability between the photostationary states (PSSs) at 375 nm (80% *Z*) and 410 nm (26% *Z*), and had substantially slower relaxation than biological timescales (τ ca. 50 days). The methoxylated compounds (“MP”, “TM” and “MTM”) had been chosen to improve the dynamic range of photoswitching by relative shifting of the isomers’ absorption bands^9^ (Fig S2), and delivered a ca. 9-fold dynamic range of *Z-*isomer photoswitchability (375 nm: 96% *Z*; 530 nm: 11% *Z*); their relaxation remained substantially slower than biological timescales (τ ca. 24 h). The *para*-amino “DMA” and “DEA” compounds featured τ values too small to observe bulk photoswitching in aqueous media under biologically applicable conditions. Yet, as aprotic environments (such as lipid vesicles, membranes, and on-protein adsorbed states) are likely intracellular localisations for hydrophobic taxanes conjugates, we determined their photochemistry in moderately polar aprotic media (EtOAc). Here they were easily bulk-switchable (τ ca. 11 min), giving a 4-fold dynamic range of *Z-*isomer photoswitchability (410 nm: 91% *Z*; 530 nm: 21% *Z*) (further discussion in the Supporting Information). As the **AzTax** reagents were intended for use with microscopy, we also examined photoswitching over a broader range of wavelengths, to determine what dynamic range of isomer photoswitchability would be accessible in practice, with standard (405 nm, 488 nm, 514 nm) or more exotic (380 nm, 440 nm, 532 nm) microscopy laser wavelengths (Fig 2a, Fig S2, Table S1).

**Figure 2.**
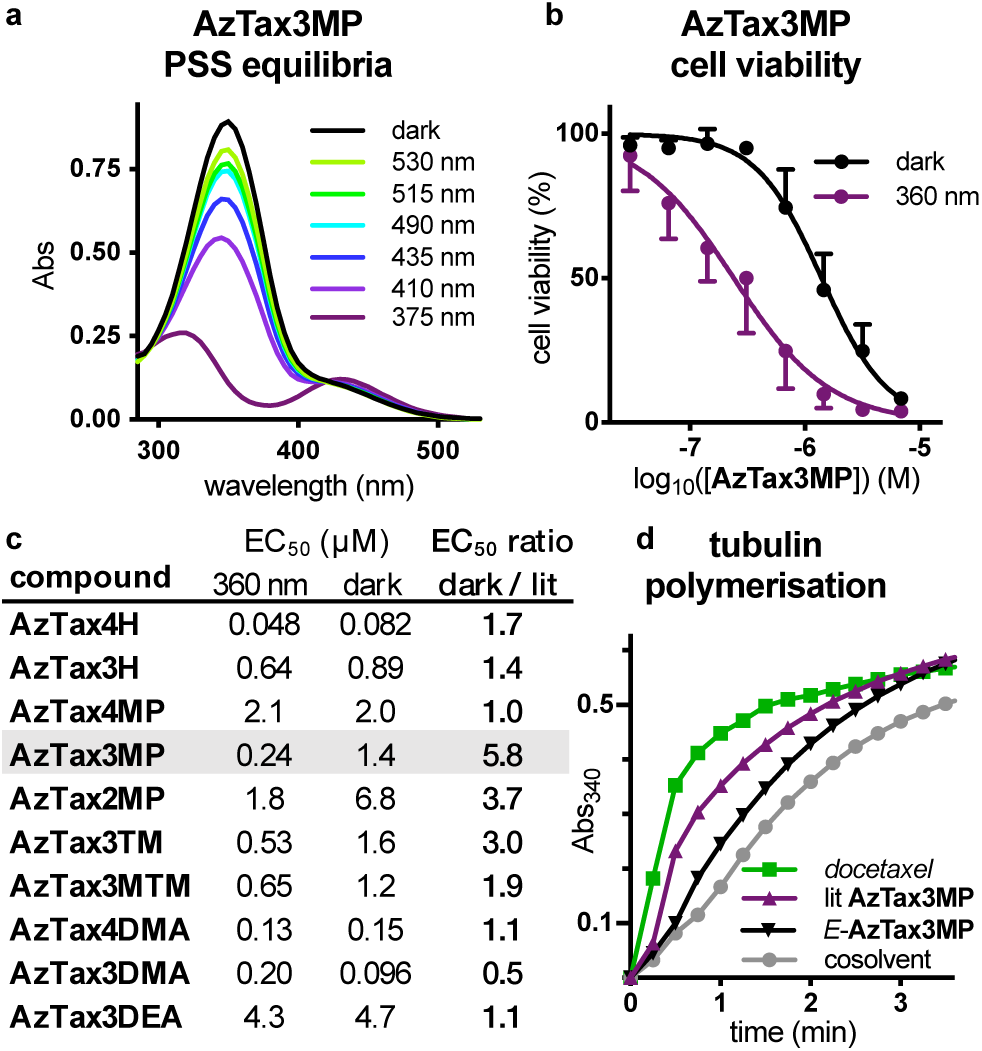
Photoswitchable performance of **AzTaxs. (a)** Photostationary state UV-Vis absorption spectra of **AzTax3MP** under a range of cell-compatible wavelengths similar to microscopy laser lines. **(b-c)** Resazurin antiproliferation assays of **AzTaxs** highlight their structure- and light-dependent cell cytotoxicity. HeLa cells, 40 h incubation in dark conditions (all-*E*) or under pulsed illuminations with low-power LEDs (75 ms per 15 s near-UV at <1 mW/cm2; “lit” = ∼ 80% Z). **(d)** A cell-free assay for polymerisation of purified tubulin comparing the MT stabilisation activity of docetaxel, all-*E*- and 360 nm-lit-**AzTax3MP** (all 10 µM) shows light-specific induction of hyperpolymerisation by *Z-***AzTax3MP**, matching the trend observed in cellular assays.

We next proceeded to explore the biological applicability of **AzTaxs** as photoswitchable MT stabilisers *in cellulo*. Since near-UV light gave PSSs with high-*Z* populations for all photoswitches, while thermal relaxation and maintenance in the dark returned the *E* isomer quantitatively, we began by comparing all-*E* “dark” (all-*E* stock applied, then maintained dark) with mostly-*Z* “360 nm” (all-*E* stock applied, then photoisomerised *in situ* by pulsed illuminations with low-power 360 nm LED light, giving a mostly-*Z* PSS) conditions, to determine which structures allowed the highest fold difference of bioactivity.

### AzTaxs display optically-controlled bioactivity *in cellulo*

Since stabilisation and hyperpolymerisation of MTs in cells over a prolonged period blocks cell proliferation and ultimately causes cell death^7^, we first assayed the **AzTaxs** for light-dependent cellular activity by the resazurin cell proliferation/viability assay. Viability dose-response curves under dark or UV conditions were assessed in the HeLa cervical cancer cell line (Fig 2b-c). All compounds displayed dose-response curves with similar Hill coefficients as the parent drug docetaxel (Fig S3), which is in line with the conjecture that they act through the same mechanism, albeit with different potency. All compounds except the fast-relaxing **AzTax3DMA** had *Z*-isomers that were more potent, or else equipotent, to the *E* isomers, suggesting that this trend in isomer-dependent cellular bioactivity across several photoswitch types has robust significance. *E-***AzTax2MP** had the poorest overall potency (EC_50_ ca. 7 µM, all-*E*), which we took as indicating the unsuitability of *ortho* substitutions that likely project the azobenzene into solution (c.f. Fig 1a) where its hydrophobicity can defavour binding stability. By contrast, the **AzTax4** set featured compounds up to 100 times more potent, and structure-dependently covered a 40-fold potency range. However, despite the good isomeric photoswitchability of e.g. **AzTax4MP**, none of the **AzTax4** set displayed substantial *photoswitchability of bioactivity* (fold difference between the 360 nm and the dark bioactivity). We interpreted this substitution-independent result as an indication that the distal ring may project too far from the protein contact surface, in both *E* and *Z* isomers, for the **AzTax4** isomer state to substantially affect binding.

In contrast, the **AzTax3** series all showed photoswitchability of bioactivity. **AzTax3MP** featured a nearly 6-fold difference between the more-toxic *Z* and less-toxic *E* isomers’ bioactivity, over the 48 h experimental timecourse (Fig 2b). Increasing the number of methoxy groups on the scaffold slightly decreased the *Z-*isomer’s cytotoxicity without greatly affecting that of the *E*-isomer (**AzTax3TM, AzTax3MTM**), and deleting the methoxy group also decreased the *Z-*isomer’s cytotoxicity (**AzTax3H**), which we took as a sign that balancing the photoswitch’s polarity was important for maximising bioactivity. In line with this interpretation, the potencies of **AzTax3DMA** were similar to **AzTax4DMA** while the more hydrophilic **AzTax3DEA** showed a 40-fold loss of potency. Despite the potential for photoswitching the *para-*amino **AzTax** inside lipid environments, they should only reach their cytosolic target tubulin as the *E*-isomers due to fast aqueous relaxation, yet surprisingly, **AzTax3DMA** appeared slightly more bioactive as the unilluminated *E* isomer, although as expected **AzTax4DMA** and **AzTax3DEA** both showed no illumination-dependency of bioactivity; and controls under 410 nm illumination (to establish the optimum PSS for the *para-*amino compounds) showed no different result to those obtained with UV illumination. The apparent cytotoxicity differential seen for **AzTax3DMA** might reflect reduced availability to the cytosol rather than differential binding of the isomers^22^ although these results do not allow further conjecture.

To continue the study we therefore selected **AzTax3MP**, due to its photoswitchability of bioactivity (6-fold), high potency (0.24 µM when UV-illuminated), bidirectional photoswitchability (optimum 9-fold-change of concentration of the more bio-active *Z* isomer), and reproducibly photoswitchable cellular performance across assays with different illumination conditions, and proceeded with further mechanistic biological evaluations.

### AzTaxs photocontrol tubulin polymerisation, cellular MT networks and cell cycle

To examine the molecular mechanism of **AzTax** isomer-dependent cellular bioactivity, we first assayed the potency of **AzTax3MP** for tubulin hyperpolymerisation in cell-free assays using purified tubulin. The majority-*Z* 360 nm-lit state gave a ca. 60% enhancement of polymerisation over control (benchmarked to docetaxel at 100%), while all-*E*-**AzTax3MP** gave only ca. 30% polymerisation enhancement (Fig 2d). This clarifies the mechanism of action of **AzTaxs** as MT stabilisers, like their parent taxanes.

We next studied the direct effects of *in situ-*photoisomerised **AzTax** upon cellular MT structure and MT-dependent processes. Immunofluorescence imaging *in cellulo* revealed that **AzTax3MP** causes light-dependent disruption of the MT network, resulting in mitotically-arrested cells, mitotic spindle defects, and multinucleated cells as its concentration increases (Fig 3a). The best window for visualising this isomer-dependent bioactivity lay around 0.3-1 µM (Fig S5). Z-stack projections additionally revealed cells which were substantially accumulated into rounded, mitotically arrested states that are not well resolved in single-plane imaging (Fig S5). Both the mitotic arrests, and the nuclear defects of cells that escape arrest, are hallmarks of MT stabiliser treatment^37^, arguing that the isomer-dependent cytotoxicity of **AzTax3MP** *in cellulo* arises from MT stabilisation preferentially by its *Z*-isomer.

**Figure 3.**
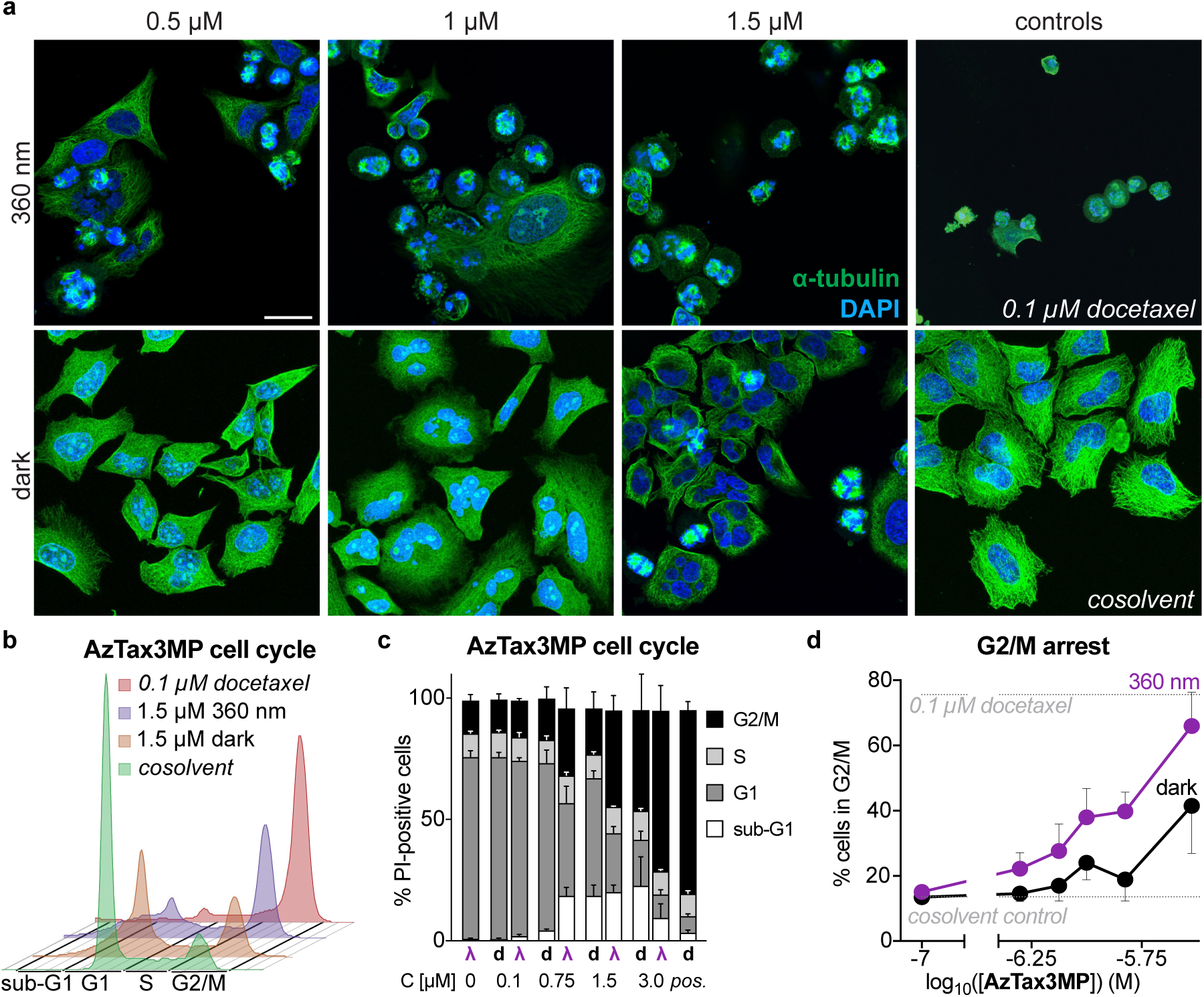
**AzTax3MP** disrupts MT structure and MT-dependent functions *in cellulo*. **(a)** Immunofluorescence staining for MT structure shows dose- and light-specific MT disruption (see also Fig S5). HeLa cells incubated for 20 h; α-tubulin in green, DNA in blue, scale bars 20 µm. **(b-d)** Flow cytometry analysis of cell cycle repartition shows that **AzTax3MP** dose-dependently gives G_2_/M (see also Fig S4) similar to that reached with positive control docetaxel (0.1 µM).

The multinucleated cells indicated that **AzTax** also inhibits MT-dependent functions such as successful completion of mitosis. To quantify this we examined cell cycle repartition after **AzTax** treatment by flow cytometry, expecting to observe G_2_/M-phase-cell cycle arrest.^38^ G_2_/M-arrest was observed with approximate EC_50_ around 1.5 µM for the “lit” **AzTax3MP**, twice as potent as *E*-**AzTax** (Fig 3b-d), mimicking the effect of docetaxel although with lower potency (Fig S4). As a control for illumination/photoswitch-dependent off-target effects, we examined the non-photoswitchably-bioactive but potent **AzTax4DMA**, which reproduced the effects of docetaxel independent of illumination conditions indicating no significant assay complications (Fig S4).

This further supported that **AzTax3MP** acts across a range of assays and readouts as a light-modulated taxane, with reproducible photocontrol over the isomers’ bioactivity - both directly against MTs, and indirectly against MT-dependent processes.

## Discussion

Photocontrol over protein function represents an attractive method to study anisotropic, multifunctional cellular systems, as it offers to address complex biology with the spatiotemporal specificity required to focus on specific roles or aspects of the system. Small molecule photopharmaceuticals have already proven valuable for their unique ability to address such targets that are not directly accessible to optogenetics, such as the MT cytoskeleton, for which a range of photoswitchable depolymerising agents have recently been reported^9,20,21^. Here we have expanded the scope of photopharmaceutical cytoskeleton reagents to demonstrate the first photoswitchable MT hyperpolymerising agents. Through early structure-photochemistry/activity-relationship studies, we have identified a lead compound **AzTax3MP** that gives robust, *in situ*-photoswitchable MT stabilising activity in cell-free and cellular assays, and can light-dependently reproduce key direct as well as downstream biological effects of the taxanes. This is a promising starting point for further reagent optimisation, and we believe that **AzTax3MP** itself will already find a range of applications particularly in embryology, neuroscience and motility, where its spatiotemporally-specific bioactivity will enable studies not previously possible.

Determining the sources of the differential bioactivity between **AzTax** isomers *in cellulo* is key for optimisation. Since modifying polarity at a distal site that should not clash sterically with the protein gave a 40-fold change of apparent potency (**AzTax3DEA** compared to **AzTax3DMA**), the sterics of the **AzTax** isomers are not necessarily the sole determinant of cellular bioactivity. Yet, polarity-dependent cellular biolocalisation or penetration cannot entirely explain the photoswitchable activity of **AzTax3MP** since it shows isomer-dependent activity in cell-free assays also, so the azobenzene must significantly impact protein-ligand affinity. We believe that maximising the bioactivity difference between isomers will require photoswitches with isomer-dependency both of sterics and of polarity. The *E*→*Z* photoswitchability of the azobenzene was not correlated to the photoswitchability of biological activity (c.f. **AzTax3MTM, AzTax4MP**). As the *Z-* **AzTax** were typically the more-active isomers, further defavouring the binding of the *E-*isomer while allowing the *Z*-isomer to retain bioactivity is likely the best way to maximise the photocontrol over inhibition. Research in these directions is underway.

Photopharmacology often assumes increasing the dynamic range of isomeric photoswitchability under a freely available choice of illumination conditions, and redshifting overall absorption wavelengths, are required for improved biological performance. In the case of **AzTax** stabilisers, probably neither consideration applies. Once MT stabilisation and hyperpolymerisation is induced by *E*→*Z* isomerisation, the altered MT biology probably cannot be instantaneously returned to its usual state even if the stabiliser would be totally removed (e.g. by complete optical, or thermal, relaxation to an inactive state). Stabilised MTs that have hyperpolymerised, potentially with loss of directionality, presumably require time to break down and return tubulin monomer to the cytoplasmic pool, so that ordinary MT structures can be rebuilt. Therefore there is probably a limit to the temporal resolution of true biological reversibility that any photoswitchable stabiliser can display, even if selected readouts (such as speed of polymerisation of individual MTs) can recover more quickly. With this consideration in mind we do not believe that improving the completeness of bidirectional isomeric photoswitchability^39^ will be as important for **AzTax** development as for other classes of inhibitors that can feature instantaneous biological response. Redshifting photoswitch absorption wavelengths is also likely to be counterproductive for a microscopy reagent, since there are few fluorescent proteins with significant excitation efficiency at laser lines above 561 nm (typically the next wavelength available is 647 nm); maintaining orthogonality to the widest possible range of imaging wavelegnths by blueshifting is probably more advantageous.^20^ However, a key property that should be readily tunable to the advantage of this system is the *E*→*Z* photoisomerisation efficiency at 405 nm, which is usually the only microscopy laser available in the 350-440 nm range. Here we consider that improved performance for **AzTax**-like reagents will depend on optimising photoconversion at the wavelength/s that will in practice be used for their photocontrol. Developing a set of standard photoswitches with better 405 nm *E*→*Z* photoconversion than these *para-*alkoxyazobenzenes (∼ 46% *Z*) yet with similar polarity and substantial stability against thermal relaxation, is a nontrivial goal of our ongoing research.

## Outlook

The **AzTax** photoswitchable microtubule stabilisers can be used in conjuction with long-term, *in situ* photoswitching in live cells to control fundamental biological processes from cytoskeleton architecture to cell survival. By complementing the existing MT-depolymerising photopharmaceuticals, **AzTaxs** now bring both principal modes of MT regulation under optical control. Through structure-photochemistry/activity-relationship studies we have identified perspectives for improving their biological photocontrol. This opens up several avenues for applying fundamental research in the rapidly evolving field of chemical photoswitches to generate specialty MT stabiliser photopharmaceuticals for cell-free mechanistic studies, cell biology, and towards *in vivo* use. We consider, more broadly, that this work will also encourage further photopharmaceutical work on other proteins inaccessible to direct optogenetics, such as the actin cytoskeleton.

**AzTax** reagents offer to aid in studying MT biology particularly where the temporally- or cell-type-specific biological roles of MT stabilisation are unclear, thus addressing a range of biological questions across the fields of neuroscience, development, cell division, signaling and migration.^25,30^ Spatially-selective induction of MT hyperpolymerisation and structural stabilization will also be of particular interest for studies where localized maintenance or growth of MT networks is thought to drive biology. Such high-spatiotemporal-precision reagents also offer an intriguing method to study the temporal and spatial dependency of biological action of their “parent” taxanes. To a large degree it is still^37^ unclear how taxanes exert their cellular/tissue-level effects *in vivo*. Yet there is enormous clinically-driven interest in increasing the understanding of taxane pharmacology, both towards improved taxol-site antimitotic therapeutics, and increasingly towards designing better combination treatment regimes involving these broad-spectrum cancer chemotherapeutics. Thus we believe the **AzTaxs** will open up a multitude of possibilities for high-precision studies not possible with previous methods, across basic and applied research.

In conclusion, we believe that this first demonstration of photoswitchable MT stabilisers represents an important step towards high-spatiotemporal-precision studies of critical proteins and processes in cell biology, and that the **AzTax** will be especially useful in studies of intracellular transport, signaling, cell motility, and neurodegeneration.

## Methods in brief

### Full and detailed experimental protocols can be found in the Supporting Information

#### Compound synthesis and characterisation

Reactions and characterisations were performed by default with non-degassed solvents and reagents (Sigma-Aldrich, TCI Europe, Fisher Scientific), used as obtained, under closed air atmosphere without special precautions. Manual flash column chromatography was performed on Merck silica gel Si-60 (40–63 µm). MPLC flash column chromatography was performed on a Biotage Isolera Spektra, using Biotage prepacked silica cartridges. Thin-layer chromatography (TLC) was run on 0.25 mm Merck silica gel plates (60, F-254), with UV light (254 nm and 365 nm) for visualization. NMR characterisation was performed on Bruker 400 or 500 MHz spectrometers. HRMS was performed by electron impact (EI) at 70 eV with Thermo Finnigan MAT 95 or Jeol GCmate II spectrometers; or by electrospray ionization (ESI) with a Thermo Finnigan LTQ FT Ultra Fourier Transform Ion Cyclotron resonance spectrometer. Analytical HPLC-MS was performed on an Agilent 1100 SL HPLC with H_2_O:MeCN eluent gradients, a Thermo Scientific Hypersil GOLD™ C18 column (1.9 µm; 3 × 50 mm) maintained at 25°C, detected on an Agilent 1100 series diode array detector and a Bruker Daltonics HCT-Ultra mass spectrometer.

#### Photocharacterisation

UV-Vis-based studies (determination of absorption spectra, photostationary states, reversibility of photoisomerisation, and *Z* to *E* relaxation) were performed on a Varian CaryScan 60 (1 cm pathlength) at room temperature with model photoswitches that were water-soluble analogues of the **AzTax** species, since reliable UV-Vis studies require compound concentrations around 25-50 µM, while the **AzTax** compounds are only reliably molecularly soluble at such concentrations with high cosolvent percentages (eg. 50% DMSO) that do not reflect the intracellular environment and also alter the isomers’ spectra, quantum yields, and relaxation times. We synthesised and used di(2-ethanol)amine carboxamides as water-soluble analogues of the taxane carboxamide **AzTaxs** (see Supporting Information) enabling measurements in PBS at pH ∼ 7.4 with only 1% of DMSO as cosolvent, thus matching the intracellular environment around the **AzTaxs**’ protein target, tubulin. “Star” 3W LEDs (360–530 nm, each FWHM ∼ 25 nm, Roithner Lasertechnik) were used for photoisomerisations in cuvette that were thus predictive of what would be obtained in the cytosol during LED-illuminated cell culture. Spectra of pure *E* and *Z* isomers were acquired from the HPLC’s inline Agilent 1100 series diode array detector (DAD) over the range 200–550 nm, manually baselining across each elution peak of interest to correct for eluent composition.

#### Tubulin Polymerisation in vitro

99% purity tubulin from porcine brain was obtained from Cytoskeleton Inc. (cat. #T240) and polymerisation assays run according to manufacturer’s instructions. Tubulin was incubated at 37 °C with “lit”- or “dark”-**AzTax** (10 µM) in buffer (with 3% DMSO, 10% glycerol) and GTP (1 mM), and the change in absorbance at 340 nm was monitored over 15 mins at 37°C^40^.

#### Cell Culture

HeLa cells were maintained under standard cell culture conditions in Dulbecco’s modified Eagle’s medium supplemented with 10% fetal calf serum (FCS), 100 U/mL penicillin and 100 U/mL streptomycin, at 37°C in a 5% CO_2_ atmosphere. Cells were transferred to phenol red free medium prior to assays. Compounds (in the all-*E* state) and cosolvent (DMSO; 1% final concentration) were added *via* a D300e digital dispenser (Tecan). Treated cells were then incubated under “dark” (light-excluded) or “lit” conditions (where pulsed illuminations were applied by multi-LED arrays to create and maintain the wave-length-dependent photostationary state isomer ratios throughout the experiment, as previously described^9^). “Lit” timing conditions were 75 ms pulses applied every 15 s.

#### Resazurin Antiproliferation Assay

As a proxy readout for viable cells, mitochondrial diaphorase activity in HeLa cell line was quantified by measuring the reduction of resazurin (7-hydroxy-3*H*-phenoxazin-3-one 10-oxide) to resorufin. 5,000 cells/well were seeded on 96-well plates. After 24 h, cells were treated with *E-***AzTaxs**, shielded from ambient light with light-proof boxes, and exposed to the appropriate light regimes. Following 48 h of treatment, cells were incubated with 20 µL of 0.15 mg/mL resazurin per well for 3 hours at 37°C. The resorufin fluorescence (excitation 544 nm, emission 590 nm) was measured using a FLUOstar Omega microplate reader (BMG Labtech). Results are represented as percent of DMSO-treated control (reading zero was assumed to correspond to zero viable cells) and represented as mean of at least three independent experiments with SD.

#### Cell Cycle analysis

*E*-**AzTaxs** were added to HeLa cells in 6-well plates (seeding density: 300,000 cells/well) and incubated under “dark” or “lit” conditions for 24 h. Cells were harvested and fixed in ice-cold 70% ethanol then stained with propidium iodide (“PI”, 200 µg/mL in 0.1 % Triton X-100 containing 200µg/mL DNase-free RNase (Thermo Fischer Scientific EN0531) for 30 min at 37°C. Following PI staining, cells were analyzed by flow cytometry using an LSR Fortessa (Becton Dickinson) run by BD FACSDiva 8.0.1 software. The cell cycle analysis was subsequently performed using FlowJo-V10 software (Tree Star Inc.). Cells were sorted into sub-G1, G1, S and G_2_/M phase according to DNA content (PI signal). Quantification from gating on the respective histograms is shown as percent of live/singlet/PI-positive parent population per cell cycle phase across different concentrations of the compound. Every experiment was performed in technical triplicates, at least three times independently, with a minimum of 10,000 (mean: 14,000) PI-positive singlet cells analyzed per replicate.

#### Immunofluorescence Staining

HeLa cells seeded on glass coverslips in 24-well plates (50,000 cells/well) were left to adhere for 24 h then treated for 24 h with **AzTaxs** under “dark” or “lit” conditions. Cover slips were washed then fixed with 0.5% glutaraldehyde, quenched with 0.1% NaBH_4_, blocked with PBS + 10% FCS, treated with rabbit alpha-tubulin primary antibody, washed, and incubated with goat-antirabbit Alexa fluor 488 secondary antibody. After washing with PBS, coverslips were mounted onto glass slides using Roti-Mount FluorCare DAPI (Roth) and imaged with a Leica SP8 confocal microscope with a 63x glycerol objective (DAPI: 405 nm, tubulin: 488 nm). Z-stacks (step size: 0.33 µm) were projected using Fiji and gamma values were adjusted for visualization.

## Acknowledgements

This research was supported by funds from the German Research Foundation (DFG: SFB1032 Nanoagents for Spatiotemporal Control project B09 to D.T. and O.T.-S.; SFB TRR 152 project P24 number 239283807, Emmy Noether grant TH2231/1-1, and SPP 1926 project number 426018126 to O.T.-S.) and the National Institutes of Health (Grant R01GM126228 to D.T.). We thank F. Ermer and M. Borowiak (LMU) for initial MTT viability assays, P.A.S (LMU) for initial synthesis, H. Harz for microscopy access (LMU microscopy platform CALM).

## Author Contributions

A.M.-D. performed synthesis, photocharacterisation, and coordinated data assembly. K.L. performed cell biology. Y.K. performed initial cell biology. C.H. performed flow cytometry. R.B. performed in vitro tubulin polymerisation assays. J.A. supervised cell biology and coordinated data assembly. D.T. designed the concept and supervised synthesis. O.T.-S. designed the study, performed and supervised synthesis, supervised all other experiments, coordinated data assembly and wrote the manuscript with input from all authors.

## Additional Information

Supporting Information (PDF) accompanies this paper: (i) synthetic protocols; (ii) photocharacterisation; (iii) biochemistry; (iv) cell biology; (v) NMR spectra.

## Dedication

*We dedicate this paper to Tim Mitchison, whose passion for microtubule biology, support of the field, and insistence on the role of small molecule reagents within it, gave much inspiration to this work.*

