## Supporting Information for "Photoswitchable microtubule stabilisers optically control tubulin cytoskeleton structure and function"

ORCIDs: J.A. 0000-0002-4879-4159; D.T. 0000-0002-6782-6056;
O.T.-S. 0000-0003-3981-651X

### Part A: Chemistry

#### *Conventions*

Abbreviations:

The following abbreviations are used: Boc – *tert*-butoxycarbonyl; brsm – based on recovered starting material; DCM: dichloromethane; DIPEA: diisopropylethylamine; DMF: dimethylformamide; DMSO: dimethylsulfoxide; EA: ethyl acetate; EDCI: 1-Ethyl-3-(3-dimethylaminopropyl)carbodiimide; Hex: distilled isohexane; HOBt: 1-hydroxybenzotriazole; Me: methyl; TFA: trifluoroacetic acid; PBS – phosphate buffered saline; T3P: propylphosphonic anhydride; wt% - percentage by weight.

Safety Hazards:

No remarkable safety hazards were encountered.

Reagents and Conditions:

Unless stated otherwise, (1) all reactions and characterisations were performed with unpurified, undried, non-degassed solvents and reagents, used as obtained, under closed air atmosphere without special precautions; (2) "hexane" used for chromatography was distilled from commercial crude isohexane fraction by rotary evaporation; (3) "column" and "chromatography" refer to manual flash column chromatography on Merck silica gel Si-60 (40–63 µm) unless otherwise specified; (4) procedures and yields are unoptimised; (5) yields refer to isolated chromatographically and spectroscopically pure materials; (6) all eluent and solvent mixtures are given as volume ratios unless otherwise specified, thus "1:1 Hex:EA" indicates a 1:1 mixture (by volume) of hexanes and ethyl acetate; (7) chromatography eluents e.g. "3:1 → 1:1" indicate a stepwise or continual gradient of eluent composition.

Thin-layer chromatography (TLC):

TLC was run on 0.25 mm Merck silica gel plates (60, F-254), typically with Hex:EA eluents. All compounds carrying an azobenzene need no visualization on the TLC-plate but are clearly visible as colored spots (color range yellow to red). The presence of an azobenzene can be verified by exposure of the colored spot to TFA vapors, which transiently changes the color to shades of purple according to basicity. For further visualization UV light (254 nm) was used. TLC characterizations are abbreviated as $\mathbf{R}_f$ = 0.64 (UV 254 nm, Hex:EA = 1:1).

Nuclear magnetic resonance spectroscopy (NMR):

Standard NMR characterisation was by $^{1}$H- and $^{13}$C-NMR spectra on an Avance III HD 400 MHz Bruker BioSpin or Bruker Ascend 400, or Avance III HD 500 MHz Bruker BioSpin (1H: 400 MHz and 500 MHz, 13C: 101 MHz and 126 MHz). Chemical shifts (δ) are reported in ppm calibrated to residual non-perdeuterated solvent as an internal reference[1]. Peak descriptions singlet (s), doublet (d), triplet (t), quartet (q), multiplet (m) and broad (br) are used. Apparent multiplicities (resolved by 2D experiments or determined by complete spectral assignment) are denoted by a tilde, eg. “appears as a triplet with apparent coupling constant *J* = 3 Hz” is denoted (~t, 3 Hz). NMR spectra are given in Part F.

High resolution mass spectrometry (HRMS):

HRMS was performed by electron impact (EI) at 70 eV with a Thermo Finnigan MAT 95 or a Jeol GCmate II spectrometer; or electrospray ionization (ESI) with a Thermo Finnigan LTQ FT Ultra Fourier Transform Ion Cyclotron resonance mass spectrometer; as specified.

High-performance liquid chromatography coupled to mass spectrometry (LCMS):

Analytical high-performance liquid chromatography (HPLC) was performed on an Agilent 1100 SL coupled HPLC system with (a) a binary pump to deliver $H_2O$:MeCN eluent mixtures containing 0.1% formic acid at a 0.4 mL/min flow rate, (b) Thermo Scientific Hypersil GOLD™ C18 column (1.9 µm; 3 × 50 mm) maintained at 25 °C, whereby the solvent front eluted at $t_{ret}$ = 0.5 min, (c) an Agilent 1100 series diode array detector, (d) a Bruker HCT Ultra mass spectrometer. Typical run conditions were a linear gradient of $H_2O$:MeCN eluent composition from 90:10 through to 1:99, applied during the separation phase (first 5 min), then 0:100 for 2 min for flushing; the column was (re)equilibrated with 90:10 eluent mixture for 2 min before each run. Ion peaks from (positive/negative mode) are reported as (+/-) with units Th (m/z). Thus “LCMS(+): $t_{ret}$ = 5.60 & 5.82 min, each 419 Th = $[MH]^+$” indicates LCMS under the standard run conditions with ESI ionisation giving two positive ion peaks eluting at 5.60 and 5.82 min retention times, each at m/z = 419 Th, attributed as the protonated molecular ion. Unless stated otherwise, all reported peaks in the positive mode were $[MH]^+$ peaks.

#### *Standard Procedures*

Where Standard Procedures were used in synthesis, unless stated otherwise, the amounts of reactants/reagents employed were implicitly adjusted to maintain the same molar ratios as in the given Procedure, and no other alterations from the Standard Procedure (eg. reaction time, extraction solvent, temperature) were made, unless stated otherwise.

Standard Procedure A: Azo coupling with a phenol partner

A flask was charged with the aniline coupling partner (1.0 eq) and MeOH (3 mL/mmol). Aqueous HCl (2 M, 6.0 eq) was added. The reaction mixture was cooled to 0 °C and a 2 M aqueous of solution of $NaNO_2$ (1.1 eq) was added dropwise. It was allowed to stir for 30 min. A solution of the phenol coupling partner (1.1 eq) in MeOH (4 mL/mmol) and 0.5 M aqueous $HK_2PO_4$ (4 mL/mmol) was prepared at 0 °C. The diazonium solution was added dropwise onto the phenol mixture. The pH was maintained between 9-10 by adding aq. KOH (1 M). Upon completion of the addition the reaction mixture was allowed to stir for 1 h in the cold. The reaction progress was monitored by LCMS/TLC analysis. The reaction was quenched by the adjustment of the pH to pH 4-6 with 2 M aqueous HCl and extracted with EA (3 × 20 mL/mmol). The combined organic phases were dried with $Na_2SO_4$, filtrated and concentrated. The crude product was purified by flash chromatography using a Hex:EA gradient.

Standard Procedure B: Azo coupling with a dialkylaniline partner

A flask was charged with the aniline coupling partner (1.0 eq) and MeOH (3 mL/mmol). Aqueous HCl (2 M, 6.0 eq) was added. The reaction mixture was cooled to 0 °C and a 2 M aqueous of solution of $NaNO_2$ (1.1 eq) was added dropwise. It was allowed to stir for 30 min. Acetic acid (3 mL/mmol) followed by the dialkylaniline coupling partner (1.5 eq) were added neat. Sodium acetate (10.0 eq) was added portionwise. Upon completion of the addition the reaction mixture was allowed to stir for 1 h in the cold. The reaction progress was monitored by LCMS/TLC analysis. The reaction was quenched by neutralization with KOH solution (1 M) and extracted with EA (3 × 20 mL/mmol). The combined organic phases were dried with $Na_2SO_4$, filtrated and concentrated. The crude product was purified by flash chromatography using a Hex:EA gradient.

Standard Procedure C: Methylation of a *para*-hydroxy azobenzene

A flask was charged with the respective azobenzene compound and acetone (5 mL/mmol) was added. Potassium carbonate (5.0 eq) was added. Iodomethane (3.00 eq) was added dropwise. The reaction mixture was heated to 50 °C for typically 5 h. The reaction progress was monitored by LCMS/TLC analysis. The reaction was quenched by the addition of water (20 mL/mmol) and extracted with EA (3 × 20 mL/mmol). The combined organic phases were dried with $Na_2SO_4$, filtrated and concentrated. The crude product was purified by flash chromatography on silica using a Hex:EA gradient.

Standard Procedure D: Hydrolysis of an azobenzenecarboxylate ester

A flask was charged with the respective azobenzenecarboxylate ester and MeOH (5 mL/mmol) was added. Potassium hydroxide (5.0 eq) was added neat. The reaction mixture was heated to 65 °C for 12 h. The reaction progress was monitored by LCMS/TLC analysis. Upon completion the reaction was quenched with water (20 mL/mmol), neutralized with 2 M aqueous KOH and extracted with EA (3 × 20 mL/mmol). The combined organic phases were dried with $Na_2SO_4$, filtrated and concentrated. Typically, no further purification was needed.

Standard Procedure E: Docetaxel deprotection and amide coupling

A flask was charged with docetaxel (16 mg, 20 $\mu$mol, 1.0 eq) and DCM (2 mL) and the solution stirred at 0 °C for 2 min. TFA (2 mL) was added and the mixture stirred at 0 °C for 1 hour. The solution was added into rapidly stirred sat. aq. $NaHCO_3$ (15 mL). Solid $NaHCO_3$ was added until all TFA was neutralized. The mixture was extracted with DCM (3 × 10 mL). The combined organic layers were washed with sat. aq. $NaHCO_3$ (10 mL), brine (10 mL), dried on $Na_2SO_4$, filtered and concentrated to a colourless crude foam (typically 10 mg, 15 $\mu$mol, 71%; LCMS(+): $t_{ret}$ = 4.71 min, 708 Th = $[MH]^+$). The crude was dissolved in HPLC-grade DMF (2 mL). The azobenzene carboxylic acid (1.2 eq.) was dissolved in HPLC-grade DMF (1 mL), HOBt·$H_2O$ (2.5 eq) and EDCI (2.25 eq) were added and the solution stirred at room temperature for 5 min. A DMF (1 mL) solution of DIPEA (4.0 eq) was added dropwise and stirring continued for 10 min. The solution of crude deprotected docetaxel was added and the solution stirred for

12 h at room temperature, then poured into 10% aq. $NaHCO_3$ (20 mL) and extracted with DCM (3 × 10 mL). The combined organic layers were washed with sat. aq. $NaHCO_3$ (10 mL), sat. aq. LiCl (10 mL), brine (10 mL), dried on $Na_2SO_4$, filtered and concentrated to a yellow solid. Chromatography on silica with a iHex:EA= 7:3→1:1 then DCM:MeOH= 1:0→9:1 gradient couple typically separated the product fractions. These were combined, concentrated, and dried under high vacuum.

Standard Procedure F: Preparation of soluable azo derivatives for photocharacterization

A flask was charged with the azobenzene carboxylic acid (1.0 eq) and DMF (25 mL/mmol) was added. Triethylamine (10.0 eq) was added. Diethanolamine (2.0 eq) dissolved in DMF (0.65 mL/mmol) was added to the reaction mixture. T3P (2.0 eq, ≥50 wt. % in EA) was added and the resulting solution was allowed to stir for 16 h at 25 °C. Progress of the reaction was monitored by LCMS. Upon completion the DMF was removed *in vacuo* and the crude product was purified by flash chromatography on silica using a DCM:MeOH gradient.

#### *Azobenzene carboxylic acids*

**4-(phenyldiazenyl)benzoic acid (4H-$CO_2$H)**

Commercially available (CAS 1562-93-2).

**4-((4-(dimethylamino)phenyl)diazenyl)benzoic acid (4DMA-$CO_2$H)**

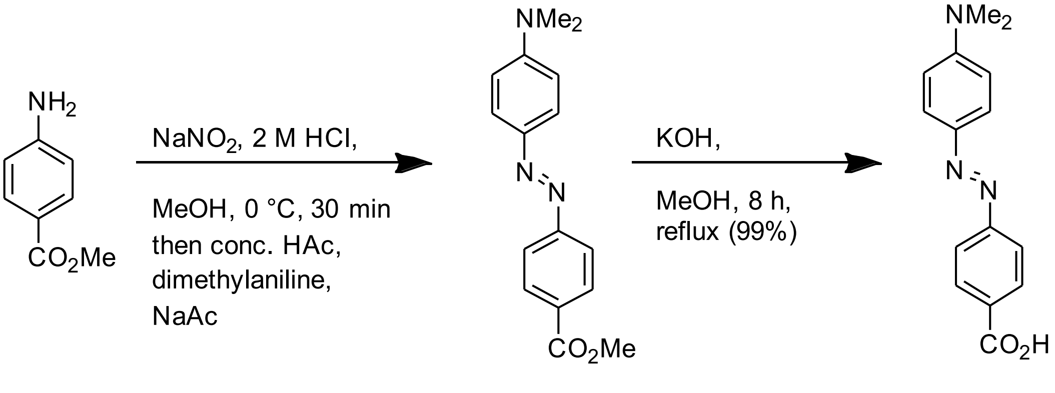

**S1** **4DMA-$CO_2$H**

By standard procedure B, commercially available methyl 4-aminobenzoate (302 mg, 2.0 mmol, 1.0 eq) was reacted with dimethyaniline (364 mg, 3.0 mmol, 1.5 eq). After purification by flash chromatography (EA:iHex, 9:1 → 8:2) the desired product **methyl 4-((4-(dimethylamino)phenyl)diazenyl)benzoate** (**S1**) (224 mg, 0.80 mmol, 40%) was obtained as red solid. Spectral data matches literature[2]: **$^1$H NMR** (400 MHz, chloroform-*d*) δ (ppm) = 8.19 – 8.10 (m, 2H), 7.96 – 7.82 (m, 4H), 6.79 – 6.73 (m, 2H), 3.94 (s, 3H), 3.11 (s, 6H). **$^{13}$C NMR** (101 MHz, chloroform-*d*) δ (ppm) = 167.1, 156.3, 153.2, 144.0, 130.8, 130.4, 125.8, 122.3, 111.7, 52.5, 40.6. **LCMS(+)**: $t_{ret}$ = 4.7 min, 284 Th = $[MH]^+$. **HRMS (EI):** calc. for $[C_{16}H_{17}O_2N_3]^+$ = $[M]^+$: 283.1321; found: 283.1314.

By standard procedure D, **S1** (100 mg, 0.35 mmol, 1.0 eq) was reacted to the desired product **4DMA-$CO_2$H** (93 mg, 0.40 mmol, 98%) which was obtained as a red solid. Spectral data matches literature[2]: **$^1$H NMR** (400 MHz, DMSO-d6) δ (ppm) = 8.13 – 8.02 (m, 2H), 7.89 – 7.78 (m, 4H), 6.92 – 6.79 (m, 2H), 3.08 (s, 6H). **$^{13}$C NMR** (101 MHz, DMSO) δ (ppm) = 166.9, 155.1, 153.0, 142.7, 130.9, 130.5, 125.3, 121.7, 111.6, 39.8. **LCMS(+):** $t_{ret}$ = 4.7 min, 270 Th = [MH]$^+$. **HRMS (EI):** calc. for $[C_{15}H_{15}O_2N_3]^+$ = [M]$^+$: 269.1164; found: 269.1158.

**4-((4-methoxyphenyl)diazenyl)benzoic acid (4MP-CO$_2$H)**

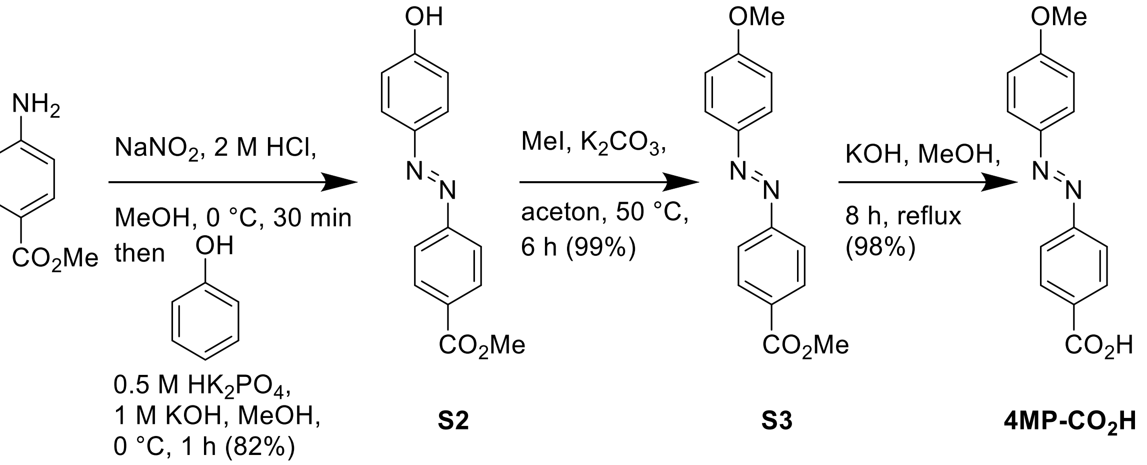

By standard procedure A, commercially available methyl 4-amino benzoate (302 mg, 2.0 mmol, 1.0 eq) was reacted with phenol (207 mg, 2.2 mmol, 1.1 eq). After purification by means of flash chromatography (EA:iHex, 8:2 → 1:1) the desired product **methyl 4-((4-hydroxyphenyl)diazenyl)benzoate** (**S2**) (421 mg, 1.6 mmol, 82%) was obtained as an orange solid. Spectral data matches literature[3]: **$^1$H NMR** (400 MHz, DMSO-*d*$_6$) δ (ppm) = 8.17 – 8.08 (m, 2H), 7.93 – 7.88 (m, 2H), 7.88 – 7.80 (m, 2H), 7.00 – 6.93 (m, 2H), 3.89 (s, 3H). **$^{13}$C NMR** (101 MHz, DMSO-*d*$_6$) δ (ppm) = 165.7, 161.8, 154.8, 145.3, 130.6, 130.5, 125.4, 122.3, 116.1, 52.4. **LCMS(+):** $t_{ret}$ = 5.0 min, 257 Th = [MH]$^+$. **HRMS (EI):** calc. for $[C_{16}H_{17}O_2N_3]^+$ = [M]$^+$: 256.0848; found: 256.0843.

By standard procedure C, **S2** (410 mg, 1.6 mmol, 1.0 eq) was reacted with methyl iodide (681 mg, 4.8 mmol, 3.0 eq). After purification by means of flash chromatography (EA:iHex, 9:1 → 7:3) the desired product **methyl 4-((4-methoxyphenyl)diazenyl)benzoate** (**S3**) (431 mg, 1.6 mmol, 99%) was obtained as an orange solid. Spectral data matches literature[4]: **$^1$H NMR** (400 MHz, chloroform-*d*) δ (ppm) = 8.21 – 8.14 (m, 2H), 7.95 (d, *J* = 9.0 Hz, 2H), 7.91 (d, *J* = 8.7 Hz, 2H), 7.03 (d, *J* = 9.0 Hz, 2H), 3.95 (s, 3H), 3.91 (s, 3H). **$^{13}$C NMR** (101 MHz, chloroform-*d*) δ (ppm) = 166.8, 162.8, 155.5, 147.2, 131.3, 130.7, 125.3, 122.5, 114.5, 55.8, 52.4.. **LCMS(+):** $t_{ret}$ = 6.0 min, 271 Th = [MH]$^+$. **HRMS (EI):** calc. for $[C_{15}H_{14}N_2O_3]^+$ = [M]$^+$: 270.1004; found: 270.0998.

By standard procedure D, **S3** (100 mg, 0.35 mmol, 1.0 eq) was reacted to the desired product **4-((4-(dimethylamino)phenyl)diazenyl)benzoic acid** (**4MP-CO$_2$H**) (93 mg, 0.40 mmol, 98%) was obtained as orange solid. Spectral data matches literature[5]: **$^1$H NMR** (400 MHz, DMSO-d$_6$) δ (ppm)= 8.03 (d, J = 8.4 Hz, 2H), 7.91 (d, J = 9.0 Hz, 2H), 7.79 (d, J = 8.4 Hz, 2H), 7.14 (d, J = 9.0 Hz, 2H), 3.87 (s, 3H). **$^{13}$C NMR** (101 MHz, DMSO) δ 166.8, 162.6, 154.4,

146.2, 132.2, 130.6, 125.0, 122.2, 114.8, 55.7. **LCMS(+):** $t_{ret}$ = 5.0 min, 257 Th = $[MH]^+$. **HRMS (EI):** calc. for $C_{14}H_{12}O_3N_2^+$ $[M]^+$: 256.0848; found: 256.0838.

**3-(phenyldiazenyl)benzoic acid (3H-CO₂H)**

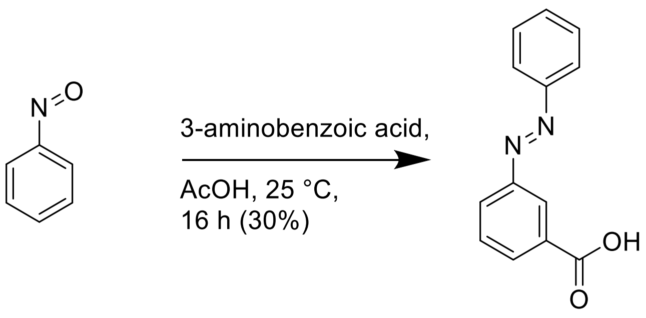

**3H-CO₂H**

Prepared according to literature[6]. A flask was charged with nitrosobenzene (214 mg, 2.0 mmol, 1.0 eq) and 5 mL AcOH was added. 3-aminobenzoic acid (274 mg, 2.0 mmol, 1.0 eq) was added and the reaction mixture was allowed to stir for 24 h at 25 °C. LCMS showed completion. The reaction was quenched with water (30 mL) and the aqueous phase was extracted with EA (3 × 50 mL). The combined organic phases were dried with $Na_2SO_4$, filtrated and concentrated. After purification by means of flash chromatography (DCM:MeOH, 98:2 → 94:6) the desired product **3-(phenyldiazenyl)benzoic acid** (**3H-CO₂H**) (134 mg, 0.59 mmol, 30%) was obtained as an orange solid. Spectral data matches literature[6]: **$^1$H NMR** (500 MHz, DMSO-$d_6$) δ (ppm) = 13.33 (s, 1H), 8.38 (t, $J$ = 1.9 Hz, 1H), 8.14 (ddt, $J$ = 13.7, 7.7, 1.4 Hz, 2H), 7.95 (dd, $J$ = 8.0, 1.8 Hz, 2H), 7.75 (t, $J$ = 7.8 Hz, 1H), 7.67 – 7.57 (m, 3H). **$^{13}$C NMR** (126 MHz, DMSO-$d_6$) δ (ppm) = 166.7, 151.9, 151.8, 132.2, 132.0, 131.8, 130.0, 129.5, 127.4, 122.7, 122.2. **LCMS(+):** $t_{ret}$ = 4.9 min, 227 Th = $[MH]^+$.

**3-((4-methoxyphenyl)diazenyl)benzoic acid (3MP-CO₂H)**

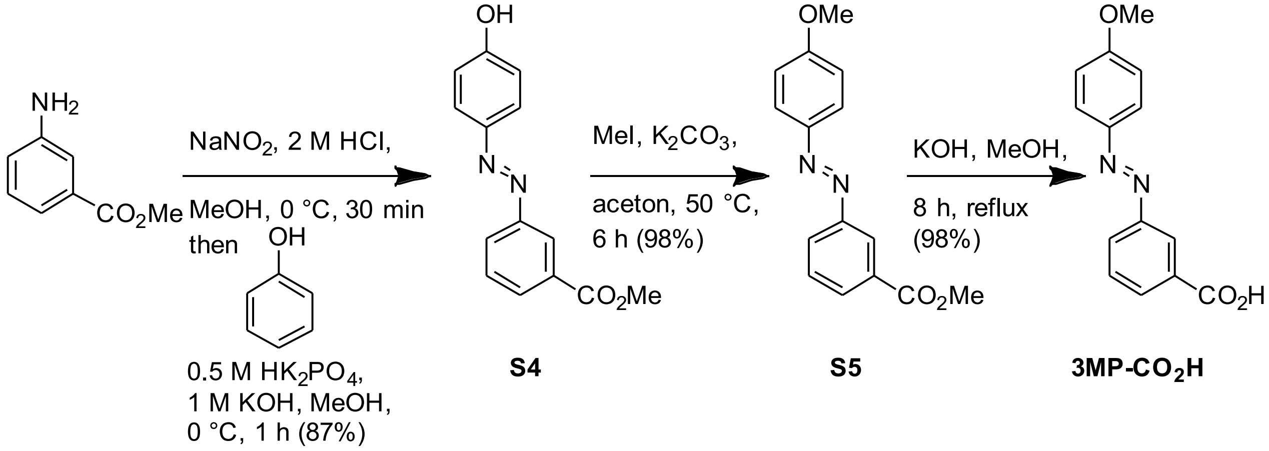

By standard procedure A, commercially available methyl 3-amino benzoate (302 mg, 2.0 mmol, 1.0 eq) was reacted with phenol (207 mg, 2.2 mmol, 1.1 eq). After purification by means of flash chromatography (EA:iHex, 9:1 → 7:3) the desired product **methyl 3-((4-hydroxyphenyl)diazenyl)benzoate** (**S4**) (445 mg, 1.6 mmol, 87%) was obtained as an orange solid. $^{1}$**H NMR** (400 MHz, chloroform-*d*) δ (ppm) = 8.53 (t, *J* = 1.9 Hz, 1H), 8.12 (dt, *J* = 7.8, 1.4 Hz, 1H), 8.07 (ddd, *J* = 8.0, 2.0, 1.1 Hz, 1H), 7.94 – 7.88 (m, 2H), 7.58 (t, *J* = 7.8 Hz, 1H), 7.01 – 6.94 (m, 2H), 3.97 (s, 3H). $^{13}$**C NMR** (101 MHz, chloroform-*d*) δ (ppm) = 166.9, 158.9, 152.8, 147.2, 131.4, 131.3, 129.3, 126.9, 125.5, 123.9, 116.0, 77.4, 52.5. **LCMS(+):** $t_{ret}$ = 5.0 min, 257 Th = $[MH]^+$, **HRMS (EI):** calc. for $[C_{14}H_{12}O_3N_2]^+$ = $[M]^+$: 256.0848; found: 256.0850.

By standard procedure C, **S4** (440 mg, 1.7 mmol, 1.0 eq) was reacted with methyl iodide (731 mg, 5.2 mmol, 3.0 eq). After purification by means of flash chromatography (EA:iHex, 9:1 → 7:3) the desired product **methyl 3-((4-methoxyphenyl)diazenyl)benzoate** (**S5**) (454 mg, 1.7 mmol, 98%) was obtained as an orange solid. $^{1}$**H NMR** (400 MHz, chloroform-*d*) δ (ppm) = 8.56 – 8.50 (m, 1H), 8.11 (ddd, *J* = 7.7, 1.7, 1.2 Hz, 1H), 8.06 (ddd, *J* = 8.0, 2.1, 1.2 Hz, 1H), 8.00 – 7.91 (m, 2H), 7.58 (td, *J* = 7.8, 0.5 Hz, 1H), 7.07 – 6.98 (m, 2H), 3.97 (s, 3H), 3.90 (s, 3H). $^{13}$**C NMR** (101 MHz, chloroform-*d*) δ (ppm) = 166.7, 162.4, 152.8, 146.9, 131.2, 131.1, 129.1, 126.8, 125.0, 123.7, 114.3, 55.6, 52.3. **LCMS(+):** $t_{ret}$ = 5.9 min, 271 Th = $[MH]^+$**. HRMS (EI):** calc. for $[C_{15}H_{14}N_2O_3]^+$ = $[M]^+$: 270.1004; found: 270.0998.

By standard procedure D, **S5** (430 mg,1.6 mmol, 1.0 eq) was reacted to the desired product **3-((4-methoxyphenyl)diazenyl)benzoic acid** (**3MP-CO$_2$H**) (399 mg, 1.6 mmol, 98%) was obtained as orange solid. $^{1}$**H NMR** (400 MHz, methanol-*d*$_4$) δ (ppm) = 10.02 (t, *J* = 1.8 Hz, 1H), 9.67 (dt, *J* = 7.7, 1.4 Hz, 1H), 9.63 (ddd, *J* = 8.0, 2.1, 1.2 Hz, 1H), 9.53 – 9.47 (m, 2H), 9.19 (t, *J* = 7.8 Hz, 1H), 8.67 – 8.62 (m, 2H), 5.46 (s, 3H). $^{13}$**C NMR** (101 MHz, methanol-*d*$_4$) δ (ppm) = 167.79, 162.82, 152.72, 146.72, 131.84, 130.89, 129.00, 126.32, 124.62, 123.06, 114.04, 54.76. **LCMS(+):** $t_{ret}$ = 5.2 min, 257 Th = $[MH]^+$**. HRMS (EI):** calc. for $[C_{14}H_{12}O_3N_2]^+$ = $[M]^+$: 256.0848; found: 256.0842.

**3-((4-(dimethylamino)phenyl)diazenyl)benzoic acid (3DMA-CO$_2$H)**

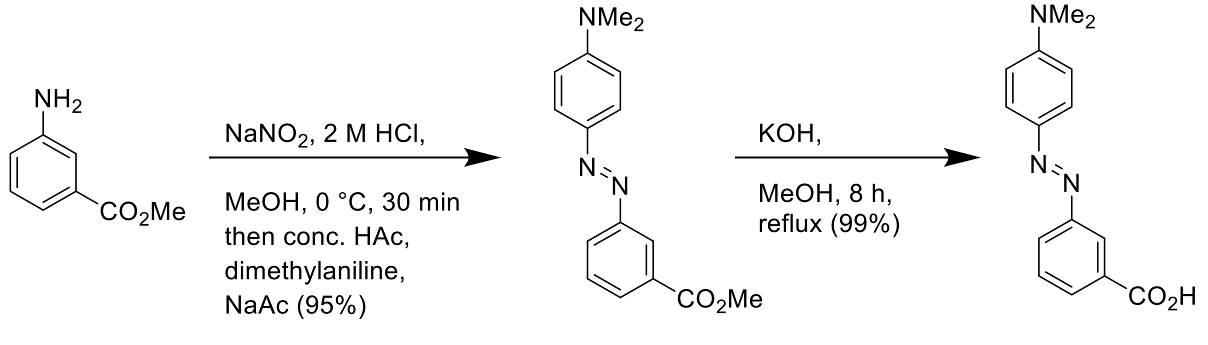

**S6** **3DMA-CO$_2$H**

By standard procedure B, commercially available methyl 3-amino benzoate (302 mg, 2.0 mmol, 1.00 eq) was reacted with dimethyaniline (364 mg, 3.0 mmol, 1.5 eq). After purification by means of flash chromatography (EA:iHex, 9:1 → 8:2) the desired product **methyl 3-((4-(dimethylamino)phenyl)diazenyl)benzoate** (**S6**) (539 mg, 1.9 mmol, 95%) was obtained as red solid. Spetral data matches literature[7]: $^{1}$**H NMR** (500 MHz, chloroform-*d*) δ (ppm) = 8.49 (t, *J* = 1.8 Hz, 1H), 8.04 (tdd, *J* = 7.3, 2.4, 1.2 Hz, 2H), 7.92 (d, *J* = 9.1 Hz, 2H), 7.54 (t, *J* = 7.8 Hz, 1H), 6.78 (d, *J* = 9.0 Hz, 2H), 3.96 (s, 3H), 3.11 (s, 6H). $^{13}$**C NMR** (126 MHz, chloroform-*d*) δ (ppm) = 166.9, 153.1, 152.7, 143.5, 131.1, 130.1, 129.0, 126.4, 125.4, 123.4, 111.7, 52.3, 40.4. **LCMS(+):** $t_{ret}$ = 6.0 min, 284 Th = $[MH]^+$. **HRMS (EI):** calc. for $[C_{16}H_{17}O_2N_3]^+$ = $[M]^+$: 283.1321; found: 283.1315.

By standard procedure D, **S6** (100 mg, 0.35 mmol, 1.0 eq) was reacted to the desired product **3-((4-(dimethylamino)phenyl)diazenyl)benzoic acid** (**3DMA-CO$_2$H**) (93 mg, 0.35 mmol, 98%) was obtained as red solid. Spectral data matches literature[8]: **$^1$H NMR** (500 MHz, methanol-*d*$_4$) δ (ppm) = 8.41 (t, *J* = 1.8 Hz, 1H), 8.04 (dt, *J* = 7.7, 1.4 Hz, 1H), 8.02 – 7.99 (m, 1H), 7.89 – 7.84 (m, 2H), 7.59 (t, *J* = 7.8 Hz, 1H), 6.87 – 6.82 (m, 2H), 3.10 (s, 6H). **$^{13}$C NMR** (126 MHz, methanol-*d*$_4$) δ (ppm) = 169.6, 154.8, 154.7, 144.8, 133.3, 131.4, 130.4, 127.5, 126.4, 124.2, 112.8, 40.5. **LCMS(+):** $t_{ret}$ = 6.2 min, 270 Th = $[MH]^+$. **HRMS (EI):** calc. for $[C_{16}H_{17}O_2N_3]^+$ = $[M]^+$: 269.1164; found: 269.1159.

**4-methoxy-3-((3,4,5-trimethoxyphenyl)diazenyl)benzoic acid (3MTM-CO$_2$H)**

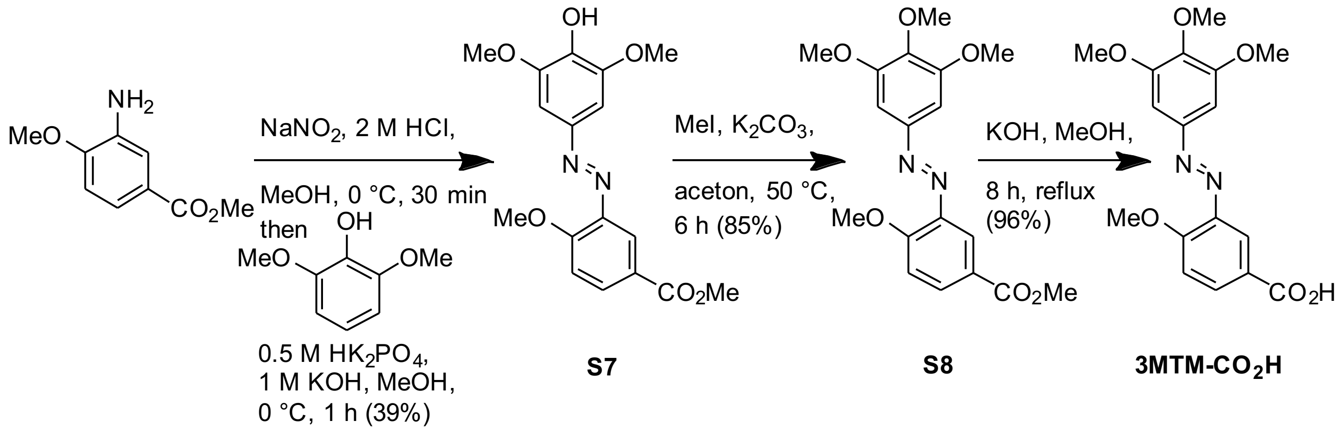

By standard procedure A, commercially available methyl 3-amino 4-methoxybenzoate (181 mg, 1.0 mmol, 1.0 eq) was reacted with 2,6-dimethoxyphenol (185 mg, 1.2 mmol, 1.2 eq). After purification by means of flash chromatography (EA:iHex, 9:1 → 1:1) the desired product **methyl 3-((4-hydroxy-3,5-dimethoxyphenyl)diazenyl)-4-methoxybenzoate** (**S7**) (136 mg, 0.39 mmol, 39%) was obtained as an yellow solid. **$^1$H NMR** (400 MHz, chloroform-*d*) δ (ppm) = 8.28 (d, J = 2.2 Hz, 1H), 8.12 (dd, J = 8.7, 2.2 Hz, 1H), 7.32 (s, 2H), 7.12 (d, J = 8.7 Hz, 1H), 5.88 (s, 1H), 4.08 (s, 3H), 4.01 (s, 6H), 3.92 (s, 3H). **$^{13}$C NMR** (101 MHz, chloroform-*d*) δ (ppm) = 166.8, 159.9, 147.4, 146.2, 142.0, 138.4, 133.2, 123.1, 119.0, 112.2, 101.1, 56.7, 56.6, 52.3. **LCMS(+):** $t_{ret}$ =4.5 min, 347 Th = $[MH]^+$. **HRMS (EI):** calc. for $[C_{17}H_{18}N_2O_6]^+$ = $[M]^+$: 346.1165; found: 346.1160.

By standard procedure C, **S7** (136 mg, 0.39 mmol, 1.0 eq) was reacted with methyl iodide (111 mg, 0.79 mmol, 2.0 eq). After purification by means of flash chromatography (EA:iHex, 9:1 → 1:1) the desired product **methyl 4-methoxy-3-((3,4,5-trimethoxyphenyl)diazenyl)benzoate** (**S8**) (120 mg, 0.33 mmol, 85%) was obtained as an orange solid. **$^1$H NMR** (400 MHz, chloroform-*d*) δ (ppm) = 8.27 (d, *J* = 2.2 Hz, 1H), 8.14 (dd, *J* = 8.7, 2.2 Hz, 1H), 7.27 (s, 2H), 7.12 (d, *J* = 8.8 Hz, 1H), 4.08 (s, 3H), 3.97 (s, 6H), 3.94 (s, 3H), 3.92 (s, 3H). **$^{13}$C NMR** (101 MHz, chloroform-*d*) δ (ppm) = 166.6, 159.9, 153.5, 148.9,

141.8, 141.0, 133.4, 122.9, 118.8, 112.1, 100.8, 61.1, 56.4, 56.3, 52.1. **LCMS(+):** $t_{ret}$ = 4.9 min, 361 Th = $[MH]^+$. **HRMS (EI):** calc. for $[C_{18}H_{20}N_2O_6]^+$ = $[M]^+$:360.1321; found: 360.1314.

By standard procedure D, **S9** (110 mg,0.32 mmol, 1.0 eq) was reacted to the desired product **4-methoxy-3-((3,4,5-trimethoxyphenyl)diazenyl)benzoic acid** (**3MTM-CO$_2$H**) (102 mg, 0.30 mmol, 96%) was obtained as yellow solid.

**$^1$H NMR** (400 MHz, DMSO-*d*$_6$) δ (ppm) = 8.11 – 8.05 (m, 2H), 7.38 (d, *J* = 9.4 Hz, 1H), 7.26 (s, 2H), 4.04 (s, 3H), 3.89 (s, 6H), 3.77 (s, 3H). **$^{13}$C NMR** (101 MHz, DMSO-*d*$_6$) δ (ppm) = 167.2, 160.1, 153.8, 148.6, 141.2, 140.9, 134.8, 134.0, 123.5, 118.1, 113.8, 101.0, 60.8, 57.0, 56.5. **LCMS(+):** $t_{ret}$ = 4.3 min, 347 Th = $[MH]^+$. **HRMS (EI):** calc. for $[C_{18}H_{20}N_2O_6]^+$ = $[M]^+$:346.1165; found: 346.1157.

**3-((3,4,5-trimethoxyphenyl)diazenyl)benzoic acid (3TM-CO$_2$H)**

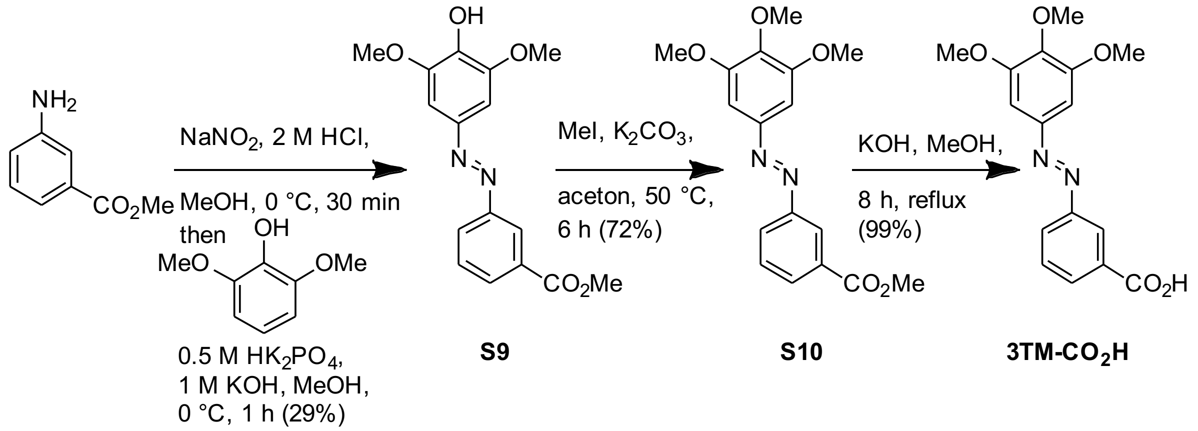

By standard procedure A, commercially available methyl 3-amino benzoate (151 mg, 1 mmol, 1.00 eq) was reacted with 2,6-dimethoxyphenol (185 mg, 1.1 mmol, 1.1 eq). After purification by means of flash chromatography (EA:iHex, 9:1 → 6:4) the desired product **methyl 3-((4-hydroxy-3,5-dimethoxyphenyl)diazenyl)benzoate** (**S9**) (229 mg, 0.72 mmol, 73%) was obtained as an orange solid.**$^1$H NMR** (400 MHz, chloroform-d) δ (ppm) = 8.46 (t, *J* = 1.7 Hz, 1H), 8.05 (ddd, *J* = 7.7, 1.7, 1.2 Hz, 1H), 7.99 (ddd, *J* = 8.0, 2.1, 1.2 Hz, 1H), 7.51 (td, *J* = 7.8, 0.5 Hz, 1H), 7.18 (s, 1H), 5.81 (s, 1H), 3.94 (s, 6H), 3.90 (s, 3H).**$^{13}$C NMR** (101 MHz, chloroform-d) δ (ppm) = 166.8, 152.8, 147.4, 138.5, 131.5, 131.3, 129.3, 126.9, 123.9, 100.9, 56.6, 52.5. **LCMS(+):** $t_{ret}$ = 4.9 min, 317 Th = $[MH]^+$. **HRMS (EI):** calc. for $[C_{16}H_{16}O_5N_2]^+$ = $[M]^+$: 316.1059; found: 316.1050.

By standard procedure C, **S9** (221 mg, 0.70 mmol, 1.0 eq) was reacted with methyl iodide (198 mg, 1.4 mmol, 2.0 eq). After purification by means of flash chromatography (EA:iHex, 9:1 → 7:3) the desired product **methyl 3-((3,4,5-trimethoxyphenyl)diazenyl)benzoate** (**S10**) (174 mg, 0.53 mmol, 72%) was obtained as a yellow solid. **$^1$H NMR** (400 MHz, chloroform-*d*) δ (ppm) = 8.55 (t, *J* = 1.6 Hz, 1H), 8.14 (ddd, *J* = 7.7, 1.7, 1.2 Hz, 1H), 8.09 (ddd, *J* = 7.9, 2.1, 1.2 Hz, 1H), 7.60 (td, *J* = 7.8, 0.5 Hz, 1H), 7.29 (s, 2H), 3.98 (s, 6H), 3.98 (s, 3H) 3.95 (s, 3H). **$^{13}$C NMR** (101 MHz, chloroform-*d*) δ (ppm) = 166.7, 153.7, 152.7, 148.5, 141.2, 131.6, 131.5, 129.4, 127.0, 124.1, 100.8, 61.2, 56.4, 52.5. **LCMS(+):** $t_{ret}$ = 5.7 min, 331 Th = $[MH]^+$. **HRMS (EI):** calc. for $[C_{17}H_{18}O_5N_2]^+$ = $[M]^+$: 330.1216; found: 330.1206.

By standard procedure D, **S10** (174 mg,0.53 mmol, 1.0 eq) was reacted to the desired product **3-((3,4,5-trimethoxyphenyl)diazenyl)benzoic acid** (**3TM-$CO_2$H**) (167 mg, 0.53 mmol, 99%) was obtained as yellow solid. **$^1$H NMR** (500 MHz, methanol-*d*$_4$) δ (ppm) = 8.53 (t, *J* = 1.8 Hz, 1H), 8.17 (dt, *J* = 7.7, 1.4 Hz, 1H), 8.13 (ddd, *J* = 7.9, 2.1, 1.2 Hz, 1H), 7.67 (t, *J* = 7.8 Hz, 1H), 7.38 (s, 2H), 3.98 (s, 6H), 3.89 (s, 3H). **$^{13}$C NMR (126 MHz,** methanol-*d*$_4$**)** δ (ppm) = 168.3, 153.6, 152.4, 148.4, 140.9, 132.8, 131.4, 129.0, 126.3, 123.2, 100.4, 59.9, 55.3, 55.2. **LCMS(+):** $t_{ret}$ = 4.6 min, 317 Th = [MH]$^+$. **HRMS (EI):** calc. for $[C_{16}H_{16}O_5N_2]^+$ = [M]$^+$: 316.1043; found: 316.1059.

**3-((4-(bis(2-hydroxyethyl)amino)phenyl)diazenyl)benzoic acid (3DEA-$CO_2$H)**

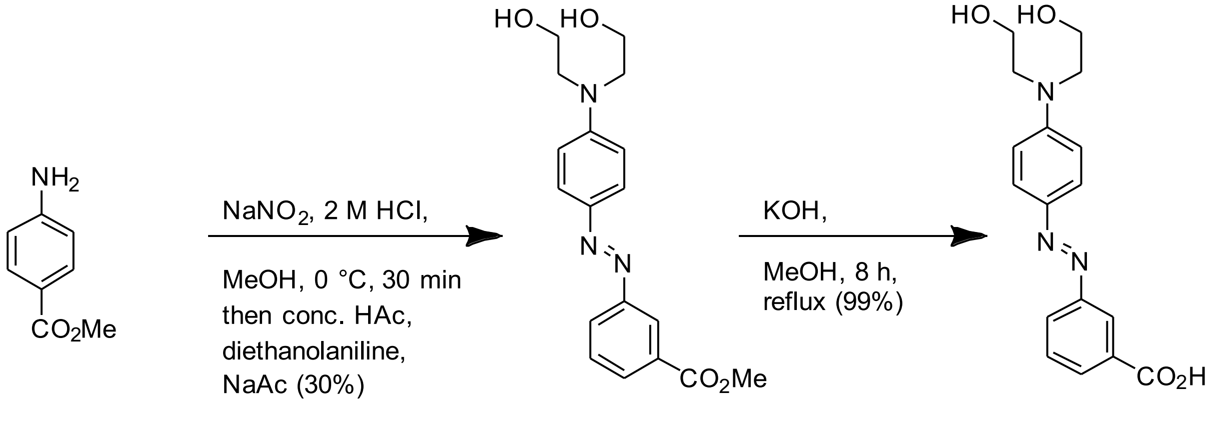

**S11** **3DEA-$CO_2$H**

By standard procedure B, commercially available methyl 3-amino benzoate (302 mg, 2 mmol, 1.00 eq) was reacted with diethanolaniline (544 mg, 3 mmol, 1.50 eq). After purification by means of flash chromatography (DCM:MeOH, 95:5) the desired product **methyl 3-((4-(bis(2-hydroxyethyl)amino)phenyl)diazenyl)benzoate (S11)** (200 mg, 0.58 mmol, 29%) was obtained as red solid.$^1$H NMR (400 MHz, chloroform-*d*) δ 8.49 (t, *J* = 1.8 Hz, 1H), 8.04 (tdd, *J* = 7.9, 2.4, 1.2 Hz, 2H), 7.89 (d, *J* = 8.7 Hz, 2H), 7.54 (t, *J* = 7.9 Hz, 1H), 6.80 (d, *J* = 8.9 Hz, 2H), 3.95 (d, *J* = 4.3 Hz, 7H), 3.82 (d, *J* = 18.8 Hz, 2H), 3.73 (t, *J* = 4.9 Hz, 4H). **$^{13}$C NMR** (101 MHz, MeOD-*d*$_4$) δ (ppm) = 168.3, 152.9, 132.6, 131.1, 130.6, 127.8, 126.7, 123.8, 113.0, 60.4, 55.2, 53.0. **LCMS(+):** $t_{ret}$ = 4.3 min, 344 Th = [MH]$^+$. **HRMS (EI)**: calc. for $[C_{18}H_{21}N_3O_4]^+$ = [M]$^+$: 343.1532; found: 343.1529.

By standard procedure D, **S11** (200 mg,0.58 mmol, 1.0 eq) was reacted to the desired product **3-((4-(bis(2-hydroxyethyl)amino)phenyl)diazenyl)benzoic acid** (**3DEA-$CO_2$H**) (154 mg, 0.47 mmol, 80%) was obtained as yellow solid. The compound has been reported[9] but no spectral data for comparison was available, so we report it here: **$^1$H NMR** (400 MHz, DMSO-*d*$_6$) δ (ppm) = 7.95 – 7.89 (m, 4H), 7.61 (dt, *J* = 6.4, 1.9 Hz, 5H), 4.83 (d, *J* = 21.9 Hz, 2H), 3.64 (s, 2H), 3.56 (d, *J* = 6.0 Hz, 2H), 3.47 (s, 3H). **$^{13}$C NMR** (101 MHz, DMSO-*d*$_6$) δ 170.3, 151.9, 151.7, 140.0, 131.8, 130.7, 129.6, 128.1, 122.7, 122.5, 58.5, 58.5, 51.6, 47.4. **LCMS(+):** $t_{ret}$ = 3.6 min, 330 Th = [MH]$^+$. **HRMS (EI)**: calc. for $[C_{17}H_{19}N_3O_4]^+$ **=** [M]$^+$: 329.1576; found: 329.1362.

**2-((4-methoxyphenyl)diazenyl)benzoic acid (2MP-$CO_2$H)**

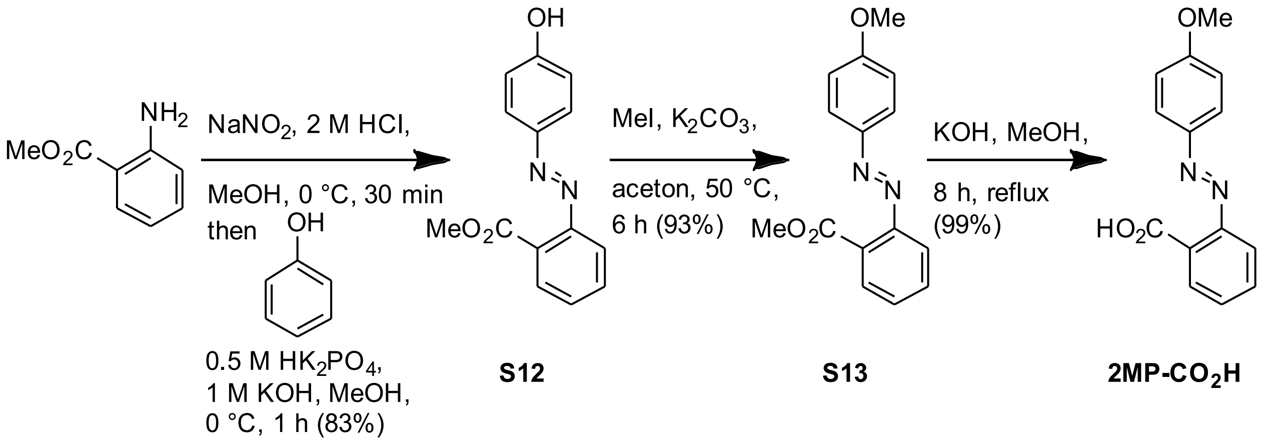

By standard procedure A, commercially available methyl 2-amino benzoate (302 mg, 2.0 mmol, 1.0 eq) was reacted with phenol (207 mg, 2.2 mmol, 1.1 eq). After purification by means of flash chromatography (EA:iHex, 9:1 → 7:3) the desired product **methyl 2-((4-hydroxyphenyl)diazenyl)benzoate** (**S12**) (423 mg, 1.8 mmol, 83%) was obtained as an orange solid. The compound has been reported but no spectral data for comparison were available so these are given here[10]: **$^1$H NMR** (400 MHz, DMSO-*d*$_6$) δ 7.79 – 7.75 (m, 2H), 7.74 (dd, *J* = 7.5, 1.4 Hz, 1H), 7.69 (ddd, *J* = 8.6, 7.1, 1.5 Hz, 1H), 7.63 (dd, *J* = 8.1, 1.4 Hz, 1H), 7.56 (td, *J* = 7.3, 1.5 Hz, 1H), 7.01 – 6.92 (m, 2H), 3.81 (s, 3H). **$^{13}$C NMR** (101 MHz, DMSO-*d*$_6$) δ (ppm) = 168.2, 161.9, 151.2, 145.7, 132.4, 130.1, 129.6, 128.8, 125.6, 119.8, 116.5, 52.7. **LCMS(+):** $t_{ret}$ =4.5 min, 257 Th = $[MH]^+$. **HRMS (EI):** calc. for $[C_{14}H_{12}N_2O_3{}^+]^+$ = $[M]^+$: 256.0848; found: 256.0838.

By standard procedure C, **S12** (402 mg, 1.57 mmol, 1.0 eq) was reacted with methyl iodide (668 mg, 4.7 mmol, 3.0 eq). After purification by means of flash chromatography (EA:iHex, 8:2) the desired product **methyl 2-((4-methoxyphenyl)diazenyl)benzoate** (**S13)** (394 mg, 0.53 mmol, 93%) was obtained as a yellow solid. The compound has been reported[11] but no spectral data for comparison were available so these are given here: **$^1$H NMR** (400 MHz, chloroform-*d*) δ (ppm) = 7.97 – 7.87 (m, 2H), 7.80 (ddd, *J* = 7.7, 1.4, 0.5 Hz, 1H), 7.64 – 7.54 (m, 2H), 7.45 (ddd, *J* = 7.7, 7.0, 1.6 Hz, 1H), 7.05 – 6.98 (m, 2H), 3.90 (d, *J* = 2.3 Hz, 6H).**$^{13}$C NMR** (101 MHz, chloroform-*d*) δ (ppm) = 168.2, 162.5, 152.1, 147.1, 131.9, 129.7, 129.2, 128.4, 125.2, 119.0, 114.3, 55.6, 52.3. **LCMS(+):** $t_{ret}$ = 5.4 min, 271 Th = $[MH]^+$. **HRMS (EI):** calc. for $[C_{15}H_{14}N_2O_3]^+$ = $[M]^+$: 270.1004; found: 270.1003.

By standard procedure D, **S15** (385 mg,0.53 mmol, 1.0 eq) was reacted to the desired product **2-((4-methoxyphenyl)diazenyl)benzoic acid** (**2MP-$CO_2$H**) (363 mg, 0.53 mmol, 99%) was obtained as yellow solid. Spectral data matches literature[12]: **$^1$H NMR** (400 MHz, chloroform-*d*) δ (ppm) = 7.93 – 7.89 (m, 2H), 7.88 (dd, *J* = 7.7, 1.5 Hz, 1H), 7.71 (dd, *J* = 8.0, 1.3 Hz, 1H), 7.66 – 7.59 (m, 1H), 7.54 (td, *J* = 7.5, 1.3 Hz, 1H), 7.12 – 7.07 (m, 2H), 3.90 (s, 3H). **$^{13}$C NMR** (101 MHz, chloroform-*d*) δ (ppm) = 170.0, 163.2, 150.9, 146.7, 131.6, 130.1, 129.7, 129.5, 125.0, 117.3, 114.2, 54.8. **LCMS(+):** $t_{ret}$ =4.9 min, 257 Th = $[MH]^+$. **HRMS (EI):** calc. for $[C_{14}H_{12}O_3N_2]^+$ = $[M]^+$: 256.0848; found: 256.0847.

#### *AzTaxs*

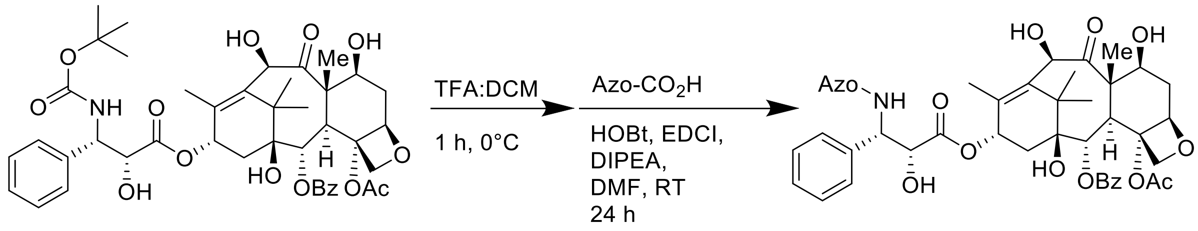

**(2aR,4S,4aS,6R,9S,11S,12S,12aR,12bS)-12b-acetoxy-4,6,11-trihydroxy-9-(((2R,3S)-2-hydroxy-3-phenyl-3-(4-(phenyldiazenyl)benzamido)propanoyl)oxy)-4a,8,13,13-tetramethyl-5-oxo-2a,3,4,4a,5,6,9,10,11,12,12a,12b-dodecahydro-1H-7,11-methanocyclodeca[3,4]benzo[1,2-b]oxet-12-yl benzoate (AzTax4H)**

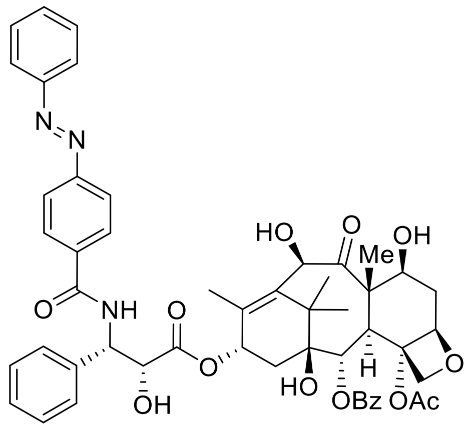

**AzTax4H**

By Standard Procedure E, docetaxel (25 mg, 31 µmol) was deprotected with TFA-DCM and the crude foam reacted with **4H-$CO_2$H** (7.5 mg, 33 µmol), DIPEA (11 mg, 86 µmol), EDCI (10.7 mg, 56 µmol), and HOBt.$H_2O$ (6.5 mg, 42 µmol) to yield a yellow crude solid. Chromatography on 5:1:0→1:1:0→1:1:0.2 iHex:EA:MeOH returned **AzTax4H** as a yellow solid (22 mg, 24 *µ*mol, 76 %). **$^1$H NMR** (400 MHz, chloroform-*d*) δ (ppm) = 8.16 – 8.09 (m, 2H), 7.95 – 7.91 (m, 2H), 7.68 – 7.31 (m, 14H), 7.28 (s, 1H), 6.24 – 6.17 (m, 1H), 5.79 (dd, *J* = 8.9, 2.8 Hz, 1H), 5.67 (d, *J* = 6.9 Hz, 1H), 5.18 (d, *J* = 4.9 Hz, 1H), 4.93 (dd, *J* = 9.6, 2.2 Hz, 1H), 4.80 (d, *J* = 2.9 Hz, 1H), 4.31 (d, *J* = 8.4 Hz, 1H), 4.24 – 4.16 (m, 2H), 3.89 (d, *J* = 7.2 Hz, 1H), 2.61 – 2.50 (m, 1H), 2.38 (s, 3H), 2.29 (dd, *J* = 9.0, 3.9 Hz, 2H), 1.89 – 1.79 (m, 2H), 1.78 – 1.73 (m, 6H), 1.72 – 1.68 (m, 1H), 1.20 (d, *J* = 6.4 Hz, 4H), 1.11 (s, 4H). **$^{13}$C NMR** (101 MHz, chloroform-*d*) δ (ppm) = 211.2, 172.5, 170.5, 167.0, 166.3, 154.5, 152.5, 138.1, 137.8, 136.1, 135.3, 133.8, 131.7, 130.2, 129.2, 129.1, 129.0, 128.8, 128.4, 128.1, 127.1, 123.1, 123.0, 84.1, 81.1, 78.7, 77.2, 74.7, 74.5, 73.2, 72.4, 72.0, 57.7, 55.2, 46.5, 43.0, 37.0, 35.9, 26.6, 22.6, 20.6, 14.4, 9.9. **LCMS(+):** $t_{ret}$ = 7.26 & 8.25 min, each 916 Th = $[MH]^+$, *Z* & *E* isomers respectively. **HRMS (ESI+)** calcd for $[C_{51}H_{54}N_3O_{13}]^+$ = $[MH]^+$: m/z 916.36566, found 916.36715.

**(2aR,4S,4aS,6R,9S,11S,12S,12aR,12bS)-12b-acetoxy-9-(((2R,3S)-3-(4-((4-(dimethylamino)phenyl)diazenyl)benzamido)-2-hydroxy-3-phenylpropanoyl)oxy)-4,6,11-trihydroxy-4a,8,13,13-tetramethyl-5-oxo-2a,3,4,4a,5,6,9,10,11,12,12a,12b-dodecahydro-1H-7,11-methanocyclodeca[3,4]benzo[1,2-b]oxet-12-yl benzoate (AzTax4DMA)**

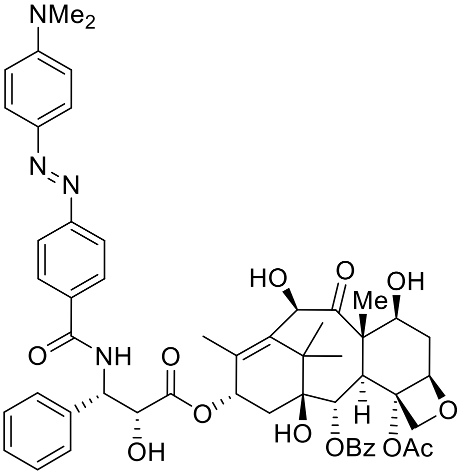

**AzTax4DMA**

By Standard Procedure E, docetaxel (23 mg, 28 µmol) was deprotected with TFA-DCM and the crude foam (20 mg) reacted with **4DMA-CO$_2$H** (11 mg, 41 µmol), Hünig base (11.1 mg, 85 µmol), EDCI (8.6 mg, 45 µmol), and HOBt·H$_2$O (6.9 mg, 45 µmol) to yield a yellow crude solid. Chromatography on 5:1:0→1:1:0→1:1:0.08 iHex:EA:MeOH returned **AzTax4DMA** as a yellow solid (13.8 mg, 14.4 *µ*mol, 51%).

**$^1$H NMR** (400 MHz, chloroform-*d*) δ (ppm) = 9.07 (d, *J* = 8.4 Hz, 1H), 8.05 (d, *J* = 8.6 Hz, 2H), 7.97 (d, *J* = 7.5 Hz, 2H), 7.86 (d, *J* = 8.4 Hz, 2H), 7.84 (d, *J* = 9.0 Hz, 2H), 7.78 – 7.68 (m, 1H), 7.69 – 7.59 (m ~t, *J* = 7.7 Hz, 2H), 7.46 – 7.41 (m, 2H), 7.43 – 7.38 (m, 2H), 7.23 (tt, *J* = 5.7, 2.9 Hz, 1H), 6.85 (d, *J* = 9.3 Hz, 2H), 6.23 (d, *J* = 7.8 Hz, 1H), 5.91 (t, *J* = 9.0 Hz, 1H), 5.39 (t, *J* = 8.6 Hz, 1H), 5.39 (d, *J* = 6.5 Hz), 5.09 (d, *J* = 2.6 Hz), 5.03 (d, *J* = 7.2 Hz, 1H), 4.98 (d, *J* = 2.4 Hz, 1H), 4.92 (dd, *J* = 9.7, 2.2 Hz, 1H), 4.57 (s, 1H), 4.59 (~t, *J* = 7.8 Hz, 1H), 4.11 – 3.96 (m, 3H, H10), 3.67 (d, *J* = 7.1 Hz, 1H), 3.09 (s, 6H), 2.35 – 2.25 (m, 1H), 2.23 (s, 3H), 1.90 – 1.81 (m, 1H), 1.73 – 1.65 (m, 1H), 1.74 (s, 3H), 1.72 – 1.63 (m, 1H), 1.53 (s, 3H), 1.01 (s, 3H), 0.98 (s, 3H) ppm. **$^{13}$C NMR** (101 MHz, chloroform-*d*) δ (ppm) = 209.7, 173.2, 170.2, 166.1, 165.7, 154.6, 153.3, 143.1, 139.7, 137.3, 136.2, 135.1, 133.9, 130.5, 130.0, 129.2, 129.0, 128.8, 128.0, 127.9, 125.6, 122.0, 112.0, 84.2, 80.7, 77.3, 75.9, 75.2, 74.2, 74.1, 71.3, 70.2, 57.4, 57.1, 46.4, 43.4, 40.6, 36.9, 35.4, 23.0, 21.5, 14.1, 10.3. **LCMS(+):** $t_{ret}$ = 8.39 min, 959 Th = [MH]$^+$, E isomer only. **HRMS (ESI+)** calcd for [$C_{53}H_{59}N_4O_{13}$]$^+$ = [MH]$^+$: m/z 959.40786, found 959.40758.

**(2aR,4S,4aS,6R,9S,11S,12S,12aR,12bS)-12b-acetoxy-4,6,11-trihydroxy-9-(((2R,3S)-2-hydroxy-3-(4-(4-methoxyphenyl)diazenyl)benzamido)-3-phenylpropanoyl)oxy)-4a,8,13,13-tetramethyl-5-oxo-2a,3,4,4a,5,6,9,10,11,12,12a,12b-dodecahydro-1H-7,11-methanocyclodeca[3,4]benzo[1,2-b]oxet-12-yl benzoate (AzTax4MP)**

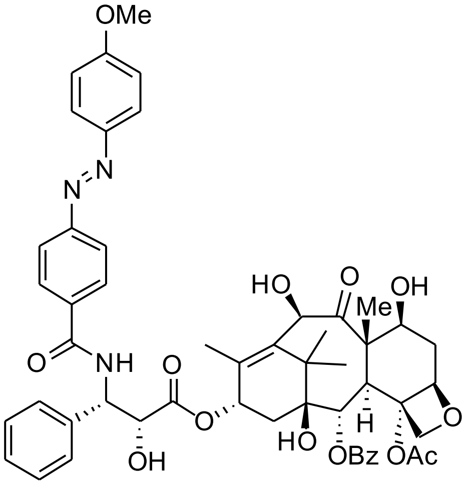

**AzTax4MP**

By Standard Procedure E, docetaxel (20 mg, 24 µmol) was deprotected with TFA-DCM and the crude foam (17 mg) reacted with **4MP-$CO_2$H** (8 mg, 28 µmol, 1.2 eq), Hünig base (12 mg, 96 µmol, 4.0 eq), EDCI (7 mg, 36 µmol), and HOBt·$H_2O$ (7 mg, 85 %wt, 39 µmol, 1.6 eq) to yield a yellow crude solid. Chromatography on (iHex:EA 7:3→1:1; DCM:MeOH 99:1→95:5) returned **AzTax4MP** as a yellow solid (10 mg, 10.4 *µ*mol, 41%). LCMS method was

**$^1$H NMR** (400 MHz, chloroform-*d*) δ (ppm) = 8.09 – 8.03 (m, 2H), 7.89 – 7.83 (m, 2H), 7.81 (d, *J* = 1.3 Hz, 3H), 7.58 – 7.51 (m, 1H), 7.49 – 7.39 (m, 4H), 7.39 – 7.32 (m, 2H), 7.32 – 7.25 (m, 1H), 7.12 (d, *J* = 9.0 Hz, 1H), 6.98 – 6.91 (m, 2H), 6.19 – 6.08 (m, 1H), 5.73 (dd, *J* = 9.0, 2.8 Hz, 1H), 5.61 (d, *J* = 7.0 Hz, 1H), 5.11 (d, *J* = 1.6 Hz, 1H), 4.91 – 4.83 (m, 1H), 4.73 (dd, *J* = 5.1, 2.8 Hz, 1H), 4.25 (d, *J* = 8.5 Hz, 1H), 4.19 – 4.10 (m, 3H), 3.83 (s, 4H), 3.55 (d, *J* = 5.3 Hz, 1H), 2.51 (ddd, *J* = 14.3, 9.6, 6.5 Hz, 1H), 2.32 (s, 3H), 2.23 (dd, *J* = 8.9, 5.1 Hz, 2H), 1.83 – 1.66 (m, 8H), 1.14 (s, 3H), 1.05 (s, 3H), 0.84 – 0.74 (m, 1H). **$^{13}$C NMR** (101 MHz, chloroform-*d*) δ (ppm) = 211.4, 172.7, 170.6, 167.1, 166.6, 162.8, 154.9, 147.1, 138.2, 138.0, 136.3, 134.8, 133.9, 130.3, 129.3, 129.2, 128.9, 128.5, 128.2, 127.2, 125.3, 122.9, 114.5, 84.3, 81.3, 78.9, 74.9, 74.7, 73.4, 72.6, 72.2, 57.8, 55.8, 55.3, 46.6, 43.2, 37.2, 36.1, 26.7, 22.7, 20.7, 14.5, 10.0. **LCMS(+):** tret =8.4 min, 946 Th = $[MH]^+$ **HRMS (ESI+)** calcd for $[C_{52}H_{56}N_3O_{14}]^+$ = $[MH]^+$: m/z 946.37568, found 946.37796.

**(2aR,4S,4aS,6R,9S,11S,12S,12aR,12bS)-12b-acetoxy-4,6,11-trihydroxy-9-(((2R,3S)-2-hydroxy-3-phenyl-3-(3-(phenyldiazenyl)benzamido)propanoyl)oxy)-4a,8,13,13-tetramethyl-5-oxo-2a,3,4,4a,5,6,9,10,11,12,12a,12b-dodecahydro-1H-7,11-methanocyclodeca[3,4]benzo[1,2-b]oxet-12-yl benzoate (AzTax3H)**

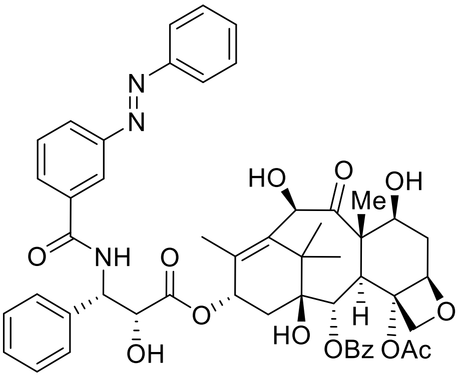

**3H**

By Standard Procedure E, docetaxel (40 mg, 50 µmol) was deprotected with TFA-DCM and the crude foam (32 mg) reacted with **3H-$CO_2$H** (5.3 mg, 23 µmol), Hünig base (6.3 mg, 49 µmol), EDCI (5.1 mg, 26 µmol), and HOBt·$H_2O$ (4.1 mg, 27 µmol) to yield a yellow crude solid (31 mg). Chromatography on 5:1:0→1:1:0→1:1:0.1 iHex:EA:MeOH returned **AzTax3H** as a yellow solid (12.2 mg, 13.3 *µ*mol, 58%).

**$^1$H NMR** (400 MHz, DMSO-*d*$_6$) δ (ppm) = 9.24 (d, *J* = 8.5 Hz, 1H), 8.44 (t, *J* = 1.9 Hz, 1H), 8.08 (t, J = 7.1 Hz, 2H), 8.00 – 7.92 (m, 4H), 7.77 – 7.69 (m, 2H), 7.64 (ddd, *J* = 7.9, 6.2, 2.1 Hz, 5H), 7.44 – 7.39 (m, 3H), 6.91 – 6.83 (m, 1H), 6.26 (d, *J* = 7.7 Hz, 1H), 5.96 – 5.88 (m, 1H), 5.45 – 5.38 (m, 2H), 5.09 (s, 1H), 5.05 – 4.94 (m, 2H), 4.91 (dd, *J* = 9.7, 2.2 Hz, 2H), 4.57 (d, *J* = 2.3 Hz, 1H), 4.08 – 3.95 (m, 4H), 3.72 – 3.62 (m, 1H), 2.21 (s, 3H), 2.09 (s, 1H), 1.75 (d, *J* = 1.4 Hz, 3H), 1.53 (s, 4H). **$^{13}$C NMR** (101 MHz, DMSO-*d*$_6$) δ (ppm) = 209.7, 173.2, 170.2, 165.9, 165.7, 153.8, 153.8, 152.3, 152.2, 139.5, 137.3, 136.2, 136.2, 133.9, 132.4, 130.8, 130.5, 130.1, 130.1, 130.0, 129.4, 129.1, 128.8, 128.0, 125.2, 123.1, 122.3, 120.4, 84.2, 83.4, 83.0, 80.9, 80.7, 77.3, 75.2, 74.2, 74.1, 71.3, 70.2, 57.4, 43.4, 41.2, 33.9, 28.7, 24.6, 21.5, 17.9, 17.2, 15.1, 14.1, 10.3, 8.3. **HRMS (ESI+)** calcd for $[C_{51}H_{54}N_3O_{13}]^+$ = $[MH]^+$: m/z 916.36566, found 916.36526.

**(2aR,4S,4aS,6R,9S,11S,12S,12aR,12bS)-12b-acetoxy-9-(((2R,3S)-3-(3-((4-(dimethylamino)phenyl)diazenyl)benzamido)-2-hydroxy-3-phenylpropanoyl)oxy)-4,6,11-trihydroxy-4a,8,13,13-tetramethyl-5-oxo-2a,3,4,4a,5,6,9,10,11,12,12a,12b-dodecahydro-1H-7,11-methanocyclodeca[3,4]benzo[1,2-b]oxet-12-yl benzoate (AzTax3DMA)**

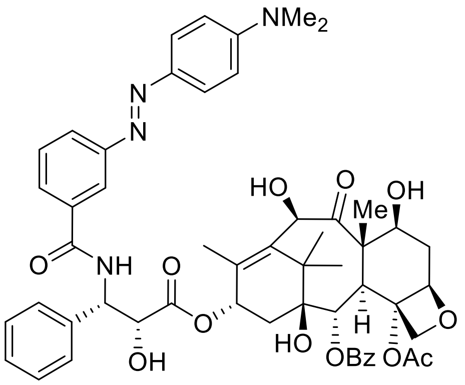

**AzTax3DMA**

By Standard Procedure E, docetaxel (20 mg, 24 µmol) was deprotected with TFA-DCM and the crude foam (17 mg) reacted with **3DMA-$CO_2$H** (8 mg, 28 µmol, 1.2 eq), Hünig base (12 mg, 96 µmol, 4.0 eq), EDCI (7 mg, 36 µmol), and HOBt·$H_2O$ (7 mg, 85 %wt, 39 µmol, 1.6 eq) to yield a yellow crude solid. Chromatography on (iHex:EA 7:3→1:1; DCM:MeOH 99:1→95:5) returned **AzTax3DMA** as a yellow solid (8 mg, 8.3 *µ*mol, 35%).

**$^1$H NMR** (400 MHz, chloroform-*d*) δ (ppm) = 8.17 – 8.10 (m, 3H), 7.98 – 7.92 (m, 1H), 7.89 – 7.82 (m, 2H), 7.79 (dt, J = 8.0, 1.3 Hz, 1H), 7.62 – 7.55 (m, 1H), 7.54 – 7.46 (m, 5H), 7.46 – 7.39 (m, 2H), 7.38 – 7.33 (m, 1H), 7.18 (d, J = 9.0 Hz, 1H), 6.79 – 6.71 (m, 2H), 6.23 (t, J = 8.9 Hz, 1H), 5.85 – 5.79 (m, 1H), 5.69 (d, J = 7.1 Hz, 1H), 5.17 (s, 1H), 4.97 – 4.90 (m, 1H), 4.80 (s, 1H), 4.31 (d, J = 8.5 Hz, 1H), 4.27 – 4.14 (m, 3H), 3.91 (d, J = 7.1 Hz, 1H), 3.65 (s, 1H), 3.10 (s, 6H), 2.57 (ddd, J = 15.2, 9.6, 6.5 Hz, 1H), 2.40 (s, 3H), 2.37 – 2.23 (m, 2H), 1.91 – 1.83 (m, 1H), 1.80 (d, J = 1.4 Hz, 3H), 1.76 (s, 3H), 1.58 (s, 1H), 1.22 (s, 3H), 1.12 (s, 3H), 0.88 (dd, J = 12.4, 8.0 Hz, 1H). **$^{13}$C NMR** (101 MHz, chloroform-*d*) δ (ppm) = 211.4, 172.6, 170.5, 167.0, 166.8, 153.2, 152.8, 143.4, 138.2, 137.9, 136.1, 134.5, 133.7, 130.2, 129.5, 129.2, 129.0, 128.8, 128.4, 128.0, 127.1, 125.5, 125.3, 120.4, 111.5, 84.1, 81.1, 78.8, 74.8, 74.5, 73.3, 72.5, 72.0, 57.7, 55.1, 46.5, 43.1, 40.3, 37.1, 36.0, 26.6, 22.6, 20.6, 14.4, 9.9. **LCMS(+):** $t_{ret}$ =8.7 min, 959 Th = $[M]^+$. **HRMS (ESI+)** calcd for $[C_{53}H_{59}N_4O_{13}]^+$ = $[MH]^+$: m/z 959.40731, found 959.40885.

**(2a*R*,4*S*,4a*S*,6*R*,9*S*,11*S*,12*S*,12a*R*,12b*S*)-12b-acetoxy-4,6,11-trihydroxy-9-(((2*R*,3*S*)-2-hydroxy-3-(3-((*E*)-(4-methoxyphenyl)diazenyl)benzamido)-3-phenylpropanoyl)oxy)-4a,8,13,13-tetramethyl-5-oxo-2a,3,4,4a,5,6,9,10,11,12,12a,12b-dodecahydro-1*H*-7,11-methanocyclodeca[3,4]benzo[1,2-*b*]oxet-12-yl benzoate (AzTax3MP)**

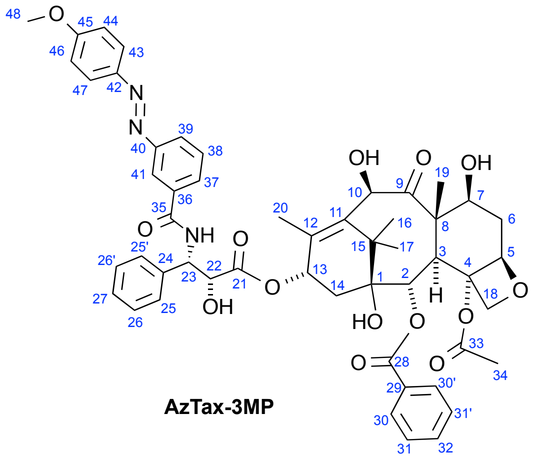

**AzTax-3MP**

By Standard Procedure E, docetaxel (21 mg, 26 µmol) was deprotected with TFA-DCM and the crude foam reacted with **3MP-$CO_2$H** (7.3 mg, 28 µmol), Hünig base (9.4 mg, 73 µmol), EDCI (7.1 mg, 37 µmol), and HOBt·$H_2O$ (5.5 mg, 36 µmol) to yield a yellow crude solid. Chromatography on 5:1:0→1:1:0→1:1:0.2 iHex:EA:MeOH returned **AzTax3MP** as a yellow solid (20 mg, 21 *µ*mol, 81%).

**$^1$H NMR** (400 MHz, DMSO-$d_6$) δ (ppm) = 8.13 (~t, *J* = 1.8 Hz, 1H), 8.05 (d, *J* = 7.8 Hz, 2H), 7.92 (d, *J* = 7.0 Hz, 1H), 7.83 (d, *J* = 9.0 Hz, 2H), 7.77 (d, *J* = 7.7 Hz, 1H), 7.51 (~t, *J* = 7.5 Hz, 1H), 7.45 (t, *J* = 7.6 Hz, 1H), 7.48 – 7.34 (m, 2H), 7.43 – 7.39 (m, 2H), 7.35 (~t, *J* = 7.5 Hz, 2H), 7.27 (t, *J* = 7.4 Hz, 1H), 6.94 (d, *J* = 9.0 Hz, 2H), 6.15 (t, *J* = 8.7 Hz, 1H), 5.75 (dd, *J* = 8.9, 2.8 Hz, 1H), 5.61 (d, *J* = 7.0 Hz, 1H), 5.11 (s, 1H), 4.86 (~d, *J* = 9.5 Hz, 1H), 4.73 (d, *J* = 2.8 Hz, 1H), 4.24 (d, J = 8.5 Hz, 1H), 4.16-4.09 (m, 2H), 3.82 (s, 3H), 3.85 – 3.80 (m overlapped, 1H), 2.56 – 2.43 (m, 1H), 2.32 (s, 3H), 2.27 – 2.19 (m, 1H), 1.89 – 1.72 (m, 1H), 1.72 – 1.64 (m, 1H), 1.71 (s, 3H), 1.68 (s, 3H, 3H19), 1.13 (s, 3H,), 1.04 (s, 3H). **$^{13}$C NMR** (101 MHz, DMSO-$d_6$) δ (ppm) = 211.3 (C9), 172.6 (C21), 170.5 (C33), 166.9 (C35), 166.6 (C28), 162.5 (C45), 152.7 (C40), 146.7 (C42), 138.1 (C12), 137.9 (C11), 136.1 (C36), 134.7 (C24), 133.7 (C32), 130.2 (C30 & C30'), 129.5 (C29), 129.1 (C37), 129.0 (C31 & C31'), 128.9 (C38), 128.7 (C26 & C26'), 128.4 (C27), 127.1 (25 & 25'), 125.8 (C39), 125.1 (C43 & C47), 121.0 (C41), 114.3 (C44 & C46), 84.2 (C5), 81.1 (C4), 78.7 (C1), 77.2 (C18), 74.8 (C2), 74.5 (C7), 73.2 (C22), 72.4 (C10), 72.0 (C13), 57.7 (C48), 55.6 (C23), 55.2 (C8), 46.5 (C3), 43.0 (C15), 37.0 (C6), 36.0 (C14), 26.6 (C34), 22.6 (C16), 20.6 (C17), 14.4 (C20), 9.9 (C19). **LCMS(+):** $t_{ret}$ = 7.20 & 8.21 min, each 946 Th = [MH]$^+$, Z & E isomers respectively. **HRMS (ESI+)** calcd for $[C_{52}H_{56}N_3O_{14}]^+$ = [MH]$^+$: m/z 946.37623, found 946.37733.

**(2aR,4S,4aS,6R,9S,11S,12S,12aR,12bS)-12b-acetoxy-4,6,11-trihydroxy-9-(((2R,3S)-2-hydroxy-3-(4-methoxy-3-((3,4,5-trimethoxyphenyl)diazenyl)benzamido)-3-phenylpropanoyl)oxy)-4a,8,13,13-tetramethyl-5-oxo-2a,3,4,4a,5,6,9,10,11,12,12a,12b-dodecahydro-1H-7,11-methanocyclodeca[3,4]benzo[1,2-b]oxet-12-yl benzoate (AzTax3MTM)**

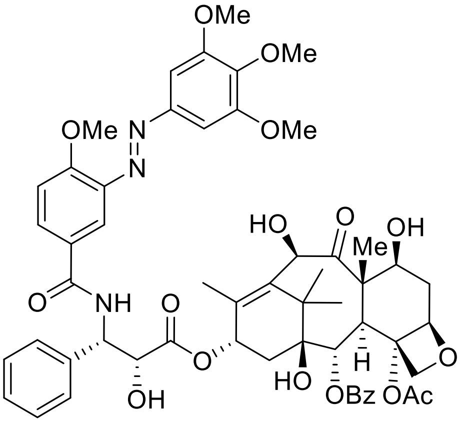

**AzTax3MTM**

By Standard Procedure E, docetaxel (20 mg, 24 µmol) was deprotected with TFA-DCM and the crude foam (17 mg) reacted with **3MTM-CO$_2$H** (8 mg, 28 µmol, 1.2 eq), Hünig base (12 mg, 96 µmol, 4.0 eq), EDCI (7 mg, 36 µmol), and HOBt·$H_2O$ (7 mg, 85 %wt, 39 µmol, 1.6 eq) to yield a yellow crude solid. Chromatography on (iHex:EA 7:3→1:1; DCM:MeOH 99:1→95:5) returned **AzTax3MTM** as a yellow solid (8 mg, 8.3 *µ*mol, 35%).

**$^1$H NMR** (400 MHz, chloroform-*d*) δ (ppm) = 8.14 – 8.06 (m, 3H), 7.94 – 7.87 (m, 2H), 7.58 – 7.52 (m, 1H), 7.48 (td, *J* = 8.6, 8.1, 1.6 Hz, 4H), 7.43 (t, *J* = 1.7 Hz, 1H), 7.42 – 7.39 (m, 2H), 7.39 – 7.32 (m, 2H), 7.19 (s, 2H), 7.13 – 7.09 (m, 1H), 7.07 (d, *J* = 8.7 Hz, 1H), 6.26 – 6.18 (m, 1H), 5.82 (dd, *J* = 9.0, 2.7 Hz, 1H), 5.69 (d, *J* = 7.0 Hz, 1H), 5.17 (s, 1H), 4.92 (d, *J* = 8.8 Hz, 1H), 4.81 (d, *J* = 2.6 Hz, 1H), 4.30 (d, *J* = 8.4 Hz, 1H), 4.24 – 4.19 (m, 3H), 4.03 (s, 3H), 3.97 (q, *J* = 2.2, 1.6 Hz, 2H), 3.94 (s, 6H), 3.93 (s, 3H), 2.56 (ddd, *J* = 14.2, 9.5, 6.6 Hz, 2H), 2.40 (s, 3H), 2.29 – 2.19 (m, 2H), 1.89 – 1.84 (m, 3H), 1.80 (d, *J* = 1.4 Hz, 3H), 1.76 (s, 3H), 1.21 (s, 3H), 1.11 (s, 3H). **$^{13}$C NMR** (101 MHz, CDCl$_3$) δ (ppm) = 211.4, 172.7, 170.7, 167.0, 166.3, 162.8, 159.3, 153.6, 148.9, 141.8, 141.2, 138.2, 138.1, 136.2, 133.8, 131.2, 130.3, 129.3, 129.1, 128.8, 128.4, 127.2, 126.1, 116.0, 112.6, 101.0, 84.3, 81.2, 78.8, 75.0, 74.6, 73.4, 72.6, 72.1, 61.2, 57.8, 56.6, 56.4, 55.2, 46.6, 43.2, 37.1, 36.7, 36.3, 31.6, 29.8, 26.7, 22.7, 20.8, 14.5, 10.0.

**(2aR,4S,4aS,6R,9S,11S,12S,12aR,12bS)-12b-acetoxy-4,6,11-trihydroxy-9-(((2R,3S)-2-hydroxy-3-phenyl-3-(3-((3,4,5-trimethoxyphenyl)diazenyl)benzamido)propanoyl)oxy)-4a,8,13,13-tetramethyl-5-oxo-2a,3,4,4a,5,6,9,10,11,12,12a,12b-dodecahydro-1H-7,11-methanocyclodeca[3,4]benzo[1,2-b]oxet-12-yl benzoate (AzTax3TM)**

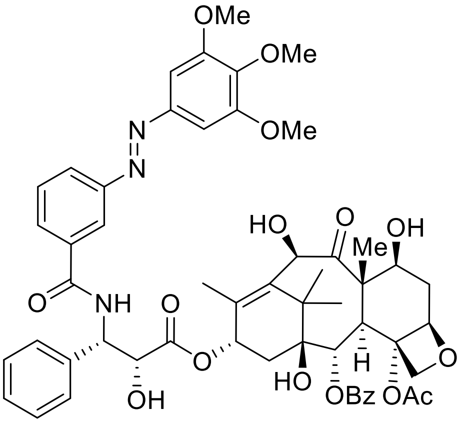

**AzTax3TM**

By Standard Procedure E, docetaxel (20 mg, 24 µmol) was deprotected with TFA-DCM and the crude foam (17 mg) reacted with **3TM-$CO_2$H** (10 mg, 29 µmol, 1.2 eq), Hünig base (12 mg, 96 µmol, 4.0 eq), EDCI (7 mg, 36 µmol), and HOBt·$H_2O$ (7 mg, 85 %wt, 39 µmol, 1.6 eq) to yield a yellow crude solid. Chromatography on (iHex:EA 7:3→1:1; DCM:MeOH 99:1→96:4) returned **AzTax3TM** as a yellow solid (9 mg, 9.0 *µ*mol, 37%).

**$^1$H NMR** (400 MHz, chloroform-d) δ (ppm) = 8.22 (t, J = 1.9 Hz, 1H), 8.15 – 8.10 (m, 2H), 8.04 – 7.99 (m, 1H), 7.87 (dt, J = 7.9, 1.3 Hz, 1H), 7.62 – 7.54 (m, 2H), 7.54 – 7.48 (m, 4H), 7.47 – 7.44 (m, 1H), 7.44 – 7.39 (m, 2H), 7.39 – 7.35 (m, 1H), 7.24 (s, 2H), 7.20 (dd, J = 8.7, 3.9 Hz, 1H), 6.26 – 6.20 (m, 1H), 5.84 (dd, J = 9.0, 2.6 Hz, 1H), 5.69 (d, J = 7.1 Hz, 1H), 5.17 (d, J = 4.7 Hz, 1H), 4.97 – 4.90 (m, 1H), 4.82 (s, 1H), 4.31 (d, J = 8.5 Hz, 1H), 4.22 (d, J = 8.2 Hz, 3H), 3.96 (s, 6H), 3.94 (s, 3H), 3.93 – 3.87 (m, 2H), 3.69 – 3.60 (m, 2H), 3.58 (s, 1H), 2.57 (ddd, J = 15.7, 9.7, 6.5 Hz, 1H), 2.40 (s, 3H), 2.37 – 2.21 (m, 3H), 1.86 (d, J = 12.4 Hz, 2H), 1.80 (d, J = 1.4 Hz, 2H), 1.76 (s, 3H), 1.21 (s, 4H), 1.12 (s, 3H), 0.93 – 0.80 (m, 2H). **$^{13}$C NMR** (101 MHz, chloroform-d) δ (ppm) = 211.3, 172.6, 170.5, 167.0, 166.4, 152.5, 148.2, 141.2, 138.1, 137.9, 136.2, 134.7, 133.7, 130.2, 129.6, 129.2, 129.2, 129.1, 128.7, 128.4, 127.1, 126.0, 121.0, 100.8, 84.1, 81.1, 78.8, 74.8, 74.5, 73.1, 72.5, 72.0, 61.1, 57.7, 56.3, 55.1, 46.5, 43.1, 37.0, 36.0, 29.7, 26.6, 22.6, 20.6, 14.4, 9.9. **LCMS(+):** $t_{ret}$ =8.2 min, 1006 Th = [M]$^+$. **HRMS (ESI+)** calcd for [$C_{55}H_{59}N_3O_{16}$]$^+$ = [MH]$^+$: m/z 1006.39681, found 1006.39929.

**(2aR,4S,4aS,6R,9S,11S,12S,12aR,12bS)-12b-acetoxy-9-(((2R,3S)-3-(3-((4-(bis(2-hydroxyethyl)amino)phenyl)diazenyl)benzamido)-2-hydroxy-3-phenylpropanoyl)oxy)-4,6,11-trihydroxy-4a,8,13,13-tetramethyl-5-oxo-2a,3,4,4a,5,6,9,10,11,12,12a,12b-dodecahydro-1H-7,11-methanocyclodeca[3,4]benzo[1,2-b]oxet-12-yl benzoate (AzTax3DEA)**

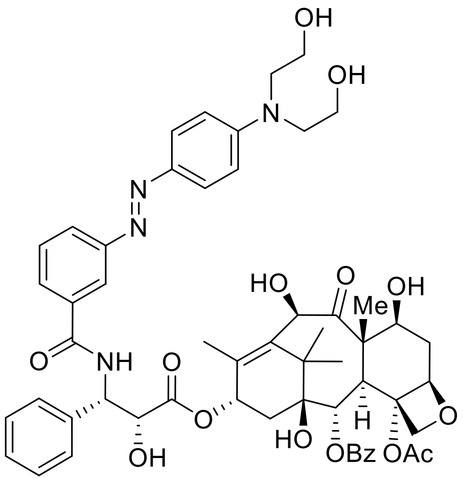

**AzTax3DEA**

By Standard Procedure E, docetaxel (20 mg, 24 µmol) was deprotected with TFA-DCM and the crude foam (17 mg) reacted with **3DEA-CO$_2$H** (10 mg, 29 µmol, 1.2 eq), Hünig base (12 mg, 96 µmol, 4.0 eq), EDCI (7 mg, 36 µmol), and HOBt·H$_2$O (7 mg, 85 %wt, 39 µmol, 1.6 eq) to yield a yellow crude solid. Chromatography on (DCM:MeOH 98:2→92:8) returned **AzTax3DEA** as a yellow solid (12 mg, 11.8 *µ*mol, 49%).

**$^1$H NMR** (400 MHz, methanol-*d*$_4$) δ (ppm) = 8.29 (t, *J* = 1.8 Hz, 1H), 8.12 (d, *J* = 1.2 Hz, 1H), 8.10 (d, *J* = 1.5 Hz, 1H), 7.96 (ddd, *J* = 8.0, 2.0, 1.1 Hz, 1H), 7.89 (dt, *J* = 7.8, 1.4 Hz, 1H), 7.85 – 7.81 (m, 2H), 7.68 – 7.62 (m, 1H), 7.61 – 7.55 (m, 3H), 7.55 – 7.52 (m, 1H), 7.50 (d, *J* = 1.2 Hz, 1H), 7.43 (t, *J* = 7.8 Hz, 2H), 7.33 – 7.27 (m, 1H), 6.89 (d, *J* = 9.3 Hz, 2H), 6.27 – 6.17 (m, 1H), 5.69 (d, *J* = 5.3 Hz, 1H), 5.64 (d, *J* = 7.2 Hz, 1H), 5.25 (s, 1H), 4.99 – 4.94 (m, 1H), 4.76 (d, *J* = 5.4 Hz, 1H), 4.58 (s, 1H), 4.20 (td, *J* = 8.6, 5.9 Hz, 3H), 3.88 (d, *J* = 7.2 Hz, 1H), 3.79 (t, *J* = 5.9 Hz, 4H), 3.68 (t, *J* = 5.9 Hz, 4H), 2.48 – 2.40 (m, 1H), 2.39 (s, 3H), 2.30 – 2.18 (m, 2H), 1.96 (dd, *J* = 15.5, 8.8 Hz, 1H), 1.89 (d, *J* = 1.4 Hz, 3H), 1.82 (td, *J* = 12.6, 11.3, 2.6 Hz, 1H), 1.69 (s, 3H), 1.16 (s, 3H), 1.11 (s, 3H). **$^{13}$C NMR** (101 MHz, methanol-*d*$_4$) δ (ppm) = 209.7, 173.1, 170.5, 170.0, 168.3, 166.3, 153.2, 151.2, 143.3, 137.8, 136.6, 130.1, 129.8, 129.0, 128.4, 128.3, 127.8, 127.6, 127.1, 125.0, 120.5, 111.4, 84.6, 80.9, 77.8, 76.2, 75.1, 74.2, 73.6, 71.2, 71.1, 58.9, 57.5, 56.5, 53.6, 46.4, 43.1, 36.1, 35.5, 25.6, 21.9, 13.0, 9.1. **LCMS(+):** $t_{ret}$ =7.3 min, 1019 Th = [M]$^+$.

**(2aR,4S,4aS,6R,9S,11S,12S,12aR,12bS)-12b-acetoxy-4,6,11-trihydroxy-9-(((2R,3S)-2-hydroxy-3-(2-((4-methoxyphenyl)diazenyl)benzamido)-3-phenylpropanoyl)oxy)-4a,8,13,13-tetramethyl-5-oxo-2a,3,4,4a,5,6,9,10,11,12,12a,12b-dodecahydro-1H-7,11-methanocyclodeca[3,4]benzo[1,2-b]oxet-12-yl benzoate (AzTax2MP)**

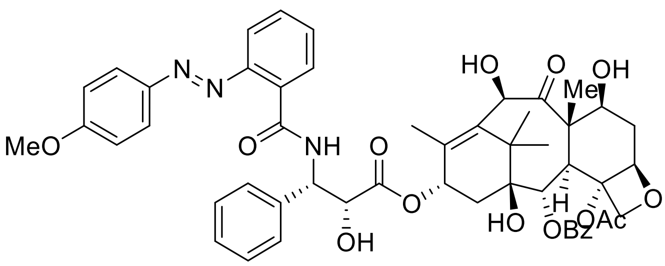

**AzTax2MP**

By Standard Procedure E, docetaxel (20 mg, 24 µmol) was deprotected with TFA-DCM and the crude foam (17 mg) was dissolved in 2 mL DMF. **4MP-CO$_2$H** (8 mg, 28 µmol, 1.2 eq) was added to the reaction mixture. Triethylamine (24 mg, 240 *µ*mol, 10 eq) was added. T3P (26 mg, 50 wt% in EA, 40 *µ*M, 1.7 eq) was added. The resulting organic solution was stirred at room temperature for 16 h. Upon completion the DMF was removed *in* vacuo and the resulting yellow crude product was purified by means of flash chromatography on silica (iHex:EA 7:3→1:1; DCM:MeOH 99:1→95:5). **AzTax2MP** was obtained as a yellow solid (10 mg, 10.4 *µ*mol, 41%).

**$^1$H NMR** (400 MHz, chloroform-*d*) δ (ppm) = 9.69 (d, *J* = 8.6 Hz, 1H), 8.28 (dd, *J* = 7.9, 1.6 Hz, 1H), 8.20 – 8.12 (m, 2H), 7.85 – 7.80 (m, 2H), 7.78 (dd, *J* = 8.2, 1.3 Hz, 1H), 7.66 – 7.60 (m, 1H), 7.56 – 7.47 (m, 5H), 7.42 (td, *J* = 7.6, 1.3 Hz, 1H), 7.38 – 7.30 (m, 3H), 6.96 (d, *J* = 9.0 Hz, 2H), 6.22 (d, *J* = 8.8 Hz, 1H), 5.95 (dd, *J* = 8.7, 2.5 Hz, 1H), 5.68 (d, *J* = 7.0 Hz, 1H), 5.16 – 5.12 (m, 1H), 4.94 (d, *J* = 9.4 Hz, 1H), 4.77 (dd, *J* = 4.9, 2.5 Hz, 1H), 4.33 (d, *J* = 8.5 Hz, 1H), 4.25 – 4.16 (m, 3H), 3.89 (s, 4H), 3.69 (d, *J* = 7.6 Hz, 2H), 2.58 (t, *J* = 15.2 Hz, 2H), 2.42 (s, 3H), 2.40 – 2.22 (m, 3H), 1.86 (d, *J* = 14.3 Hz, 2H), 1.80 (d, *J* = 1.4 Hz, 3H), 1.77 (d, *J* = 5.5 Hz, 3H), 1.51 (s, 2H), 1.19 (s, 3H), 1.11 (s, 3H), 0.95 – 0.81 (m, 2H).

**$^{13}$C NMR** (101 MHz, chloroform-*d*) δ (ppm) = 172.5, 170.5, 167.0, 165.9, 163.2, 150.1, 146.7, 138.7, 138.5, 135.9, 133.7, 132.2, 131.9, 130.7, 130.3, 129.4, 129.3, 128.9, 128.8, 128.3, 128.3, 128.0, 127.2, 126.8, 125.8, 116.1, 114.6, 84.1, 81.1, 78.8, 74.8, 74.6, 73.8, 72.3, 72.1, 57.7, 55.7, 46.5, 43.0, 37.0, 36.0, 29.7, 26.5, 22.7, 20.6, 14.6, 9.9. **LCMS(+):** $t_{ret}$ =8.7 min, 946 Th = [MH]$^+$, **HRMS (ESI+)** calcd for [$C_{52}H_{55}N_3O_{14}$]$^+$ = [MH]$^+$: m/z 946.37740, found 946.37568.

#### *Water-soluble model photoswitch carboxamides*

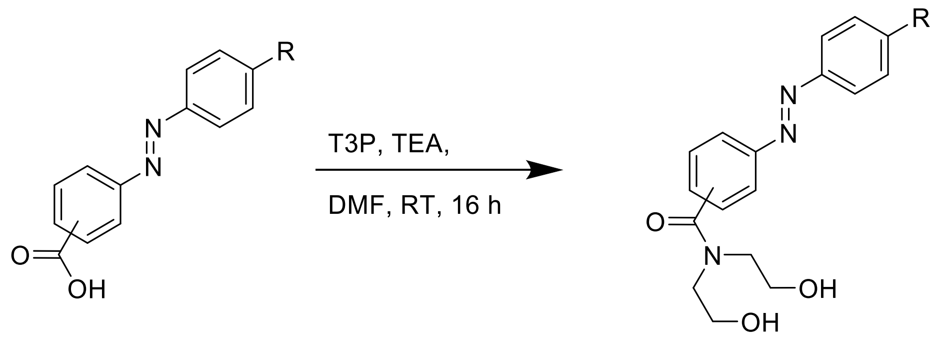

para, R: H, **4H**
meta, R:OMe, **3MP**
meta, R: $NMe_2$, **3DMA**

##### ***N,N*-bis(2-hydroxyethyl)-4-(phenyldiazenyl)benzamide (4H)**

By standard procedure F, Commercial compound **4H-$CO_2$H** (20 mg, 0.089 mmol, 1.0 eq) was reacted with diethanolamine (19 mg, 0.18 mmol, 2.0 eq). After purification by means of flash chromatography (DCM:MeOH, 100:0→98:2) the desired product ***N,N*-bis(2-hydroxyethyl)-4-(phenyldiazenyl)benzamide** (**4H**). **$^1$H NMR** (400 MHz, methanol-*d*$_4$) δ (ppm) = 8.01 – 7.97 (m, 2H), 7.97 – 7.92 (m, 2H), 7.68 – 7.63 (m, 2H), 7.60 – 7.51 (m, 3H), 3.87 (t, *J* = 5.5 Hz, 2H), 3.74 (t, *J* = 5.6 Hz, 2H), 3.64 (t, *J* = 5.6 Hz, 2H), 3.54 (t, *J* = 5.7 Hz, 2H). **$^{13}$C NMR** (101 MHz, DMSO-*d*$_6$) δ (ppm) = 170.3, 151.9, 151.7, 140.0, 131.8, 130.7, 129.6, 128.1, 122.7, 122.5, 58.5, 58.5, 51.6, 47.4. **LCMS(+):** $t_{ret}$ = 3.8 min, 314 Th = [MH]$^+$. **HRMS (EI):** calc. for $C_{17}H_{19}N_3O_3^+$ [M]$^+$: 313.1426; found: 343.1409.

##### **3-((4-(dimethylamino)phenyl)diazenyl)-*N,N*-bis(2-hydroxyethyl)benzamide (3DMA)**

By standard procedure F, **3DMA-$CO_2$H** (20 mg, 0.074 mmol, 1.0 eq) was reacted with diethanolamine (16 mg, 0.15 mmol, 2.0 eq). After purification by means of flash chromatography (DCM:MeOH, 98:2→96:4) the desired product **3-((4-(dimethylamino)phenyl)diazenyl)-N,N-bis(2-hydroxyethyl)benzamide** (**3DMA**) (16 mg, 0.045 mmol, 60%) was obtained as a yellow solid. **$^1$H NMR** (500 MHz, methanol-*d*$_4$) δ (ppm) = 7.92 – 7.81 (m, 4H), 7.57 (dd, *J* = 8.5, 7.6 Hz, 1H), 7.48 (dt, *J* = 7.5, 1.4 Hz, 1H), 6.86 – 6.81 (m, 2H), 3.88 (t, *J* = 5.7 Hz, 2H), 3.74 (t, *J* = 5.7 Hz, 2H), 3.64 (t, *J* = 5.8 Hz, 2H), 3.54 (t, *J* = 5.7 Hz, 2H), 3.10 (s, 6H). **$^{13}$C NMR** (126 MHz, methanol-*d*$_4$) δ (ppm) = 173.1, 165.0, 153.1, 153.0, 143.3, 137.5, 129.0, 127.2, 124.8, 123.0, 119.8, 111.2, 59.2, 59.0, 52.3, 39.0. **LCMS(+):** $t_{ret}$ = 4.0 min, 357 Th = [MH]$^+$. **HRMS (EI):** calc. for $C_{14}H_{12}O_3N_2^+$ [M]$^+$: 356.1848; found: 356.1839.

##### ***N,N*-bis(2-hydroxyethyl)-3-((4-methoxyphenyl)diazenyl)benzamide (3MP)**

By standard procedure F, **3MP-$CO_2$H** (20 mg, 0.078 mmol, 1.0 eq) was reacted with diethanolamine (16 mg, 0.16 mmol, 2.0 eq). After purification by means of flash chromatography (DCM:MeOH, 98:2→95:5) the desired product **N,N-bis(2-hydroxyethyl)-3-((4-methoxyphenyl)diazenyl)benzamide** (**3MP**) (17 mg, 0.050 mmol, 64%) was obtained as a yellow solid. **$^{1}$H NMR** (400 MHz, methanol-*$d_4$*) δ (ppm) = 7.97 – 7.90 (m, 4H), 7.64 – 7.58 (m, 1H), 7.56 (dt, J = 7.6, 1.5 Hz, 1H), 7.12 – 7.04 (m, 2H), 3.89 (s, 5H), 3.74 (t, J = 5.7 Hz, 2H), 3.64 (t, J = 5.7 Hz, 2H), 3.53 (t, J = 5.7 Hz, 2H). **$^{13}$C NMR** (101 MHz, methanol-*$d_4$*) δ (ppm) = 172.1, 162.0, 151.7, 145.9, 136.8, 128.4, 127.6, 123.8, 122.6, 119.5, 113.2, 58.4, 58.1, 54.0, 51.5. **LCMS(+):** $t_{ret}$ = 3.6 min, 344 Th = $[MH]^{+}$. **HRMS (EI):** calc. for $C_{14}H_{12}O_{3}N_{2}^{+}$ $[M]^{+}$: 343.1532; found: 343.1522.

##### *Discussion of arylazopyrazole-dextran heterodimerisation (Liu et al., 2018)*

As far as we are aware, Liu and coworkers (Angewandte 2018)[13] disclosed the only work in the direction of optically-localisable microtubule stabilisation. They aimed to use the known photoswitchability of host-guest interactions between beta-cyclodextran and [1]arylazopyrazoles (*trans*-arylazopyrazole:dextran binding constant up to 2300 $M^{-1}$ while the *cis*- is essentially nonbinding)[14] to aim at photoswitchably reversible, noncovalent heterodimerisation of two paclitaxel conjugates each applied to cells at 100 nM. Their hypothesis (as Fig 1[13]) is that the heterodimer (max taxol-to-taxol distance approx 2 nm) should allow crosslinking binding to two microtubules (minimum taxol-to-taxol distance ca. 7 nm), but this is to our mind geometrically unlikely since taxol's binding site is on the luminal (inner) face of the microtubules[15] and crosslinked binding would require the linker to penetrate directly through both protein walls, as well as stretching substantially beyond its limit end-to-end distance and reorienting taxol away from its usual binding structure. It is also noteworthy that the maximum concentration of heterodimer in free solution that could be predicted from this approach using the literature binding constant is only ca. 23 pM; and that the separated monomeric paclitaxel halves counterintuitively do not appear to hyperpolymerise microtubule structure *in cellulo* (as Fig 4[13]) which may reflect poor potency when the 2'-hydroxyl group, usually thought necessary for binding as it is oriented into the protein, is masked with large although potentially enzymatically cleavable esters. Nevertheless, the authors reported that the combination of the two heterodimeriser halves isomer-dependently alters the proportion of subG1-phase (dying) cells in a treated sample from 8% to 12% (error bars ± 2%; as Fig S14[13]); but this approach has not been shown to allow *in cellulo* photoswitchability of effect, nor an apparent switch-on/switch-off of bioactivity. To our mind it cannot be excluded that the relatively modest isomer-dependency of cellular effect results from slightly improved dextran trafficking of the *trans*-conjugate and also invokes intracellular ester hydrolysis; nor that the differences of cell-free MT aggregate shapes (as Fig 2[13]), if such are reproducible and significant, could arise from the differential solubilities of the species employed. The choice of a ~4 kDa construct design with two applied drugs also seems to leave room for reagent design improvement.

### Part B: Photocharacterisation *in vitro*

#### *Materials and Methods*

##### HPLC for UV-Vis spectroscopy on separated isomers

During HPLC (as in Part A), the diode array detector was used to acquire peak spectra of separated photoswitch isomers over the range 200–550 nm, manually baselining across each elution peak of interest to correct for eluent composition effects.

##### UV-Vis spectrophotometry to monitor photoswitching and relaxation in bulk samples

Absorption spectra in cuvette ("UV-Vis") were acquired on a Varian CaryScan 60 (1 cm pathlength). For photoisomerisation measurements, Hellma microcuvettes (108-002-10-40) taking 500 $\mu$L volume to top of optical window were used with test solution such that the vertical pathlength of the isomerization light is less than 7 mm to the bottom of the cuvette, with the default test solution concentrations of 25 $\mu$M. Measurements on soluble photoswitches were performed by default in PBS at pH ~7.4 with 1% of DMSO to better mimic the intracellular environment during cell culture conditions (with 1% DMSO). Photoisomerisations and relaxation rate measurements were performed at room temperature. "Star" LEDs (H2A1-models spanning 360–590 nm from Roithner Lasertechnik) were used for photoisomerisations in the cuvette that were also predictive of what would be obtained in LED-illuminated cell culture.

The **AzTaxs** were not reliably soluble enough to be assayed in physiologically relevant aqueous media (~1% DMSO max, aqueous buffer) at ~50 $\mu$M as is necessary for long-term UV-Vis based studies on our setup. Therefore the spectra of the excellently water-soluble diethanolamide model photoswitches were instead acquired, in physiologically relevant aqueous media (PBS with <1% DMSO), to give the closest approximation of the PSSs to be expected in cell assays with the cognate series of **AzTax**s. Their absorption spectra at the photostationary states (PSSs) under illumination at different biocompatible and photoswitching-relevant wavelengths, were measured. Note that "dark" represents a solution quantitatively relaxed to all-*E* by warming overnight to 60 °C.

#### *Thermally reversible and photoreversible photoisomerisation*

Azobenzenes photoswitches featuring *para*-dialkylamino groups (**3DMA**, **3DEA**, **4DMA**) did not appear to undergo bulk photoisomerisation in homogeneous aqueous physiological media (1 cm UV-Vis cuvette measurement, 25 $\mu$M, PBS pH ~7.4, <1% DMSO, 37 °C, detection limit for photoisomerisation implies maintenance of PSS with ca. 2% *Z* isomer) which literature suggests is caused by fast (halflife < ms range) spontaneous ("thermal"), quantitative, unidirectional *Z*→*E* relaxation in this solvent[16]. All other azobenzenes were photoreversibly isomerisable in this homogeneous aqueous physiological media, which literature supports for azobenzenes not featuring strong resonance donor groups in *para* to the diazene[17]; results are shown for representative photoswitch **3MP** (Fig S1).

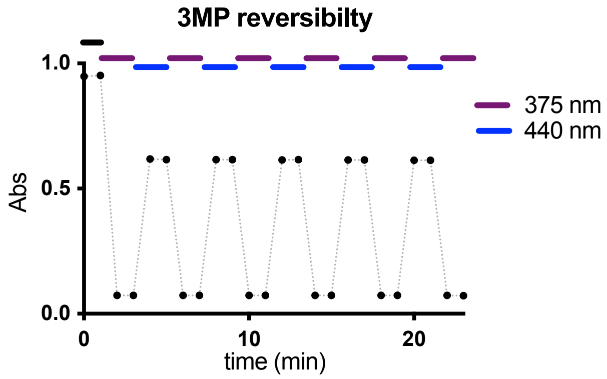

Figure S1: Photoisomerisations in homogeneous aqueous physiological media (PBS pH ~7.4, <1% DMSO, 37 °C) are perfectly photoreversible over many cycles, with no signs of degradation, implying robust and reproducible photoswitching can be possible under biological conditions.

We have previously observed however that non-azobenzene photoswitches that were not bulk-photoswitchable in homogeneous aqueous physiological media, can reliably display photoswitchability of bioactivity when used in the heterogeneous context of cell biology.[18] There are also reports of light-dependent activity for very fast-relaxing azobenzene photopharmaceuticals intended to address intracellular protein targets located in aqueous environments[19]; although, as far as we are aware, those fast-relaxing azobenzenes required illumination with such high photon flux to trigger irreversible bioactivity, that it is conceivable (given e.g. the mismatch between the *trans*-active structure-activity relationship expected, and the experimental *cis*-active result, as well as non-photoreversibility of biological effect) that transient photoisomerisation could to some extent be followed by glutathione (GSH) degradation of the more GSH-sensitive *cis*-azobenzene isomer[20], yielding a range of undefined, non-photoswitchable byproducts presumably including the diazene scission product aniline, several of which could be expected to be potent, photoirreversible, and biologically essentially irreversible enzyme-inhibiting species. In this work we therefore determined that biological evaluations for fast-relaxing **AzTax** conjugates would proceed with very limited photon flux, applied from short and low-intensity LED pulses, which we estimate to be insufficient[21–23] to give confounding results. Assuming that biological photoswitchability for these species can only arise by their biolocalisation in relatively hydrophobic environments (membranes, lipid vesicles, adsorbed onto proteins) which allow greater thermal stability of the metastable isomer, we therefore measured all photoproperties of the *para*-dialkylaminoazobenzenes in ethyl acetate solution, which we consider to be a reasonable mimic of an aprotic, moderately polar environment. We observed that this allowed **3DMA** to exhibit fully photoreversible isomerisations (Fig S2) which gave hope that *para*-dialkylaminoazobenzene **AzTax**s might prove to display photoswitchable bioactivity *in cellulo*.

We also monitored the rate of spontaneous ("thermal"), quantitative, unidirectional *Z*→*E* relaxation of all azobenzenes. The photoswitches used in this study could be split in two groups according to their performance as relevant to conditions for biological use: (1) The *para*dialkaylamino switches had *cis*-halflife $t_{1/2}$ ~ 11 min in EtOAc (although no switching was observed in water); whereas (2) *para*-alkoxy and *para*-unsubstituted azobenzenes showed relaxation that is much slower (in PBS with < 1% DMSO at 37 °C) than the typical 1-to-60 min timescale that biological assays would require for delivering functional reversibility; these would therefore require active *Z*→*E* photoisomerisation and/or diffusion-based reduction of localised *Z* isomer concentration in order to display biological reversibility. *Para*-alkoxy compounds displayed $t_{1/2}$ (halflife) values on the order of 10 h - 5 days (representative **3MP** had $t_{1/2} \sim$ 24 h); unsubstituted compounds displayed $t_{1/2}$ values substantially above 1 day (representative **4H** had $t_{1/2} \gg$24 h (estimated by exponential decay fit to be ca. 1200 h)). Note however that it is not important for this reagent development research to know precisely the values of the switches' halflives in homogeneous media *in cuvette*: they are however either far below, or else far above, the biological timescale and can be simply treated accordingly.

##### *Photostationary state (PSS) equilibria*

PSSs were measured. Results for three compounds representative of the three electronic classes of azobenzenes (**3MP** for *p*-OMe, **3DMA** for *p*-$NR_2$, **4H** for *p*-unsubstituted switches) are shown in Fig S2.

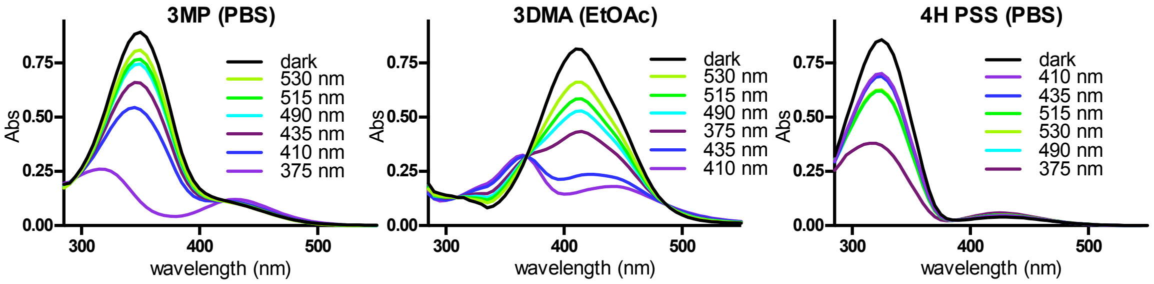

Figure S2: PSS spectra for photoswitches representative of the three structural classes explored in this work.

##### *PSS analysis*

For photopharmaceutical assays in biology it is helpful to anticipate the *E/Z* ratio at any wavelength's photostationary state (PSS), to choose optimal wavelengths for illumination during biological assays or to understand the limits of what is possible on a given setup: e.g. on a microscope with laser lines 405 nm, 488 nm and 515 nm, what dynamic range of photoswitchability is possible by establishing localised PSSs (inside a single cell) that alternate between 405 nm and 515 nm? Ideally, 405 nm would establish a PSS with 100% of one isomer (e.g. *Z* ), and 515 nm would establish a PSS with 100% of the other isomer, implying a 100% dynamic range of isomer photoswitchability; and ideally, one isomer (e.g. *Z*) would be bioactive while the other isomer would be entirely biologically inactive, therefore that 405 nm / 515 nm photoswitching would also allow 100% dynamic range of biological photoswitchability. However, no azobenzenes have ever been shown to enable 100% dynamic range of isomer photoswitchability, and anyway as far as we are aware only **PSTs**[23] feature one isomer that is entirely biologically inactive. Therefore it is impossible that any azobenzene-based photopharmaceutical according to current designs features 100% dynamic range of biological photoswitchability. Analysing PSS ratios *in cuvette* can extract the isomer photoswitchability, and analysing bioactivity determinations under those same PSSs in light of the isomer photoswitchabilities can determine the isomer *bioactivity differentials*; both analyses are needed to be able to gauge the dynamic range of biological photoswitchability that is theoretically obtainable under any arbitrary wavelength.

We have previously published a workflow[20] to estimate PSS at any wavelength in biological media, based on acquiring separated *E* and *Z* isomer spectra by LCMS-UV, and relying on isosbestic point determination in physiological media. In brief, to determine the ratios of *E* and *Z* isomers in PSS equilibria, the UV/Vis spectra of the HPLC-separated isomers are measured by inline DAD, extracted, scaled relative to each other using the isosbestic point determined from UV-Vis studies in biological media, then fitted as a linear combination to the measured PSS absorption spectra, with the linear combination coefficients then being the PSS fractions of each isomer.

However, in this research, the isolated E and Z spectra of the amides in LCMS eluent did not match up to their dark (all-*E*) and illuminated (mostly-*Z*) spectra as recorded in PBS buffer. The n→π* band notably showed lower intensity in the LCMS spectra which we attribute to influence of solvent, as the acidic acetonitrile/water eluent provides a different environment for the azobenzene than the neutral aqueous PBS buffer. We decided that results from measurements in PBS buffer however provide far more useful information for biology, than would performing PSS measurements in the HPLC solvent system, so we changed to a different "envelope" method that establishes entirely robust and assumption-free upper and lower bounds for the true PSS ratios. Since these bounds are often remarkably close to each other the method can allow remarkably precise, assumption-free estimation of the PSS isomer ratios in biological media, and also gives a maximum possible error of that PSS which it is useful to know.

<u>Envelope Method</u>

(0) The absorption spectrum of a fully-relaxed sample (60 °C overnight relaxation) is acquired and assumed to be the all-*E* spectrum. This step can also be checked by NMR (in contrast, we consider that establishing illuminated PSSs by NMR is not a straightforward approach due to (a) Lambert-Beer shielding at relatively high concentrations plus high reflection from the tube surface making establishment of many PSSs in many NMR tubes a very timeconsuming process; and (b) need for deuterated biological buffers).

(1) *Upper Bounds:* The *relative completeness* of photoreversions towards the all-*trans* state under different illuminations was examined first. For each PSS wavelength, the "relative completeness fraction" RCF(λ) calculated as the ratio [[P(λ)-MC(λ)]/[D(λ)-MC(λ)]], where P(λ) is the PSS absorption spectrum being evaluated, MC(λ) is the PSS absorption spectrum under the wavelength giving the most-*cis*-containing PSS, and D(λ) is the all-*trans* absorption spectrum, was calculated. This “relative completeness fraction” was calculated across the data range where the variation in absorbances with different PSSs is strongest (typically 370-420 nm), then the data were averaged to give the mean, and their standard deviation determined as a measure of the error in this fitting method. These completeness fraction

values are, by definition, lower bounds for the PSS values of *E*-content, so **[1-RCF(λ)] determines robust upper bounds** for the PSS values of *Z*-content.

(2) *Lower Bounds:* **Lower bounds** for the PSS *Z*-content values may separately be obtained by assuming the absorption of the *cis* isomer is zero at a single wavelength $\lambda_{strong}$ (the wavelength with the largest fold differential of extinction coefficients between *cis* and *trans* forms) and then tabulating $A(\lambda_{strong, PSS})/A(\lambda_{strong, all\text{-}trans})$. Typically, $\lambda_{strong}$ is approx. 385 nm.

(3) *Envelope:* The interval from lower to upper bound is an assumption-free bounded range for the true PSS at any measured wavelength. This will be seen to be more than sufficient to give a PSS range with typically only ±5-10% possible error at the wavelengths of most interest to this study. We here represent the envelope midpoint as the fitted "PSS %Z", and give the half-width of the envelope (100% CI) as the possible error "± %" (Table S1).

| λ (nm) | 3MP | | 4H | | 3DMA (in EA) | |
|---|---|---|---|---|---|---|
| | PSS %Z | +/- % | PSS %Z | +/- % | PSS %Z | +/- % |
| **375** | 96% | 4% | 80% | 20% | 53% | 5% |
| **410** | 44% | 2% | 26% | 6% | 91% | 9% |
| **435** | 29% | 1% | 28% | 7% | 80% | 8% |
| **490** | 18% | 1% | 39% | 10% | 39% | 4% |
| **515** | 16% | 1% | 39% | 9% | 32% | 3% |
| **530** | 11% | 1% | 38% | 9% | 21% | 2% |

Table S1 - Estimated PSSs from the envelope method and maximum error in the estimated PSSs, for the three different families of azobenzene photoswitches used in this study (*p*-OMe, *p*-unsubstituted, *p*-$NR_2$).

#### Part C: Biochemistry: tubulin polymerisation *in vitro*

99% tubulin from porcine brain was obtained from Cytoskeleton Inc. (cat. #T240). The polymerisation reaction was performed at 5 mg/mL tubulin, in polymerisation buffer BRB80 (80 mM piperazine-N,N′-bis(2-ethanesulfonic acid) (PIPES) pH = 6.9; 0.5 mM EGTA; 2 mM $MgCl_2$), in a cuvette (120 µL final volume, 1 cm path length) in a Varian CaryScan 60 with Peltier cell temperature control unit maintained at 37 °C; with glycerol (10 µL). Tubulin was incubated at 37 °C with “pre-lit”- [360 nm-pre-illuminated; mostly-*Z*-] or dark- [all-*E*] **AzTax3MP**, or docetaxel (final inhibitor concentration 10 µM), or without inhibitor ("cosolvent" control), in buffer with 3% DMSO and 1 mM GTP, and the change in absorbance at 340 nm was monitored, scanning at 15 s intervals[24]. Docetaxel showed the strongest microtubule hyperpolymerisation effect; pre-lit **AzTax3MP** had ca. 2/3 of docetaxel's hyperpolymerising potency compared to cosolvent-only control; all-*E* **AzTax3MP** had had ca. 1/3 of docetaxel's potency (Fig 2d).

### Part D: Cell Biology

#### *Cell assay methods*

##### General cell culture

HeLa cells were maintained under standard cell culture conditions in Dulbecco's modified Eagle's medium (DMEM; PAN-Biotech: P04-035550) supplemented with 10% fetal calf serum (FCS), 100 U/mL penicillin and 100 U/mL streptomycin. Cells were grown and incubated at 37 °C in a 5% $CO_2$ atmosphere. Cells were cultured in phenol red free medium prior to assays (DMEM; PAN-Biotech: P04-03591). Compounds and cosolvent (DMSO; 1% final concentration) were added *via* a D300e digital dispenser (Tecan); all photoswitches were added in their all-*E* state (thermal relaxation of the DMSO stocks at 60 °C overnight, applied under light exclusion conditions). Cells were either incubated under "lit" or "dark" conditions; "lit" indicates a pulsed illumination protocol applied by multi-LED arrays to create, *in situ* in cells, the wavelength-dependent PSS isomer ratio of the compounds, and then maintain it throughout the experiment, as described previously.[20,22] Typical "lit" timing conditions were 75 ms pulses applied every 15 s. "Dark" indicates that compounds were applied while working, sterile, under red-light conditions, and cells were then incubated in light-proof boxes to shield from ambient light, thereby maintaining the all-*E*-isomer population throughout the experiment.

##### Resazurin antiproliferation assay

Cells were seeded in 96-well plates at 5,000 cells/well and left to adhere for 24 h before treating with various concentrations of different compounds. *E*-**AzTax** were added and incubated under the indicated lighting conditions for 48 h (final well volume 100 $\mu$L, 1% DMSO; three technical replicates); the "cosolvent control" ("ctrl") indicates treatment with DMSO only. Cell viability was measured by addition of resazurin, which is reduced to resorufin under metabolic activity in live cells. Fluorescence of the resorufin product was measured using a FLUOstar Omega microplate reader (BMG Labtech) at 544/590 nm (ex/em). Fluorescence data was averaged over technical replicates, then normalized to viable cell count from cosolvent control cells (%control) as 100%, where 0% viability was assumed to correspond to zero. Three independent experiments were performed and data is shown as mean±SD; data were plotted against the log of **AzTax** concentration ($\log_{10}$([**AzTax**]) (M)).

##### Cell cycle analysis

HeLas were seeded in 6 well plates (300,000/well) 24 h prior to treatment. **AzTax3MP** and **AzTax4DMA** were added to the wells and cells were incubated either under "dark" or "lit" regimens. 0.1 $\mu$M Docetaxel served as positive control and 1% DMSO as cosolvent control. Cells were harvested 24 h later and fixed overnight in 70% ice cold ethanol. After 12 h, cells were washed and re-hydrated for 15 min in PBS before staining with propidium iodide ("PI", 200 $\mu$g/mL in 0.1 % Triton X-100 containing 200$\mu$g/mL DNase-free RNase (Thermo Fischer Scientific EN0531) for 30 min at RT. Flow cytometry was done with an LSR Fortessa (BD Biosciences) run by BD FACSDiva 8.0.1 software and at least 10,000 individual PI-positive cells per condition were collected. FlowJo software (BD Biosciences) was used for gating, first selecting alive cells, then single cells and then setting gates in the PI channel that correspond to less than two sets of chromosomes (subG1), two sets of chromosomes (G1), more than two and less than four (S) and four sets of chromosomes (G2/M). Results plotted as % of parent gate and are given as the mean±SD of at least three biological replicates.

**Immunofluorescence staining**

For visualization of polymerized tubulin and DNA cells were seeded on glass coverslips in 24 well plates (50,000 cells/well) 24 h prior to treatment. **AzTax3MP**, DMSO or 0.1 µM docetaxel was applied the next day (concentration range **AzTax3MP**: 0.1 µM-3 µM, all wells with 1% DMSO) and cells were incubated either in the dark or with the regular illumination protocol. The next day medium was removed, Cells were washed with pre-warmed (37 °C) MTSB buffer (80 mM PIPES, pH 6.8; 1 mM $MgCl_2$, 5 mM ethylene glycol tetraacetic acid (EGTA) dipotassium salt; 0.5% Triton X-100) for 30 s to remove tubulin monomers then fixed with 0.5% glutaraldehyde for 10 min. After quenching with 0.1% $NaBH_4$ cells were blocked for 30 min in PBS containing 30% FCS before incubation with anti-α-tubulin primary antibody (1:400 rabbit Abcam ab18251) for 1 h. Secondary antibody was donkey-anti-rabbit Alexa488 (Thermo Fisher Scientific A21206; 1:400 in PBS + 10% FCS). Coverslips were then mounted on slides with Roti-Mount FluorCare DAPI (Carl Roth) and left to dry. Confocal images were acquired on a Leica SP8 with a 405 nm laser and a white light laser, using a 63× glycerol objective. Confocal stacks (0.33 µm step size) were z-projected and gamma adjusted for better visualization in Fiji/ImageJ.

##### *Resazurin viability assay results for all compounds*

Results for all compounds are shown in Fig S3. The results of the resazurin assays can be correlated with the compounds' structures. The important parameters are the general potency of the compound (roughly, the average $IC_{50}$ of the lit and dark states) and the dynamic range (difference of $IC_{50}$ dark vs lit).

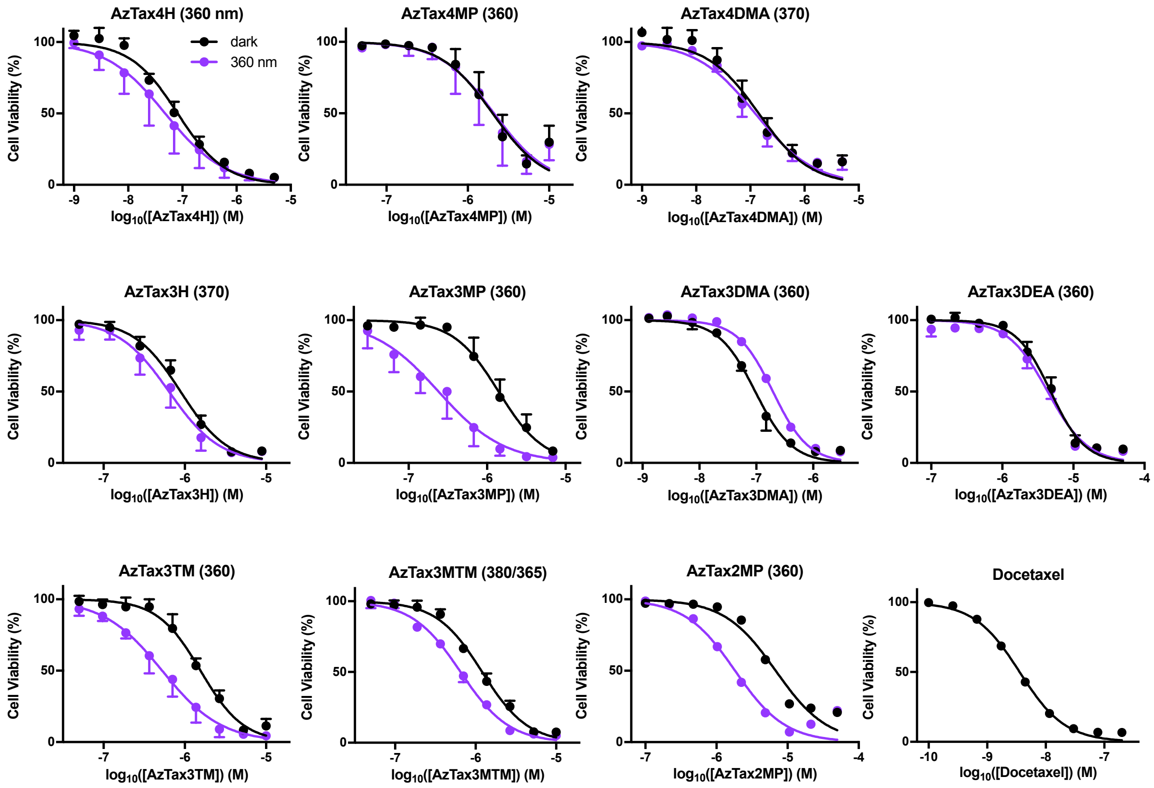

Figure S3: resazurin viability assay results for all compounds. The wavelength/s (in nm) used for the "lit" experiments (shown as purple curves) are given in brackets after each graph title, as different wavelengths were used for the assays. The "dark" experiments are depicted in black.

The first structural element to be examined is the attachment point of the azobenzene to the taxane scaffold. The number in the compound name specifies the attachment relative to the diazene bridge (2 = *ortho*, 3 = *meta*, 4 = *para*). It can be generalized that no *para* connected compound showed significant lit vs. dark difference of $IC_{50}$. *Meta* connected compounds show the highest dynamic range, and attachment in *ortho* reduced the overall potency significantly.

The second structural element examined is the substitution on the azobenzenes. Unsubstituted compounds **AzTax4H** and **AzTax3H** show no significant toxicity change upon illumination, although **AzTax4H** is an order of magnitude more toxic than **AzTax3H**. Alkylated *para*-amino compounds with fast relaxation times also show no strong difference between dark and lit experiments; **AzTax4DMA** and **AzTax3DMA** are approximately equally toxic and **AzTax3DMA** is the only compound that appears to show a higher toxicity under dark conditions, while more polar **AzTax3DEA** shows substantially lower toxicity than either dimethylamino compound. The last group of **AzTax** compounds are variously methoxylated. **AzTax4MP** shows no difference in $IC_{50}$ upon irradiation. The meta connected **AzTax3MP** shows the highest dynamic range as well as satisfactory toxicity. The two derivatives

**AzTax3TM** and **AzTax3MTM** have roughly the same cytotoxicities but more moderate dynamic range. **AzTax2MP** shows a drop in toxicity while displaying higher toxicity under illuminated conditions.

#### *FACS cell cycle analysis*

Results of cell cycle analysis for **AzTax3MP** were shown in Fig 3b-c. Docetaxel and DMSO and lighting controls are shown in Fig S4a; results for non-photoswitchable yet cytotoxic control compound **AzTax4DMA** are shown in Fig S4b; and gating strategy is depicted in Fig S4c. Lighting and cosolvent cause no change to cell cycle repartition; **AzTax4DMA** (whose short aqueous *cis*-halflife should prevent any light-dependent bioactivity being visible) shows no light-dependent effects, and it also dose-dependently recapitulates the cell cycle repartition seen for positive control docetaxel.

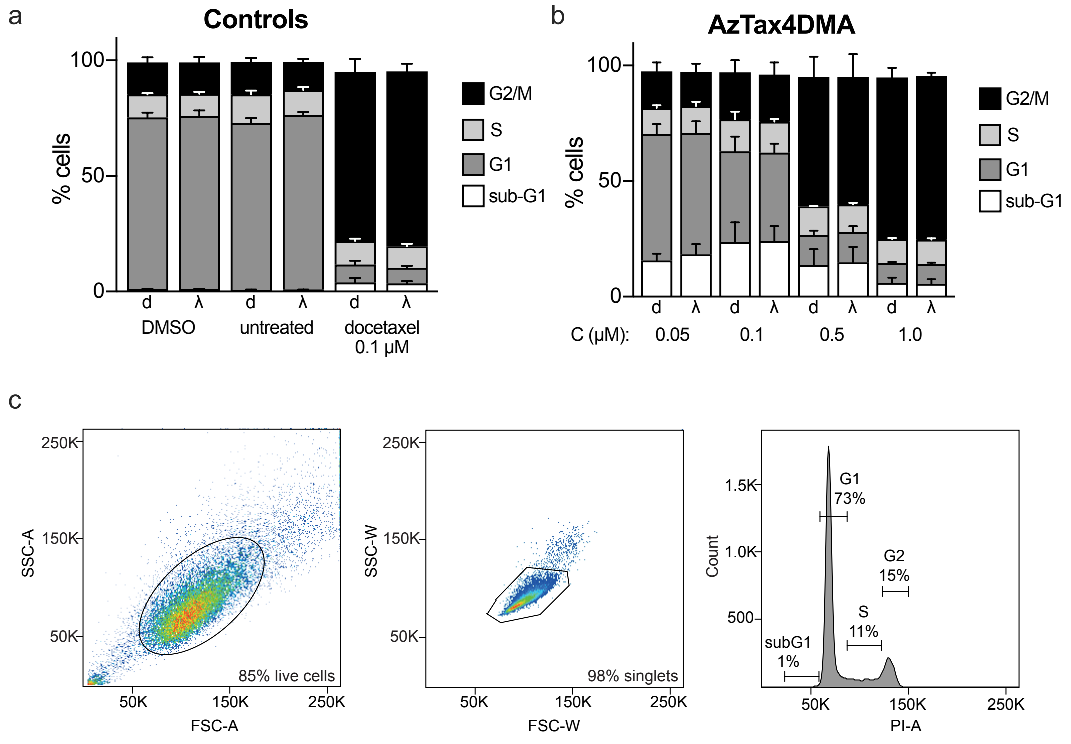

Figure S4: Controls for cell cycle repartition. **a** Controls without ("untreated") and with ("DMSO") cosolvent show no cosolvent-induced change of cell cycle repartition; positive control docetaxel gives strong G2/M arrest; and none of these controls show light-dependency of cell cycle repartition. **b** Non-photoswitchable **AzTax4DMA** shows no light-dependency of cell cycle repartition, but does show dose-dependent G2/M arrest (compare Fig 3c). **c** Gating strategy for the FACS cell cycle analyses.

#### *Immunofluorescence imaging of microtubule network structure*

**AzTax3MP** caused light- and dose-dependent disruption of microtubule structure, as well as cell toxicity (visible as the density of cells) (Fig 3a, Fig S5). At 0.1 $\mu$M, disorganisation of the MT network is evident under lit conditions but no change is seen under dark conditions. By 0.5 $\mu$M, **AzTax3MP** causes extensive mitotic arrest under lit, but not dark, conditions. At 1 $\mu$M, under lit conditions only a minor population of adherent cells persists (although the latter have severe MT organisational defects and multinucleation) and mitotic spindle defects are visible (see arrows and inset), while under dark conditions cells mainly escape mitotic arrest (albeit with nuclear defects) and no mitotic arrests are visible. Nuclei are especially fragmented and highly condensed under lit treatment. Fantastically disrupted microtubule patterns are seen with higher doses of lit **AzTax3MP** and some cells appear to have no remaining microtubule structures, perhaps since non-microtubular tubulin aggregates (expected at high doses of MT stabiliser, as seen in the cell-free polymerisation assay) are removed during the wash steps of the staining process; under dark conditions, mitotic arrests are seen and nuclear disorganisation becomes severe although the MT networks persist, in a disorganised state.

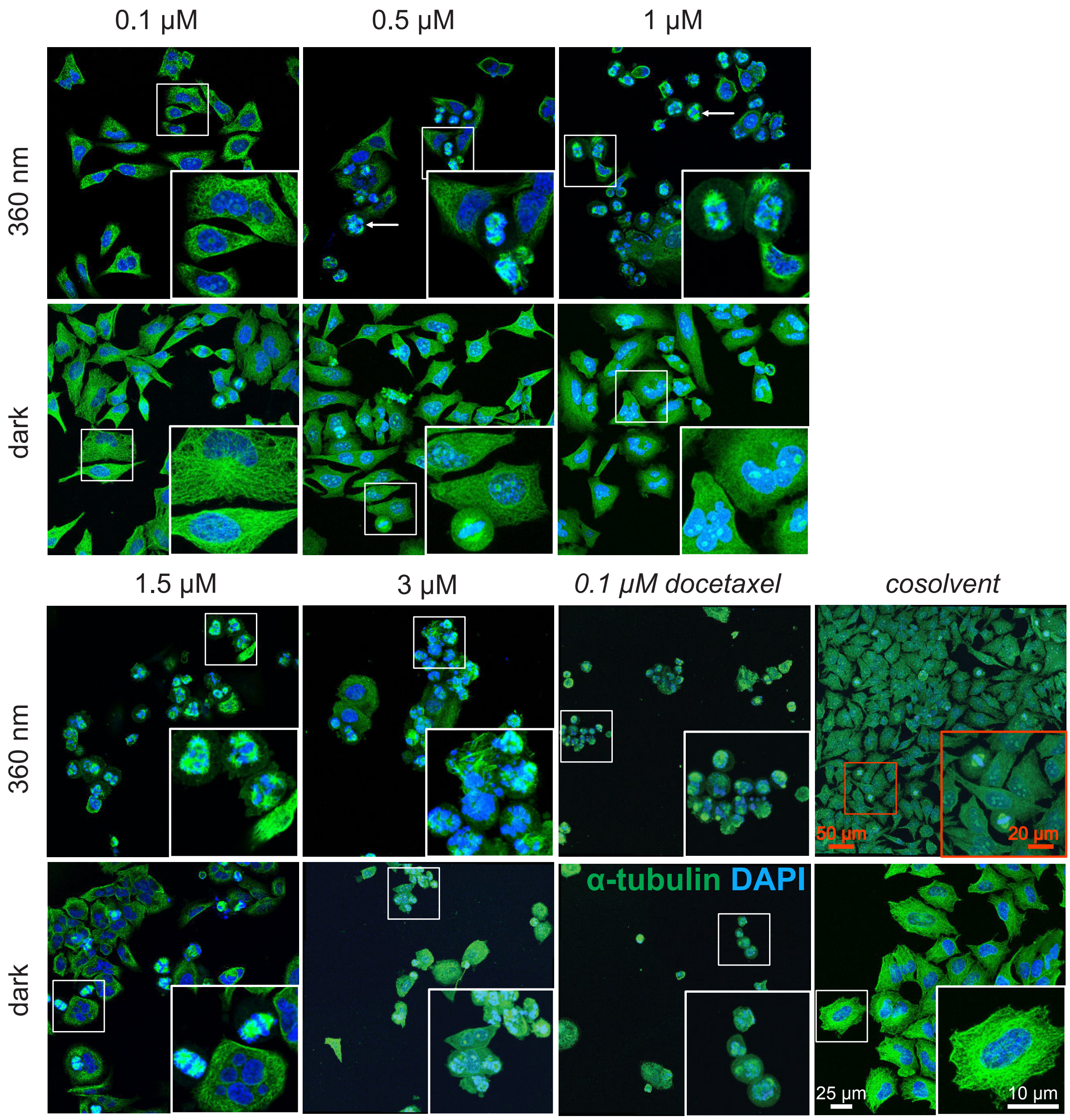

Figure S5 (expanded from data shown in Fig 3a): Immunofluorescence imaging of cells treated under lit/dark conditions with **AzTax3MP**, docetaxel (positive control), or 1% DMSO only (negative control) (Hela cells, MTs immunostained with anti-$\alpha$-tubulin (green), nuclei stained with DAPI (blue), 25 $\mu$m scale for overviews, 10 $\mu$m scale for insets (white) except for the orange-indicated cosolvent control panel (50 $\mu$m / 20 $\mu$m for better overview).

#### Supporting Information Bibliography

### Part F: NMR Spectra

#### methyl 4-((4-(dimethylamino)phenyl)diazenyl)benzoate (S1) : [1]H-NMR

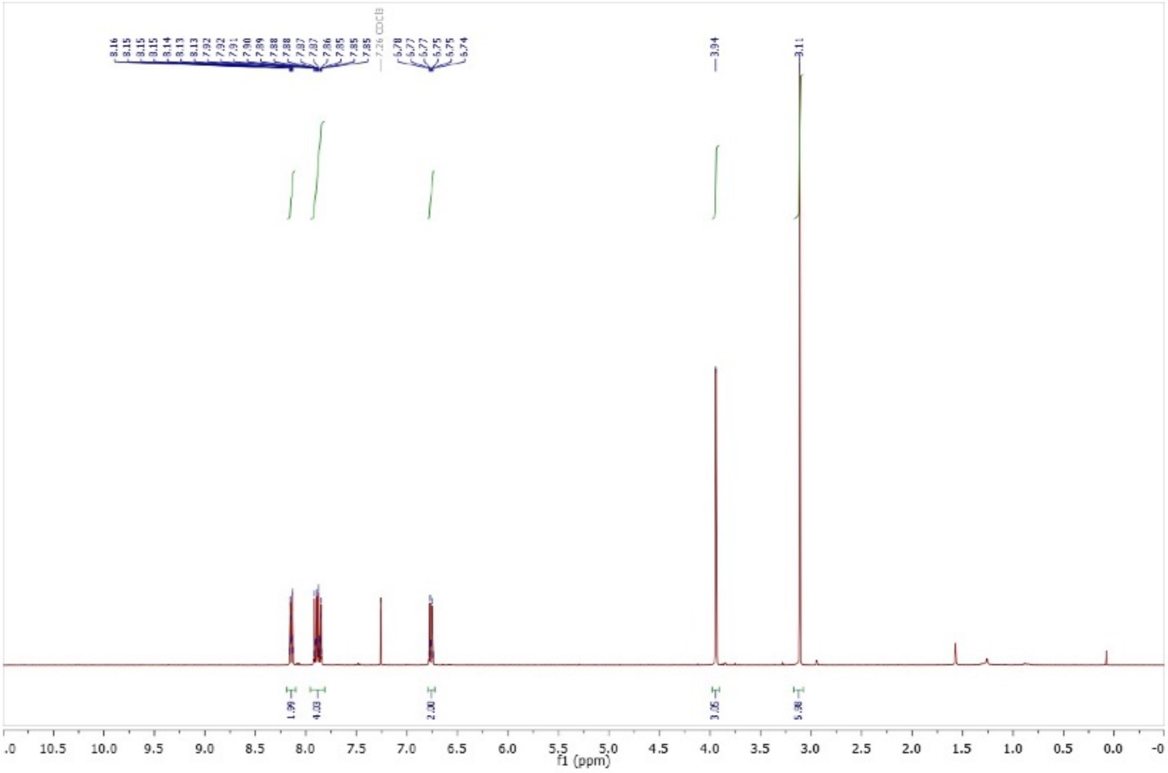

#### [13]C-NMR

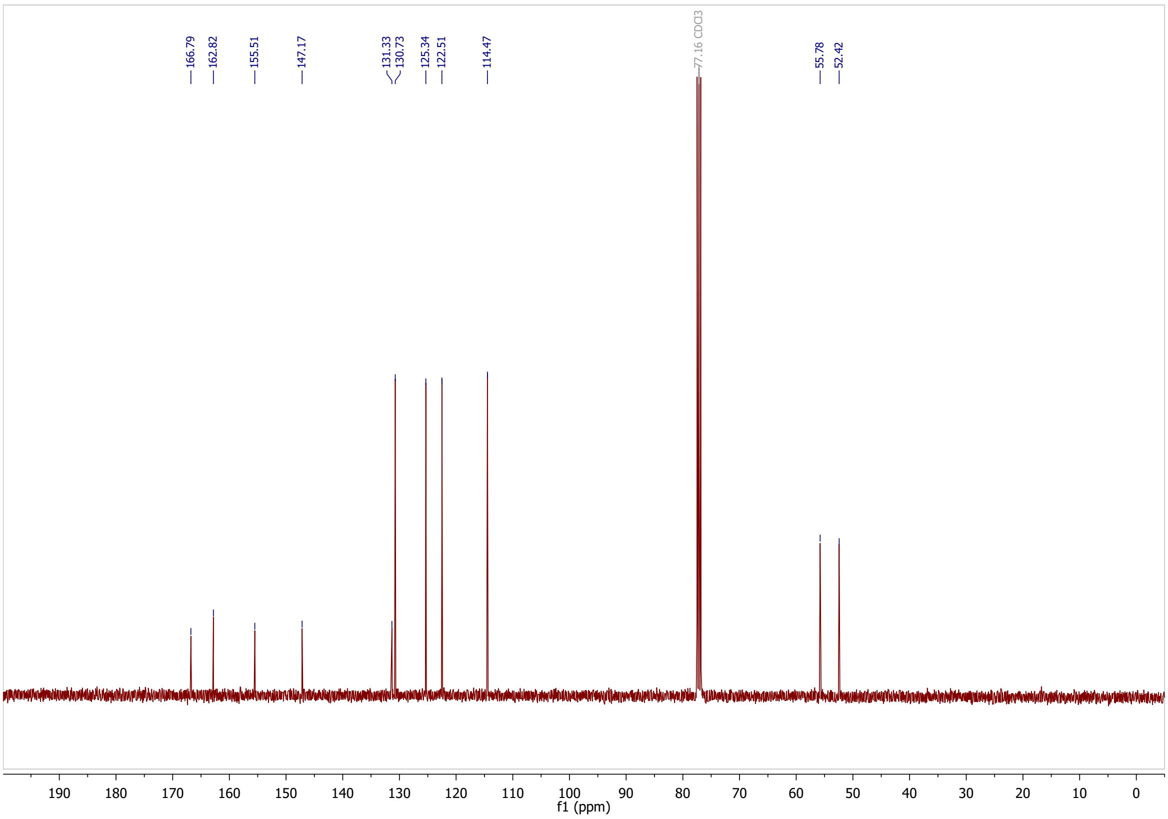

#### 4-((4-(dimethylamino)phenyl)diazenyl)benzoic acid (4DMA-$CO_2$H): $^1$H-NMR

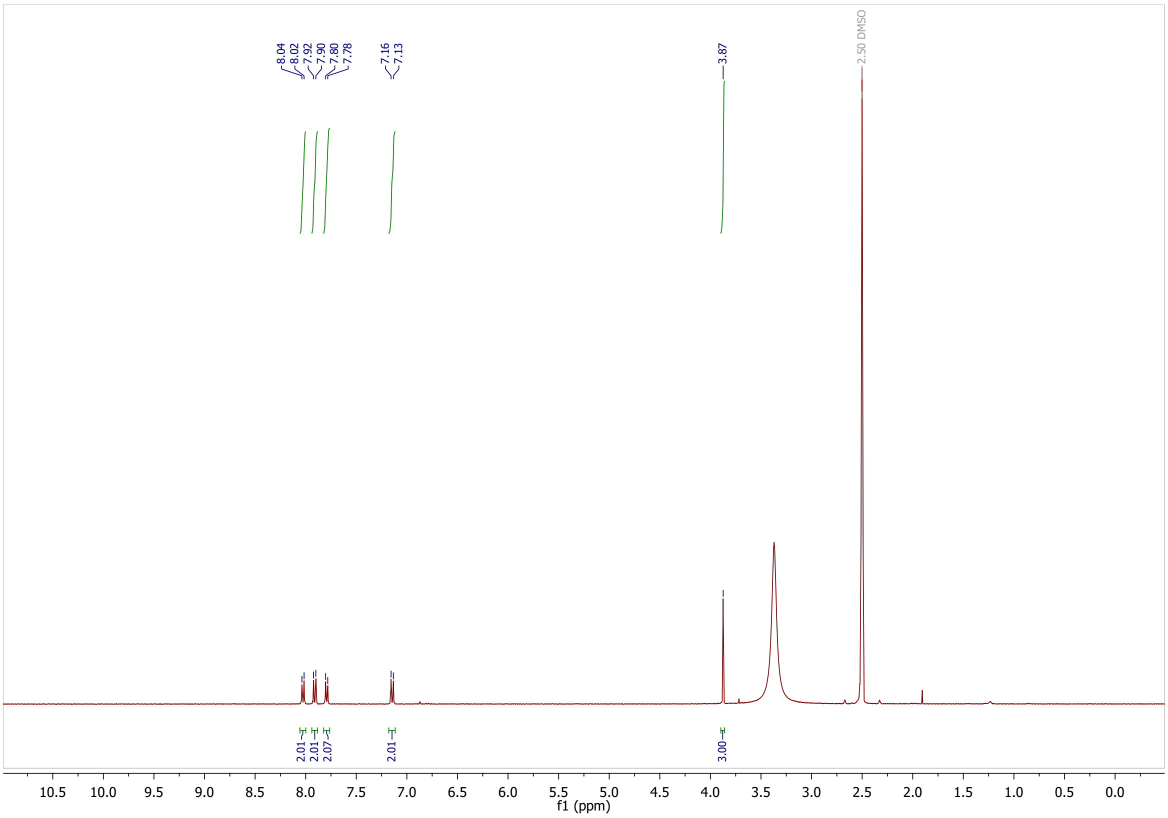

#### $^{13}$C-NMR

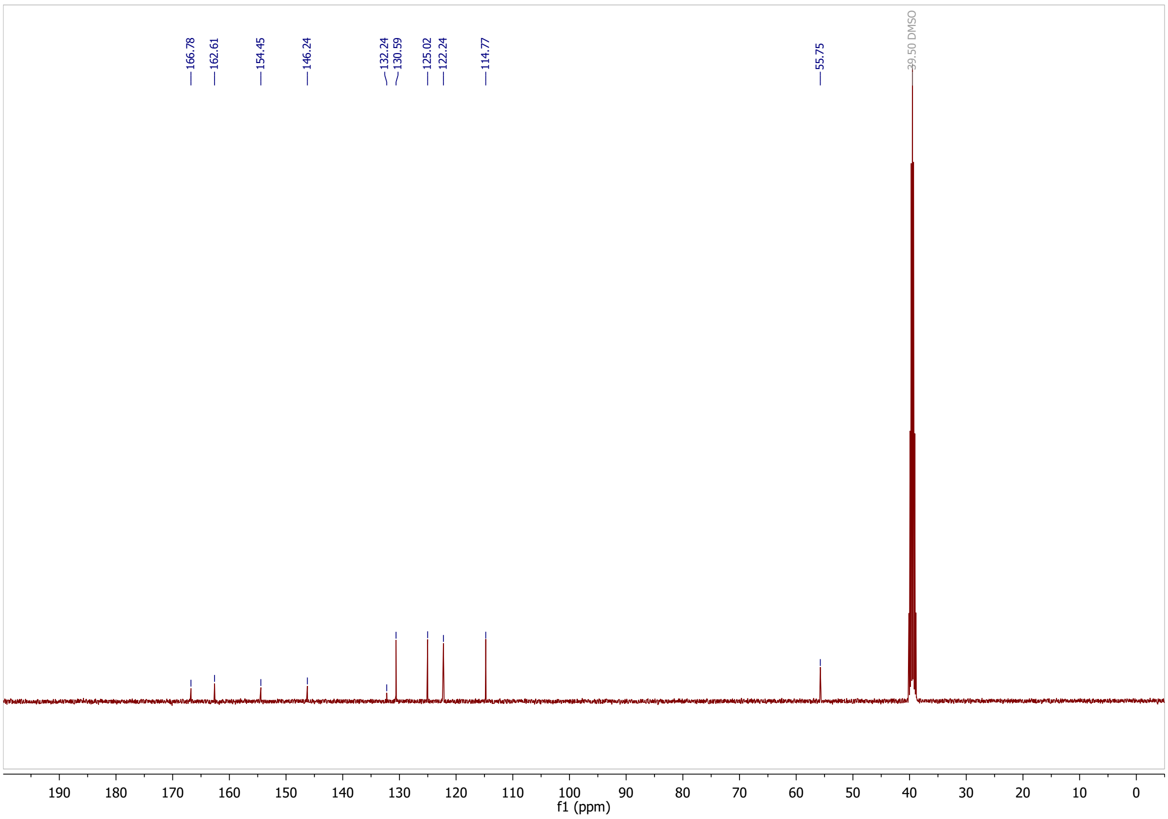

**methyl 4-((4-hydroxyphenyl)diazenyl)benzoate (S2) [1]H-NMR**

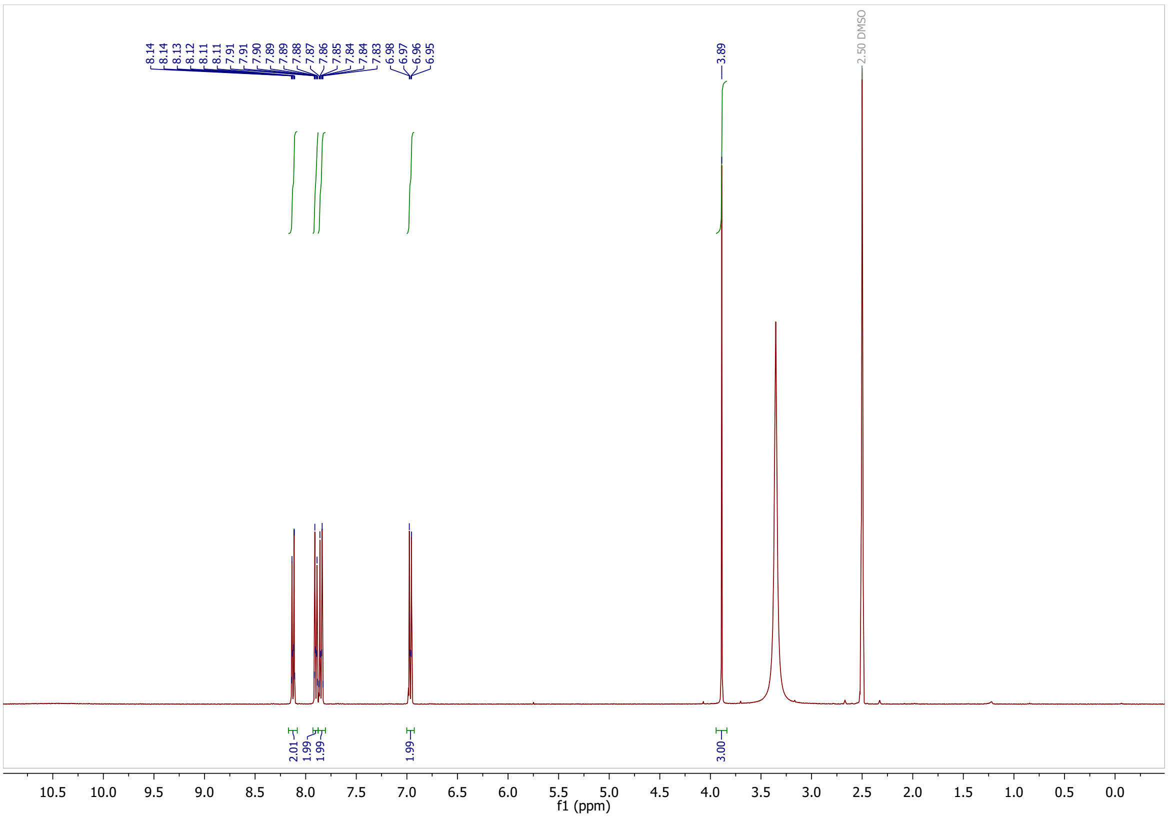

**[13]C-NMR**

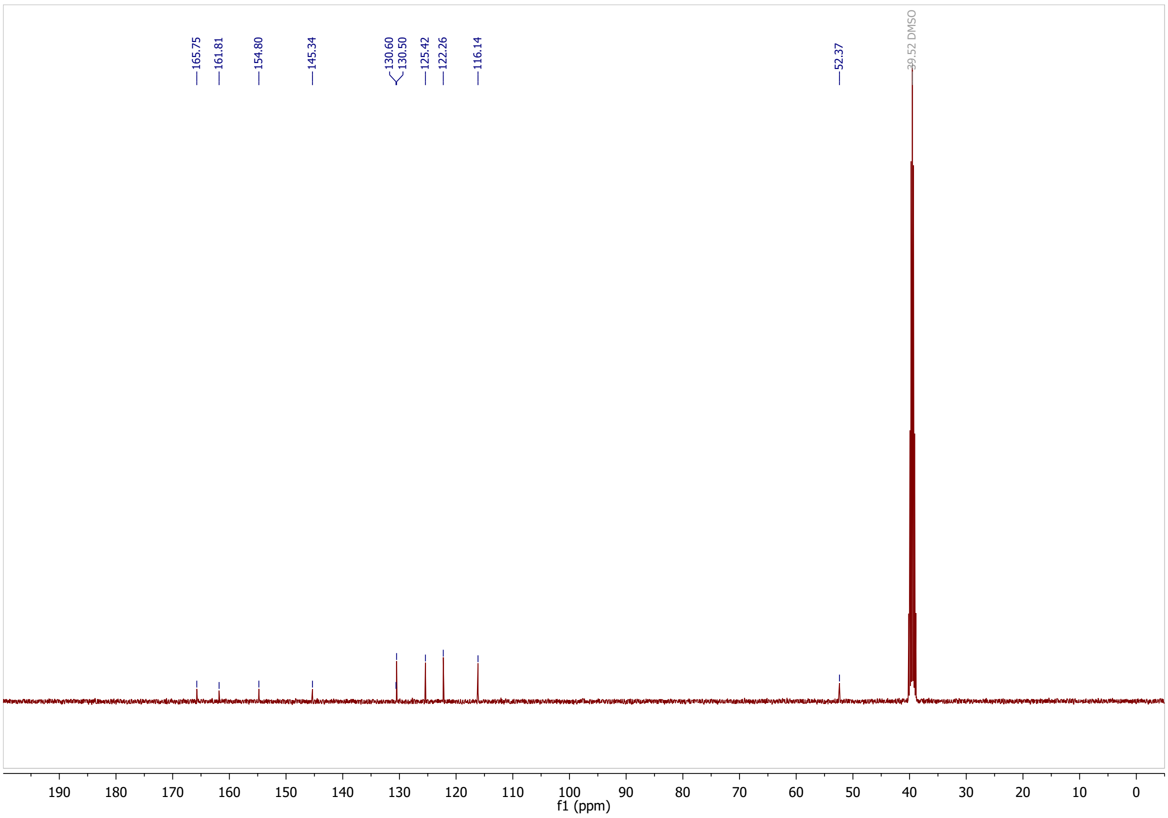

**methyl 4-((4-methoxyphenyl)diazenyl)benzoate (S3) $^1$H-NMR**

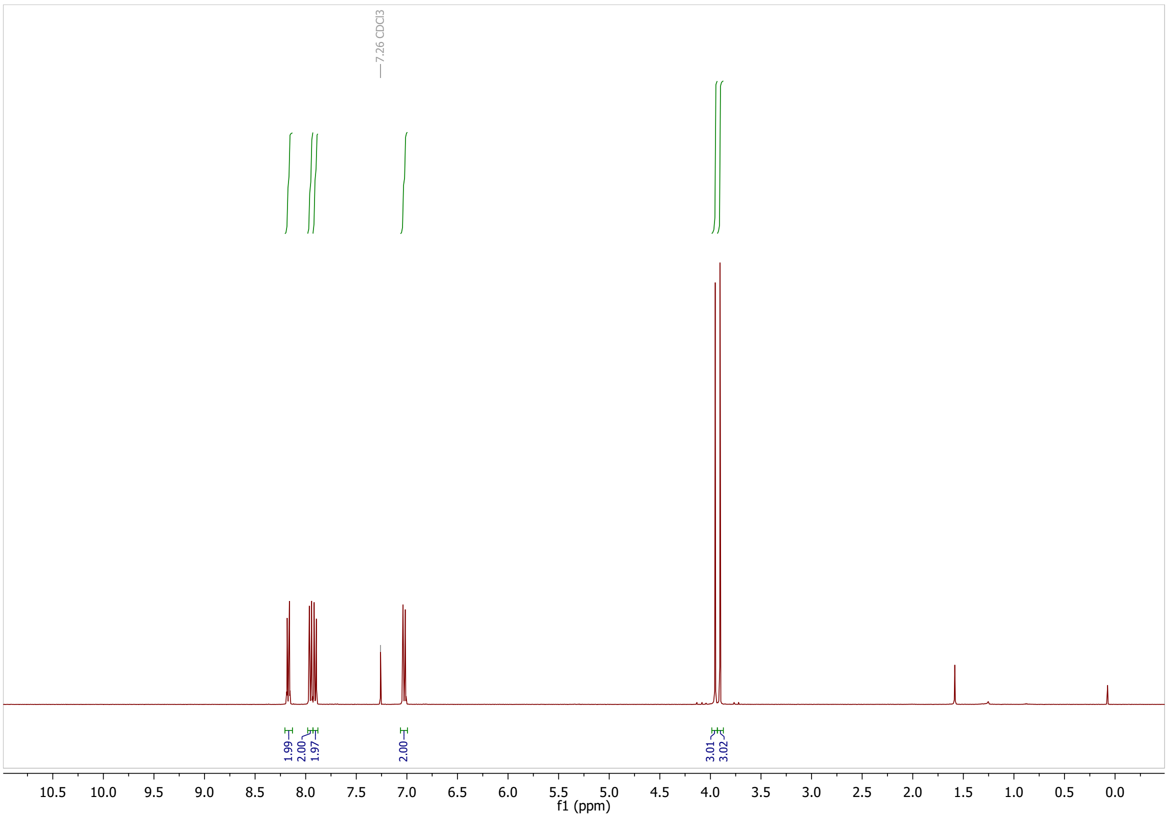

**$^{13}$C-NMR**

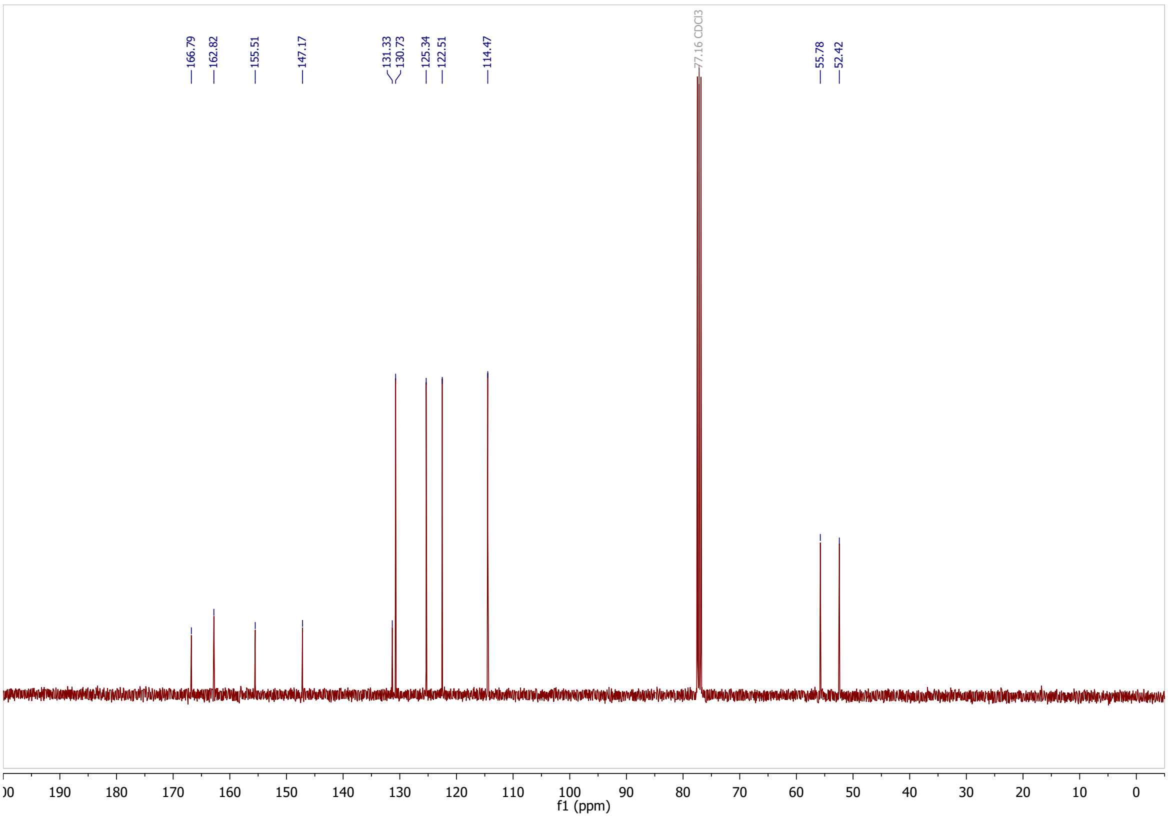

#### 4-((4-methoxyphenyl)diazenyl)benzoic acid (4MP-$CO_2$H): $^1$H-NMR

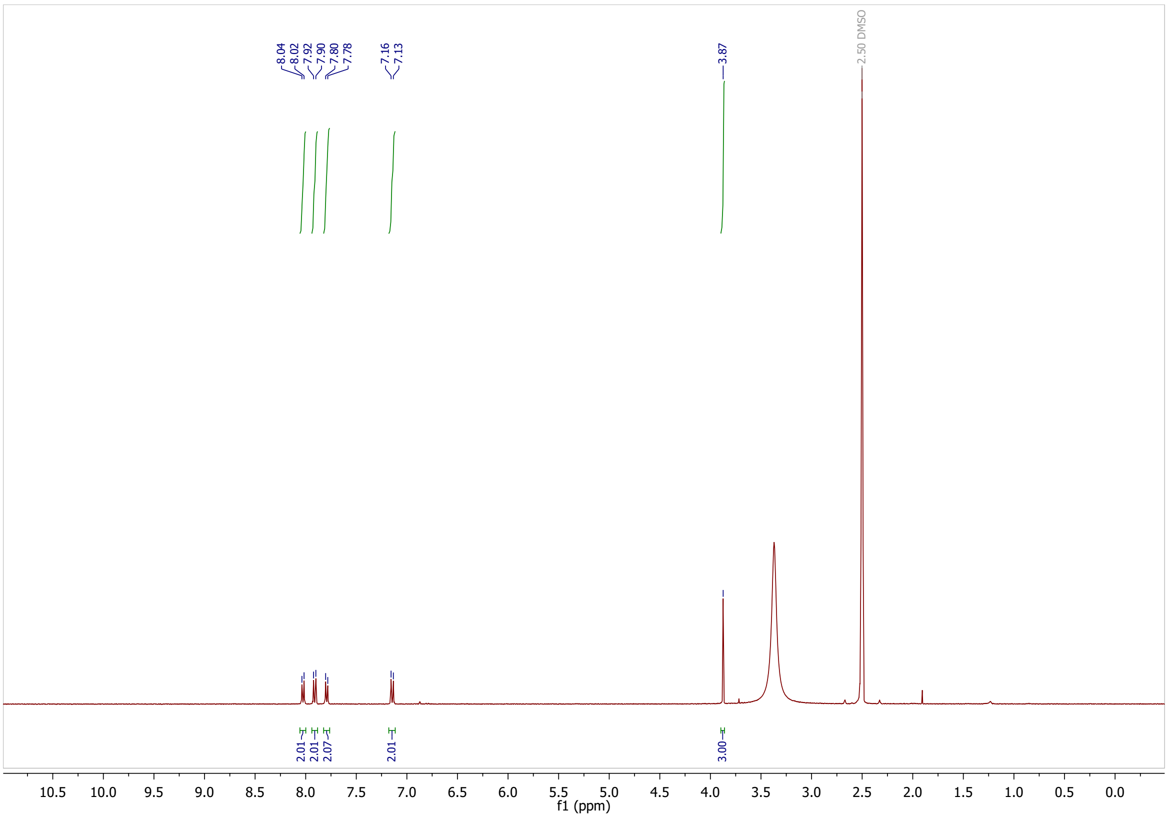

#### $^{13}$C-NMR

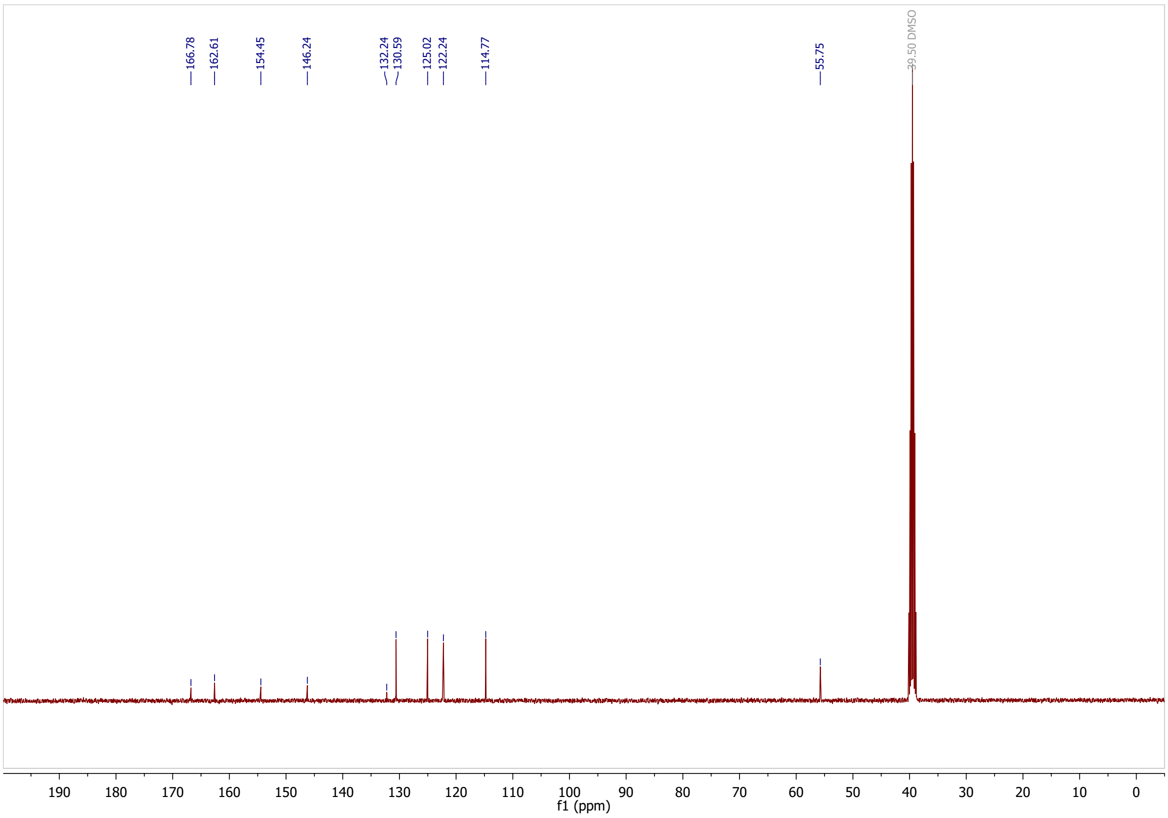

#### 3-(phenyldiazenyl)benzoic acid (3H-CO$_2$H): $^1$H-NMR

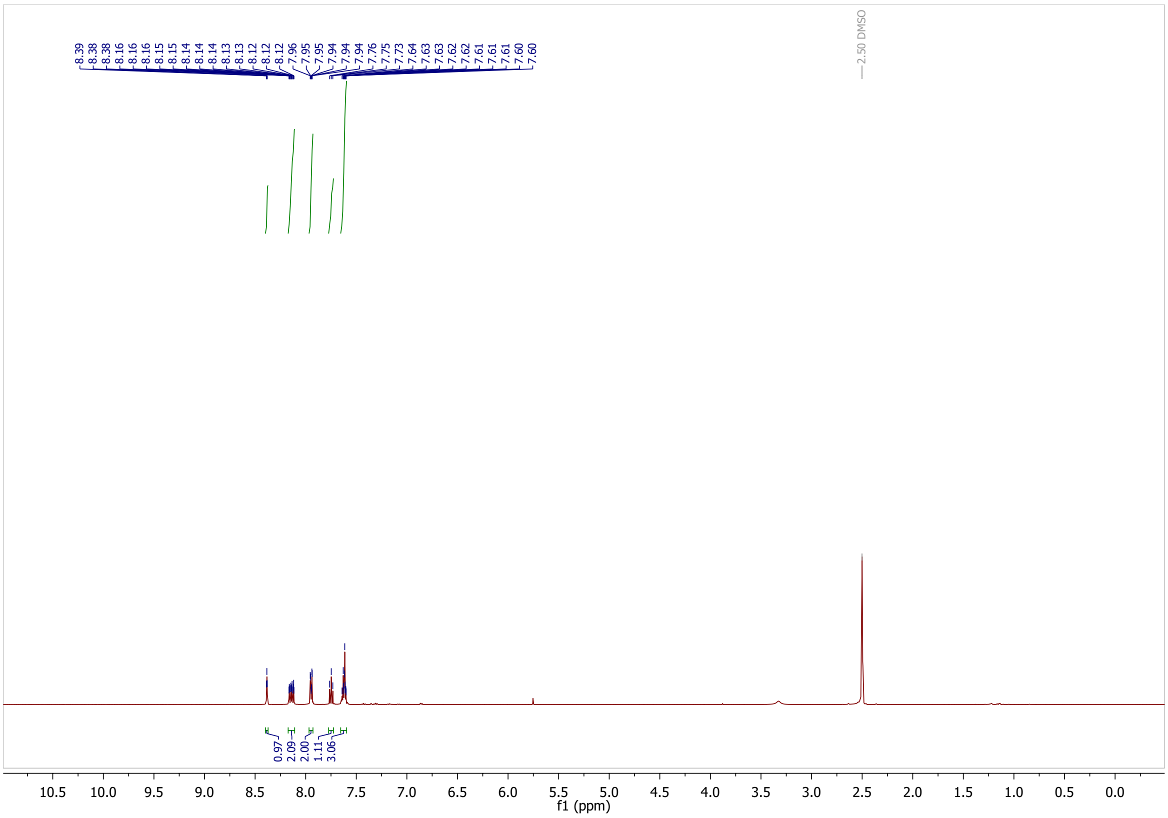

#### $^{13}$C-NMR

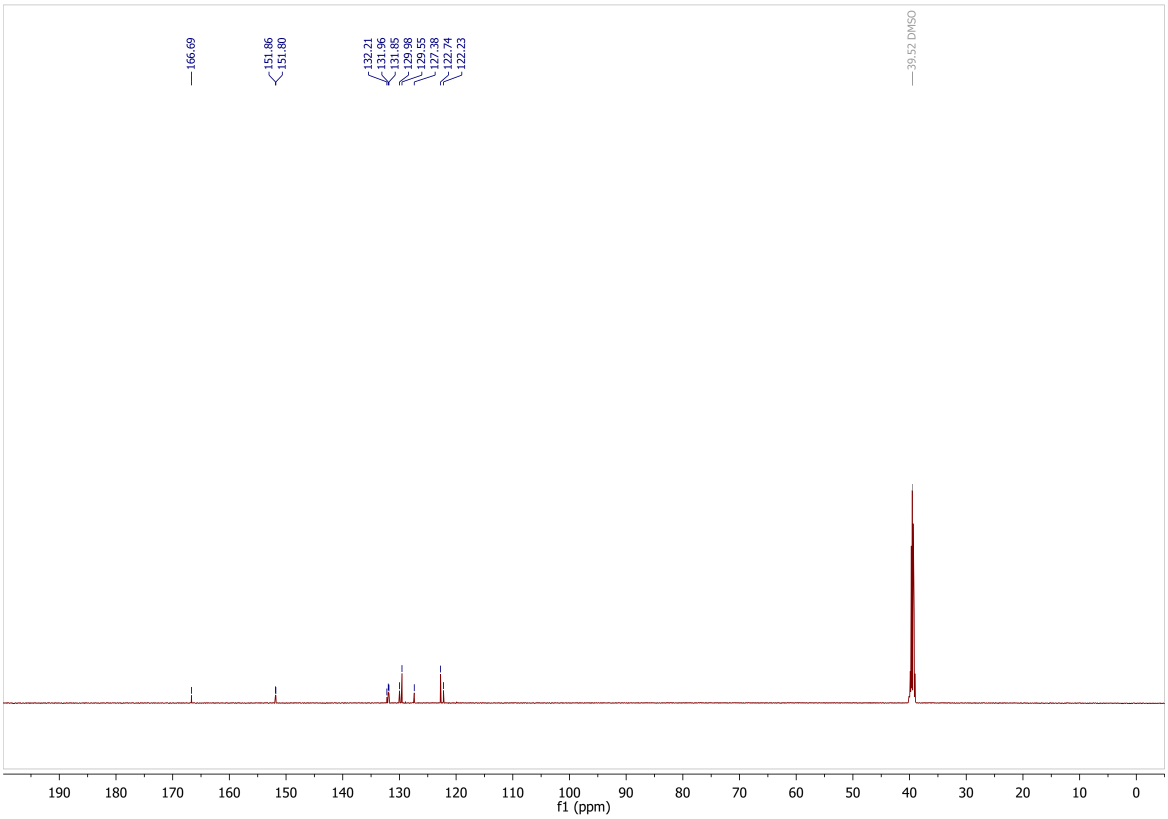

**methyl 3-((4-hydroxyphenyl)diazenyl)benzoate (S4): $^1$H-NMR**

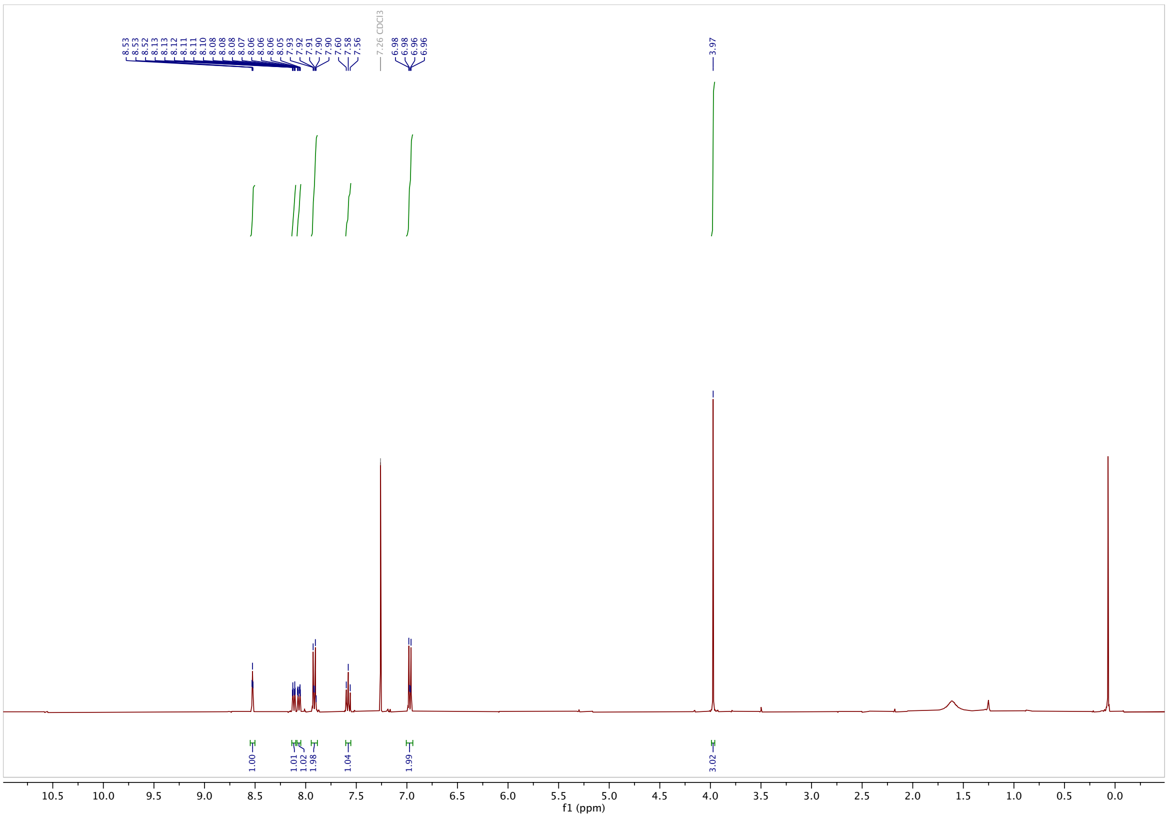

**$^{13}$C-NMR**

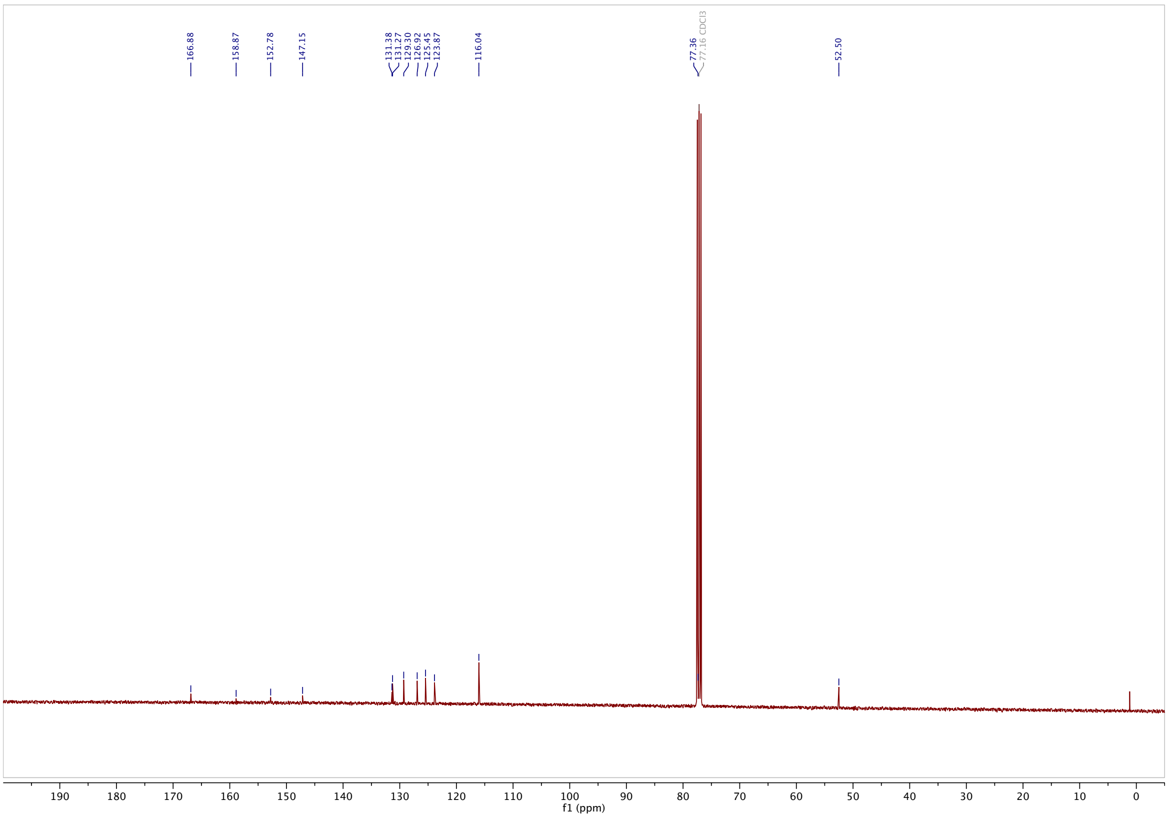

**methyl 3-((4-methoxyphenyl)diazenyl)benzoate (S5): $^{1}$H-NMR**

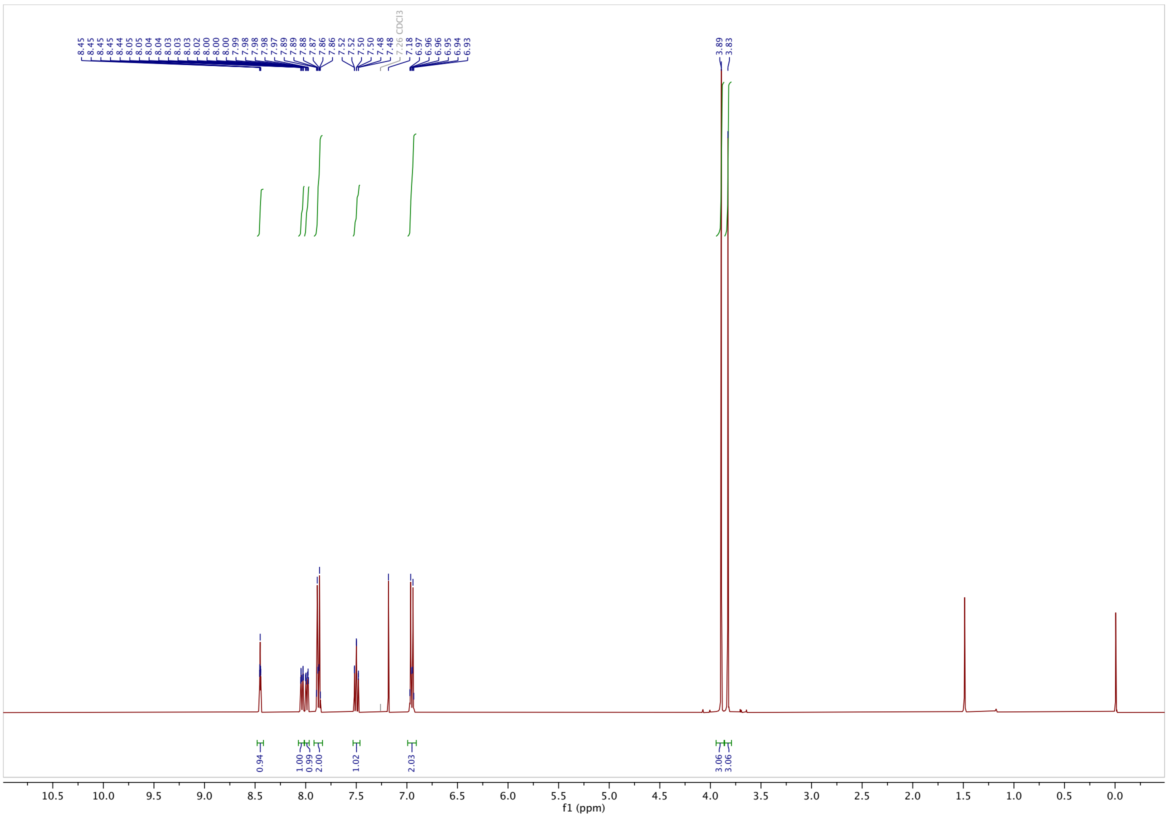

**$^{13}$C-NMR**

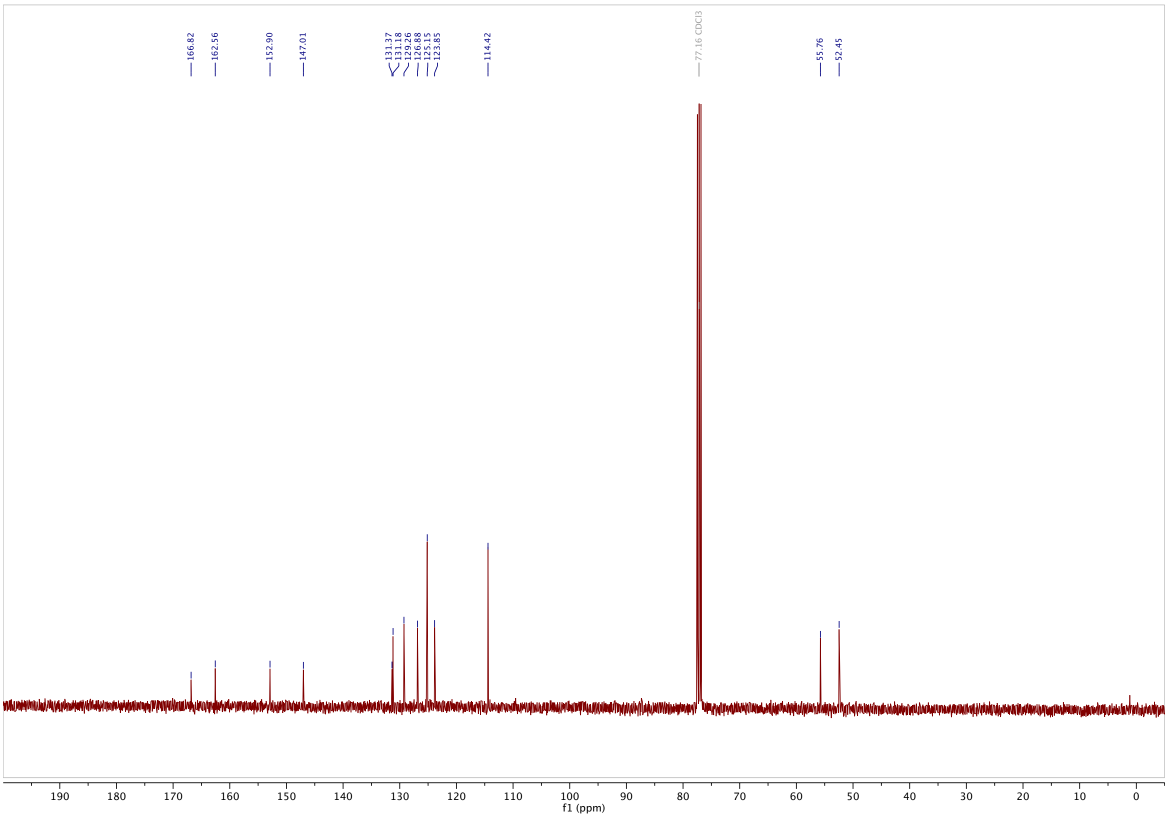

#### 3-((4-methoxyphenyl)diazenyl)benzoic acid (3MP-$CO_2$H): $^1$H-NMR

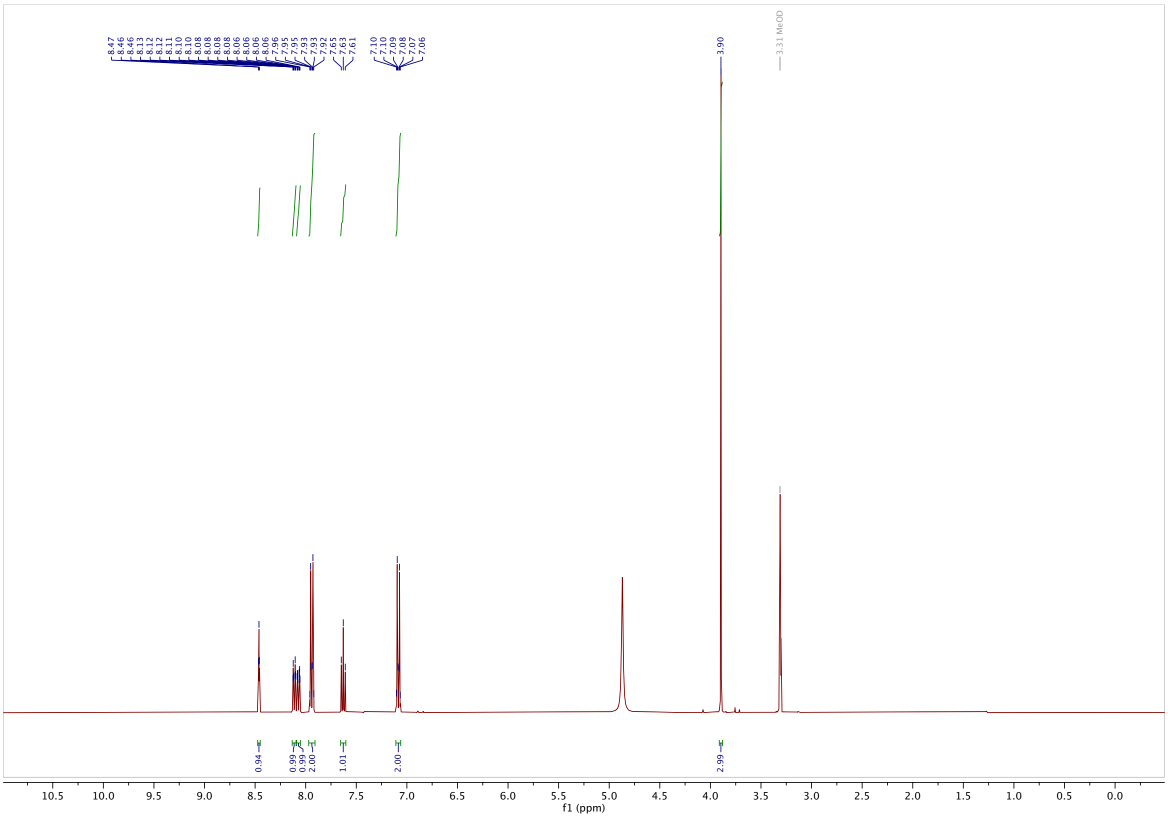

#### $^{13}$C-NMR

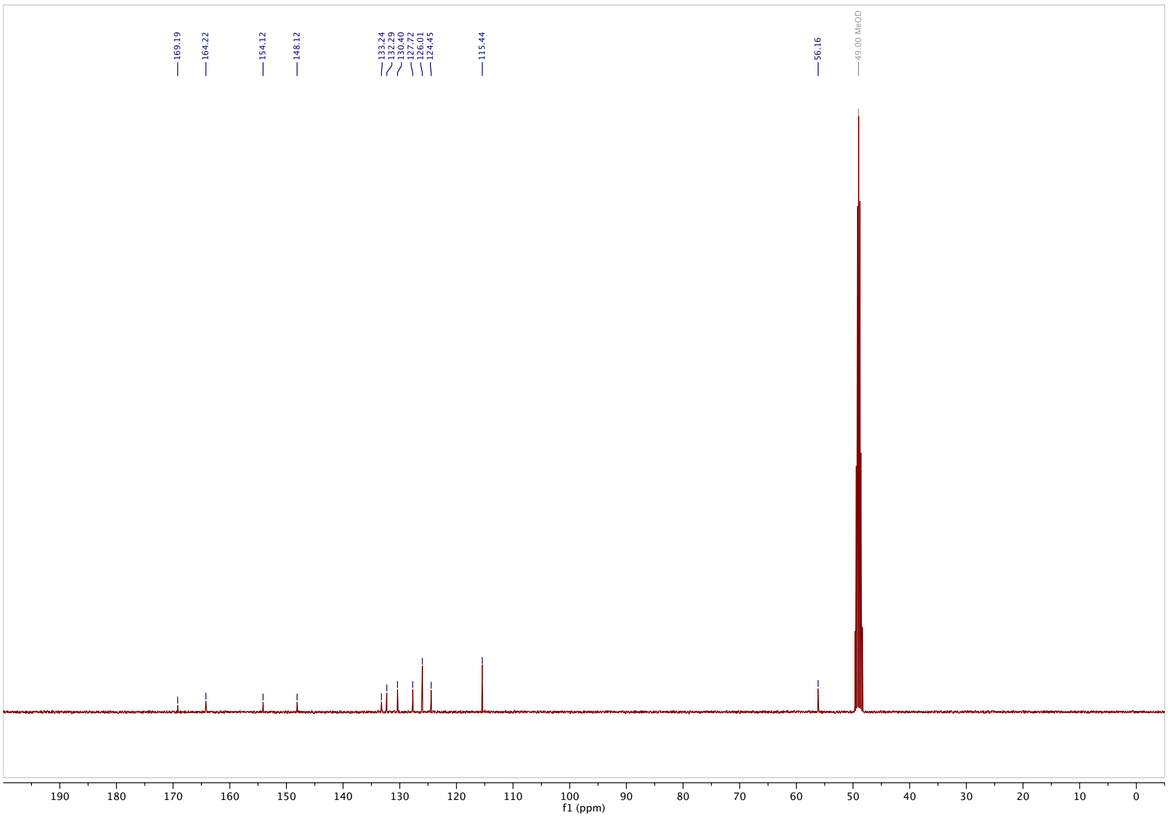

**methyl 3-((4-(dimethylamino)phenyl)diazenyl)benzoate (S6): ¹H-NMR**

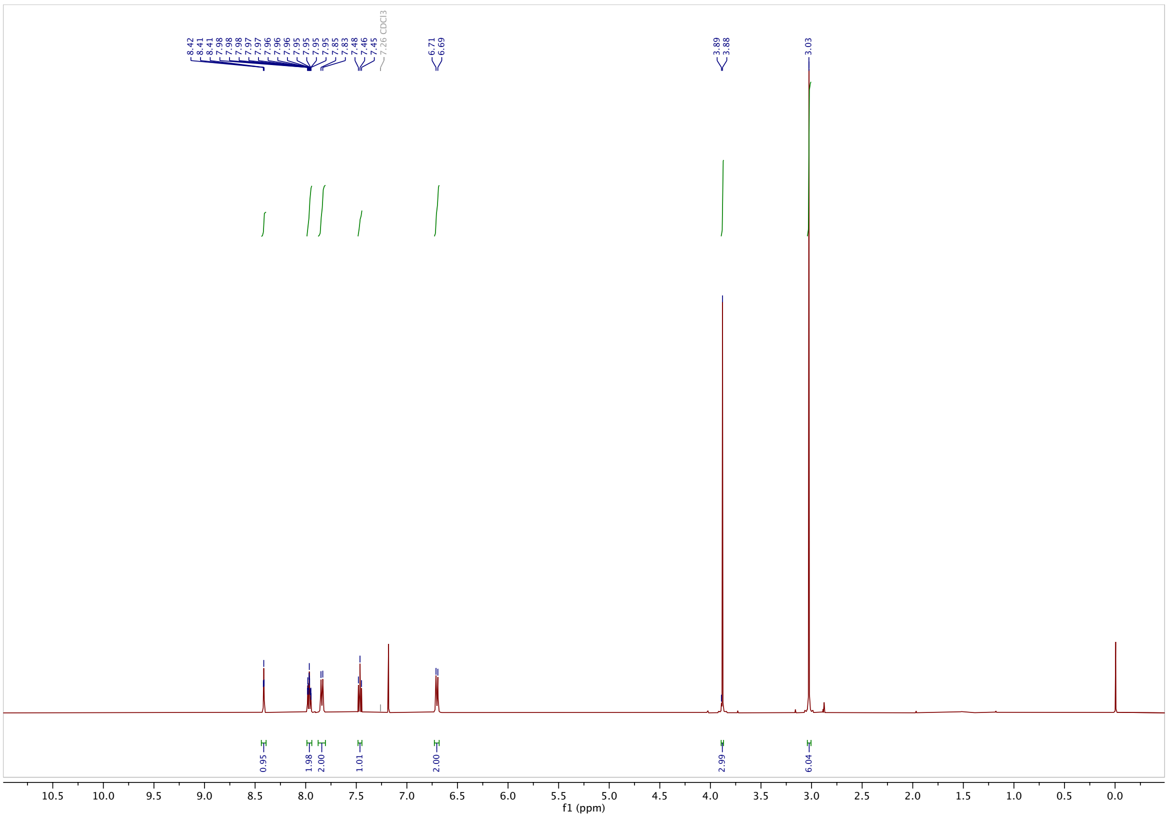

**¹³C-NMR**

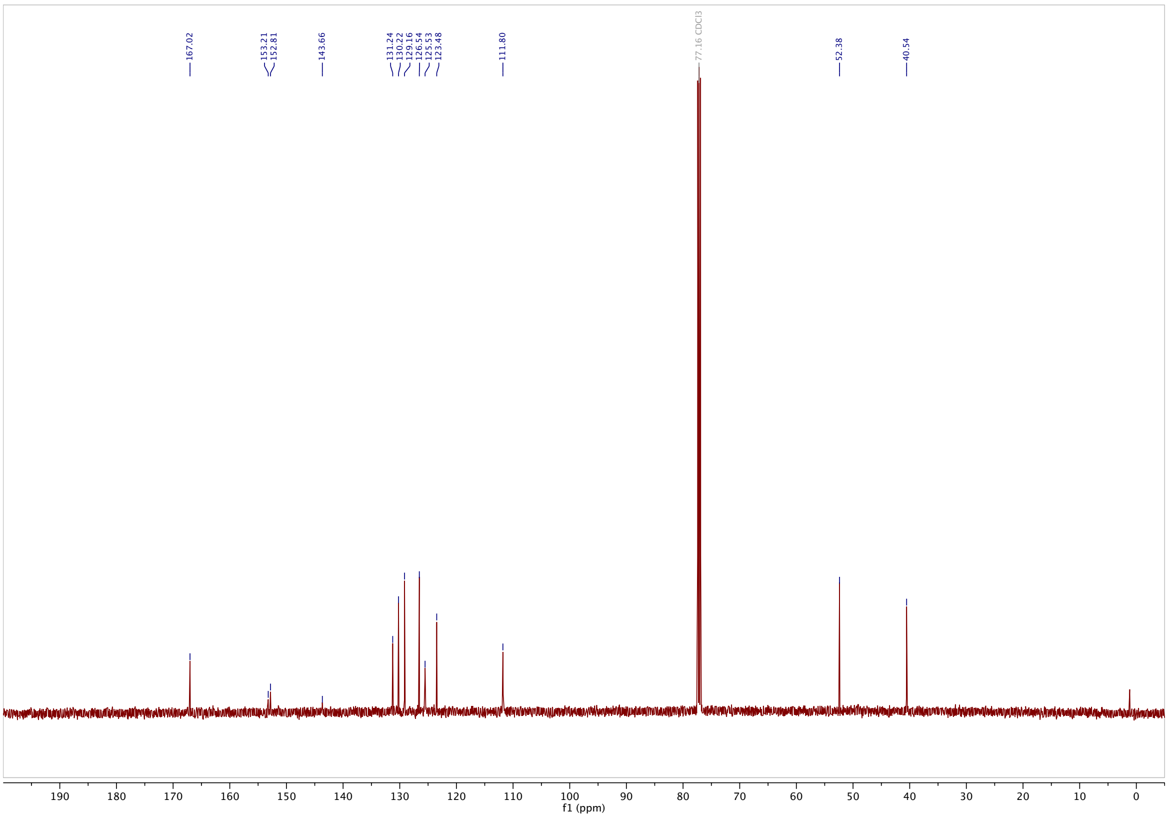

#### 3-((4-(dimethylamino)phenyl)diazenyl)benzoic acid (3DMA-$CO_2$H): [1]H-NMR

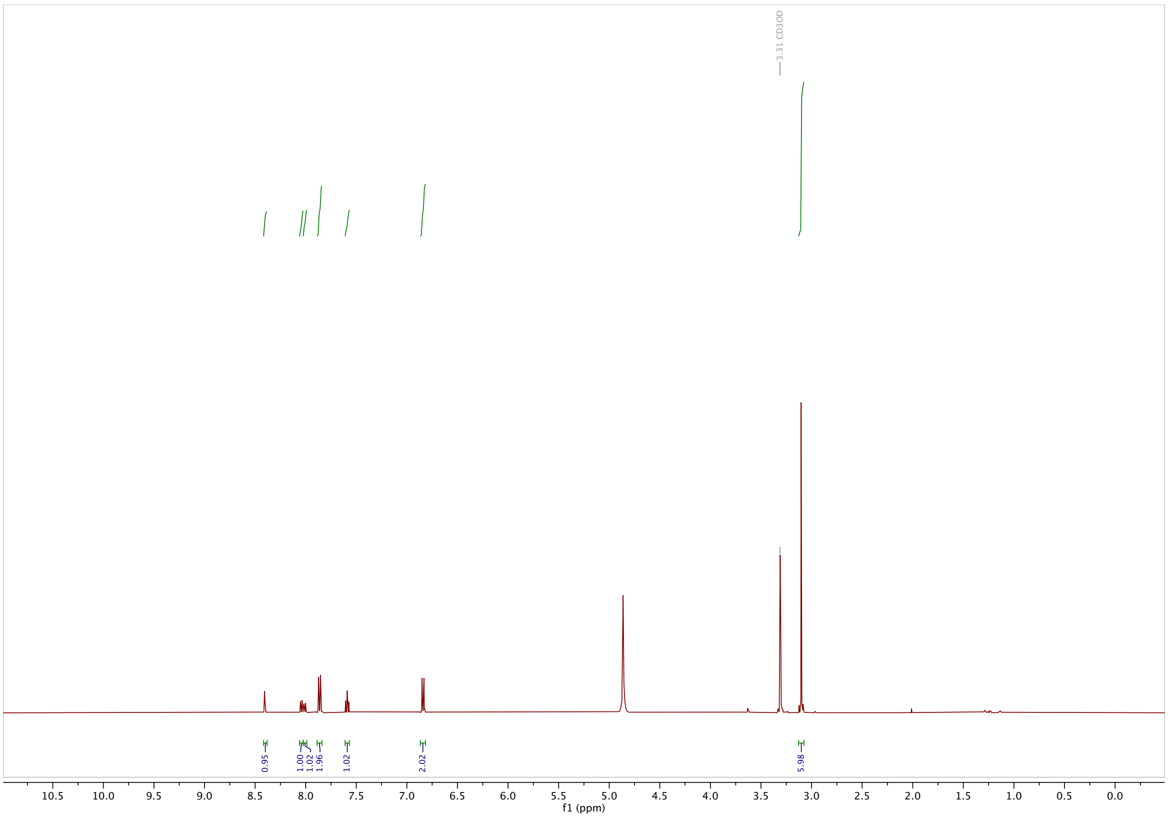

#### [13]C-NMR

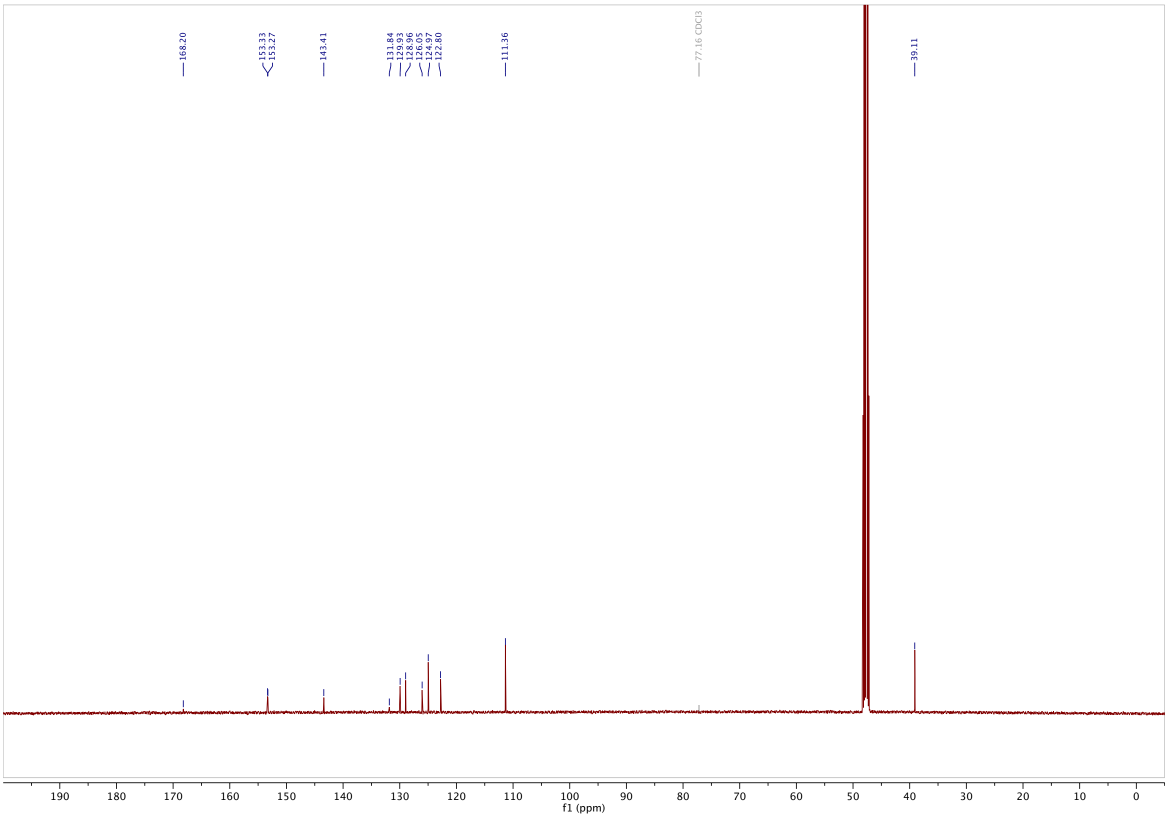

**methyl 3-((4-hydroxy-3,5-dimethoxyphenyl)diazenyl)-4-methoxybenzoate (S7): $^{1}$H-NMR**

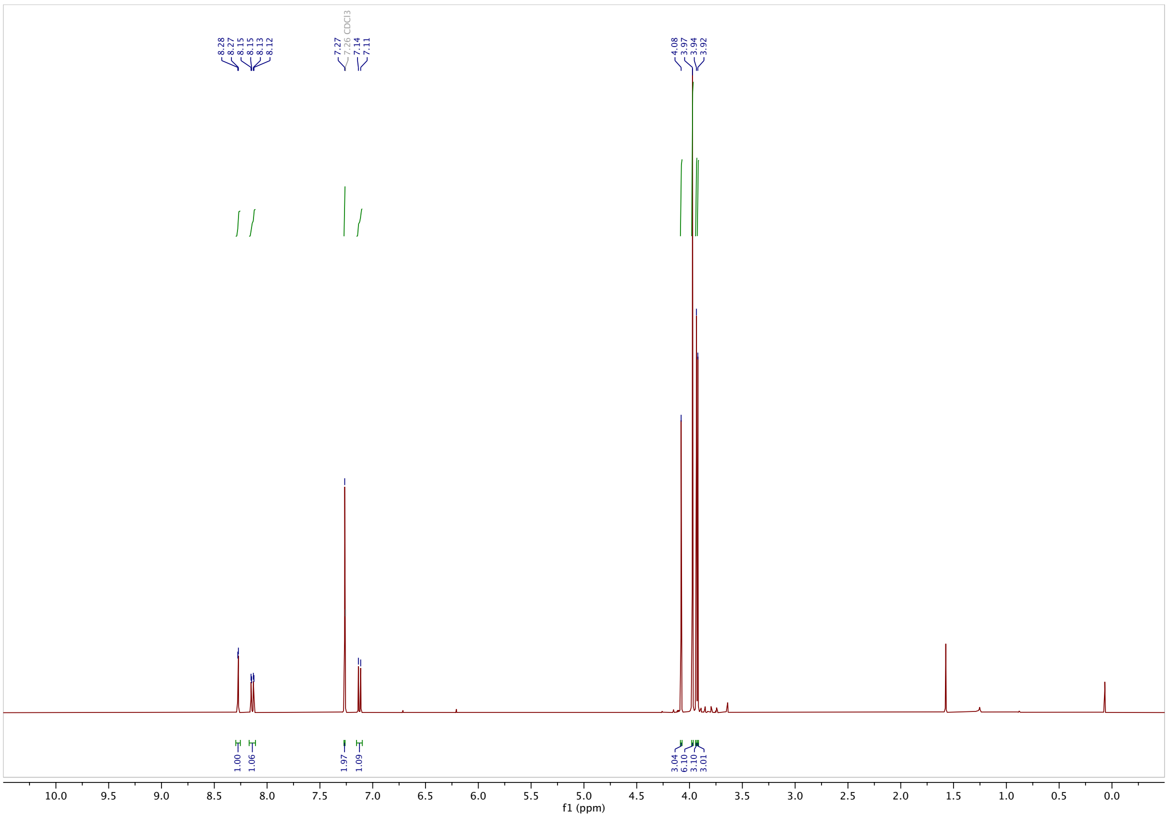

**$^{13}$C-NMR**

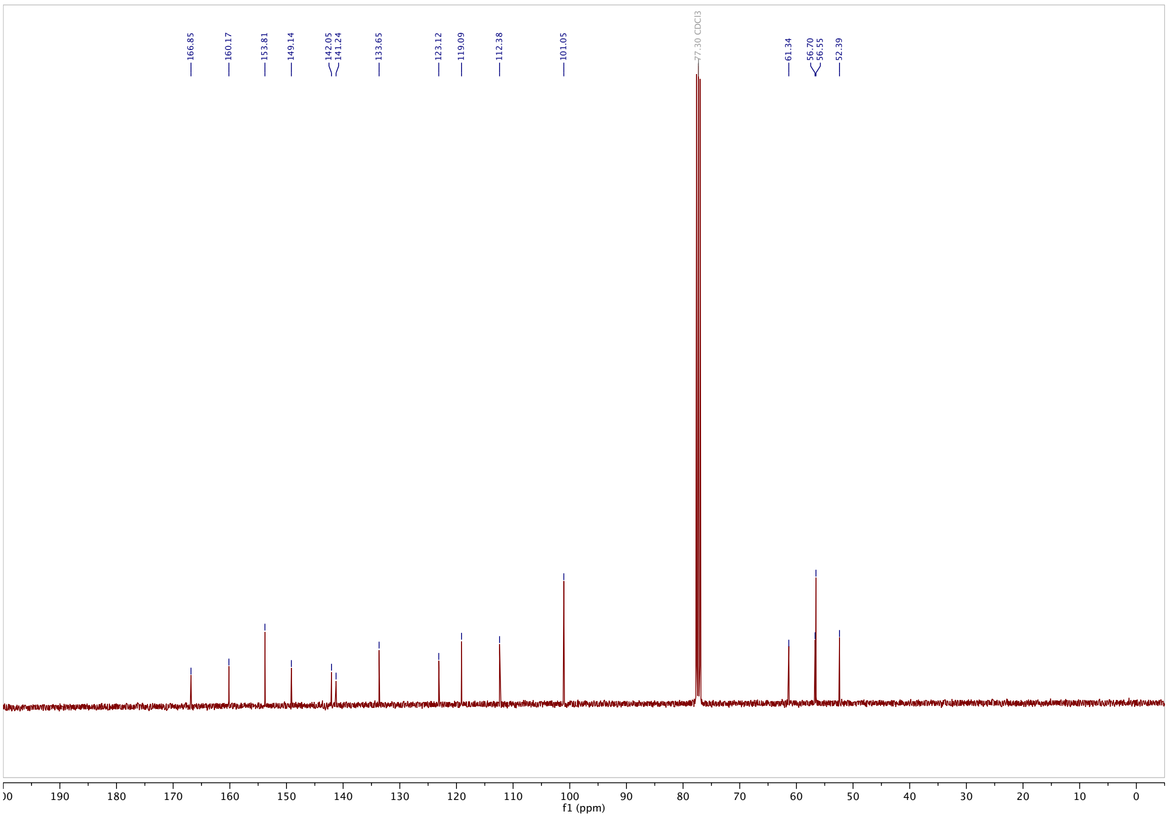

**methyl 4-methoxy-3-((3,4,5-trimethoxyphenyl)diazenyl)benzoate** (**S8**): $^{1}$H-NMR

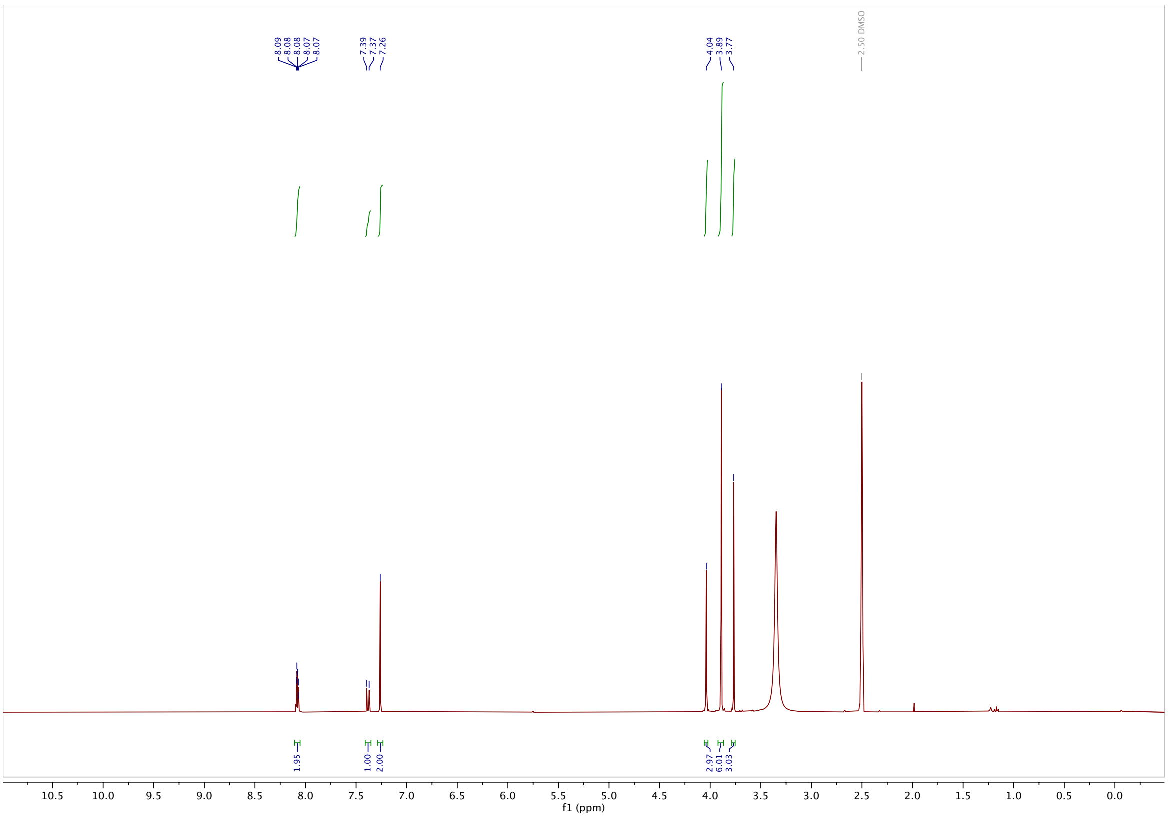

$^{13}$C-NMR

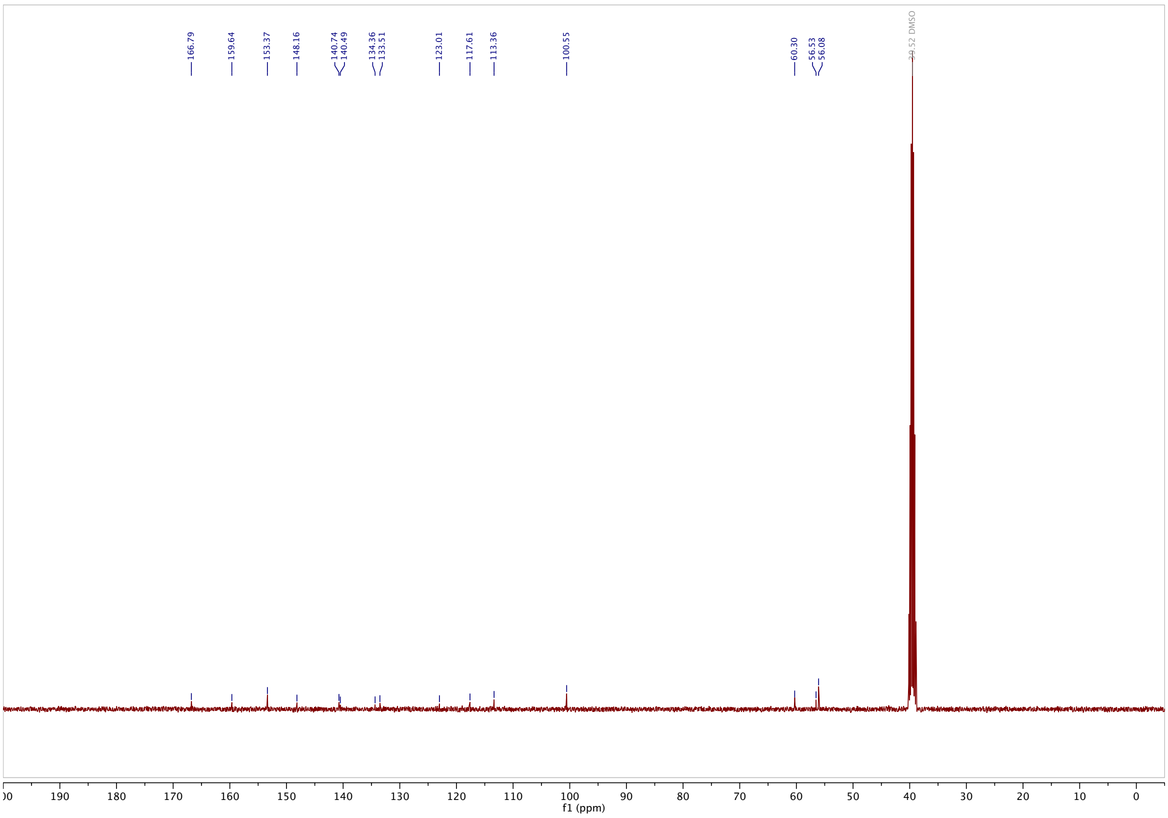

#### 4-methoxy-3-((3,4,5-trimethoxyphenyl)diazenyl)benzoic acid (3MTM-$CO_2$H): $^{1}$H-NMR

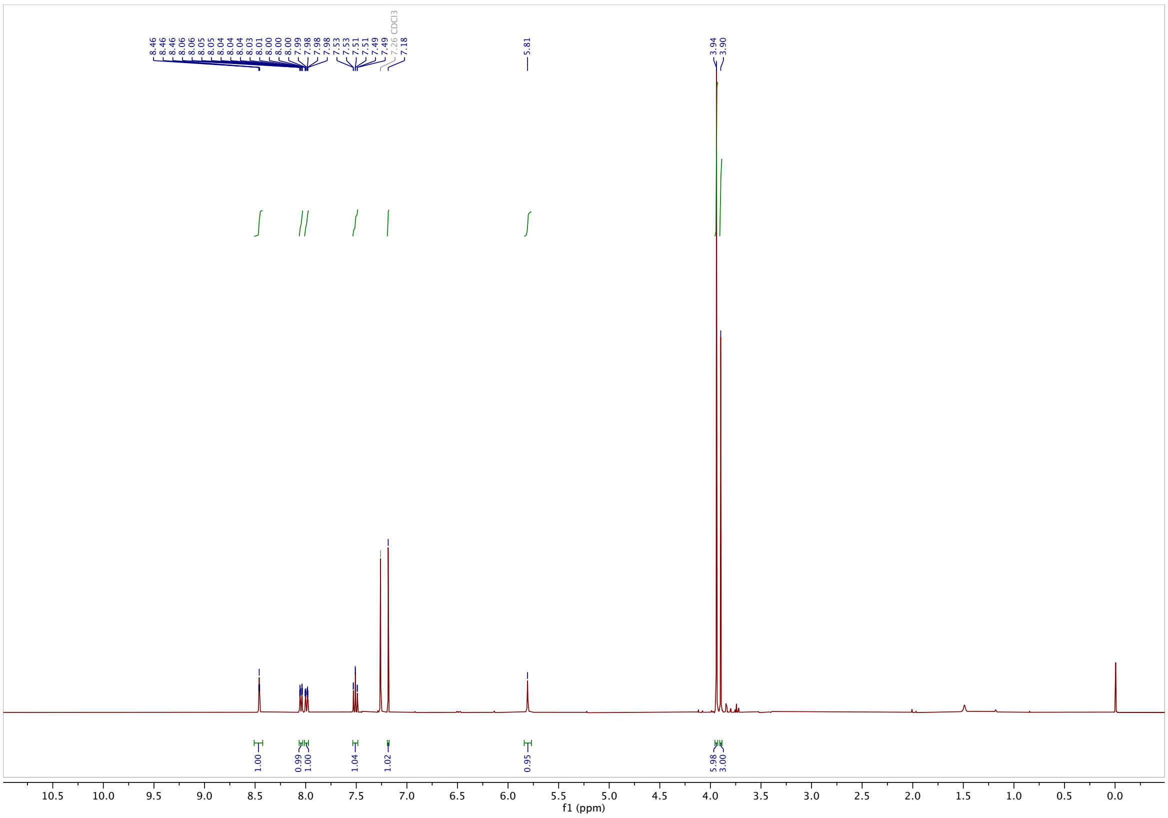

#### $^{13}$C-NMR

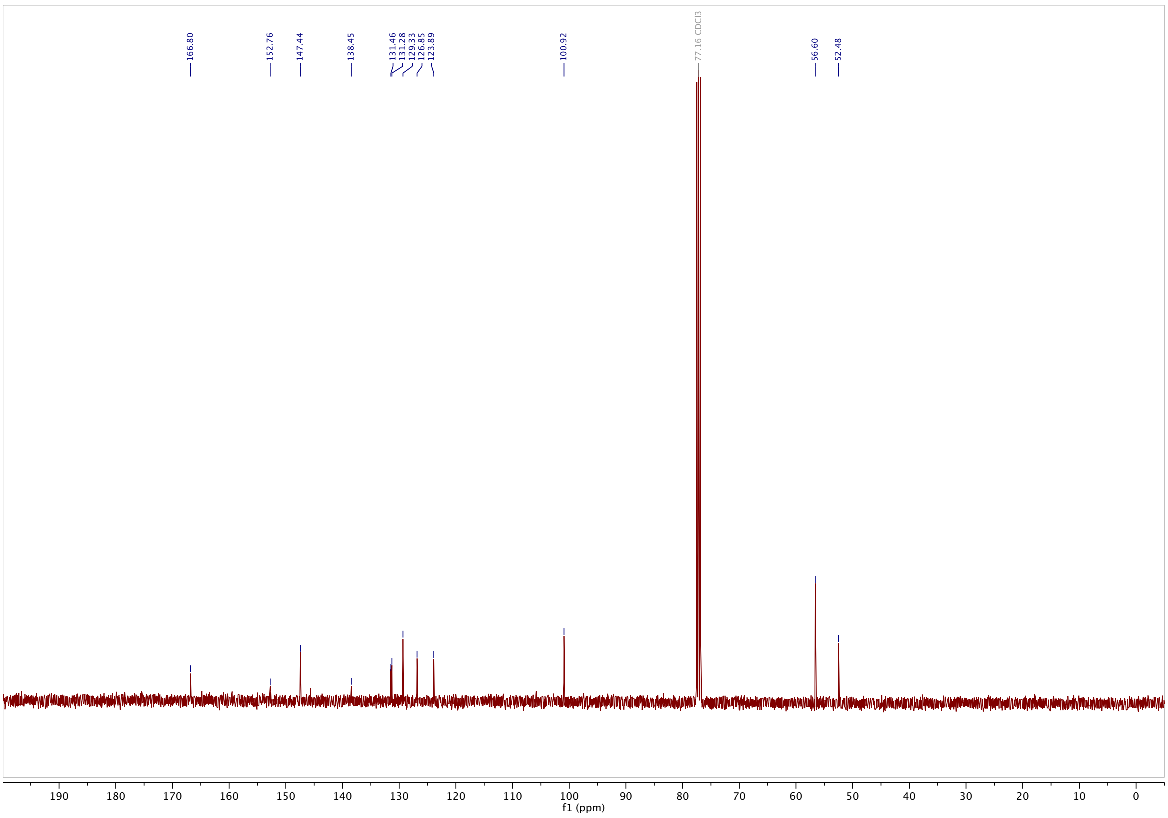

**Methyl 3-((4-hydroxy-3,5-dimethoxyphenyl)diazenyl)benzoate (S9): $^{1}$H-NMR**

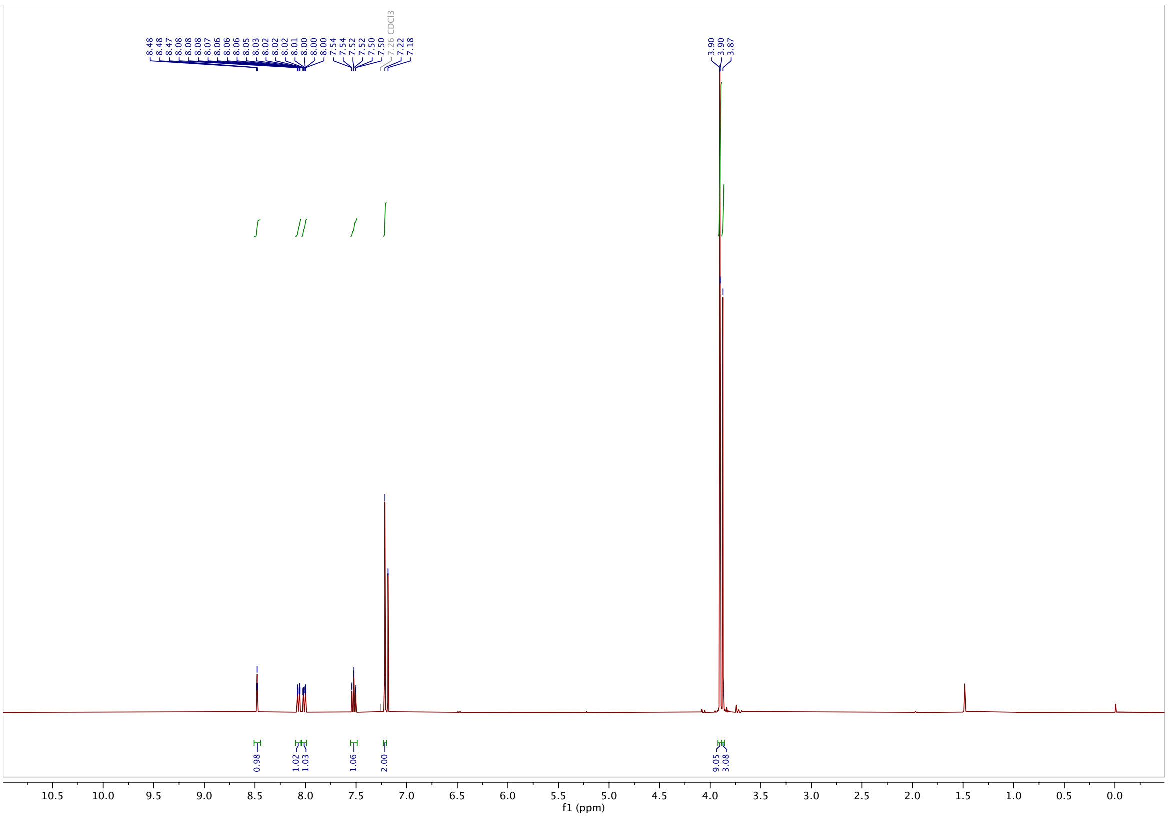

**$^{13}$C-NMR**

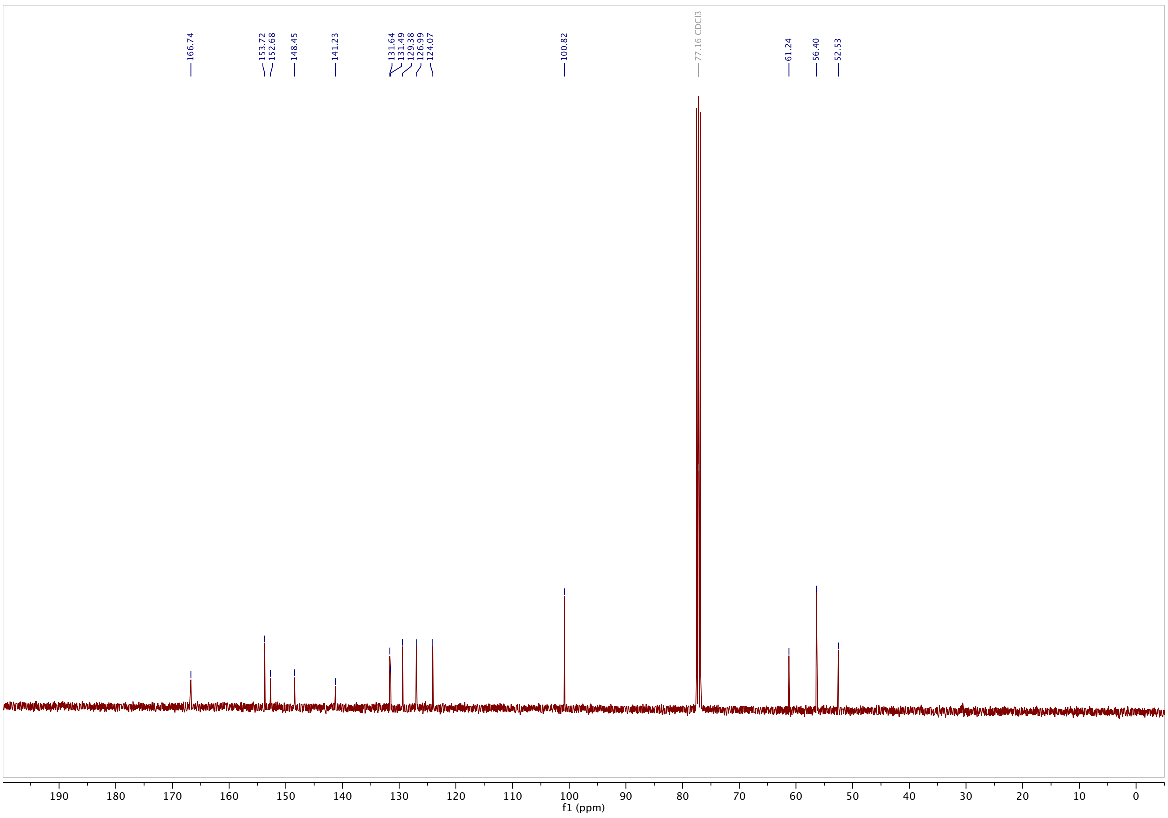

**methyl 3-((3,4,5-trimethoxyphenyl)diazenyl)benzoate (S10): $^{1}$H-NMR**

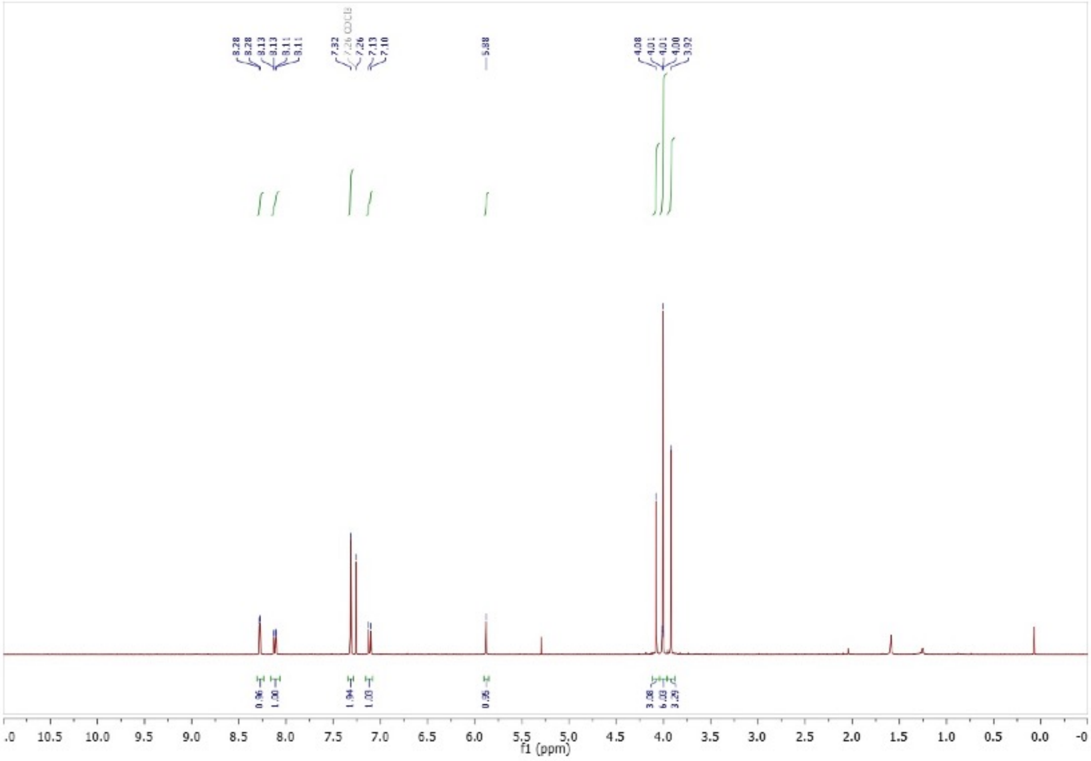

**$^{13}$C-NMR**

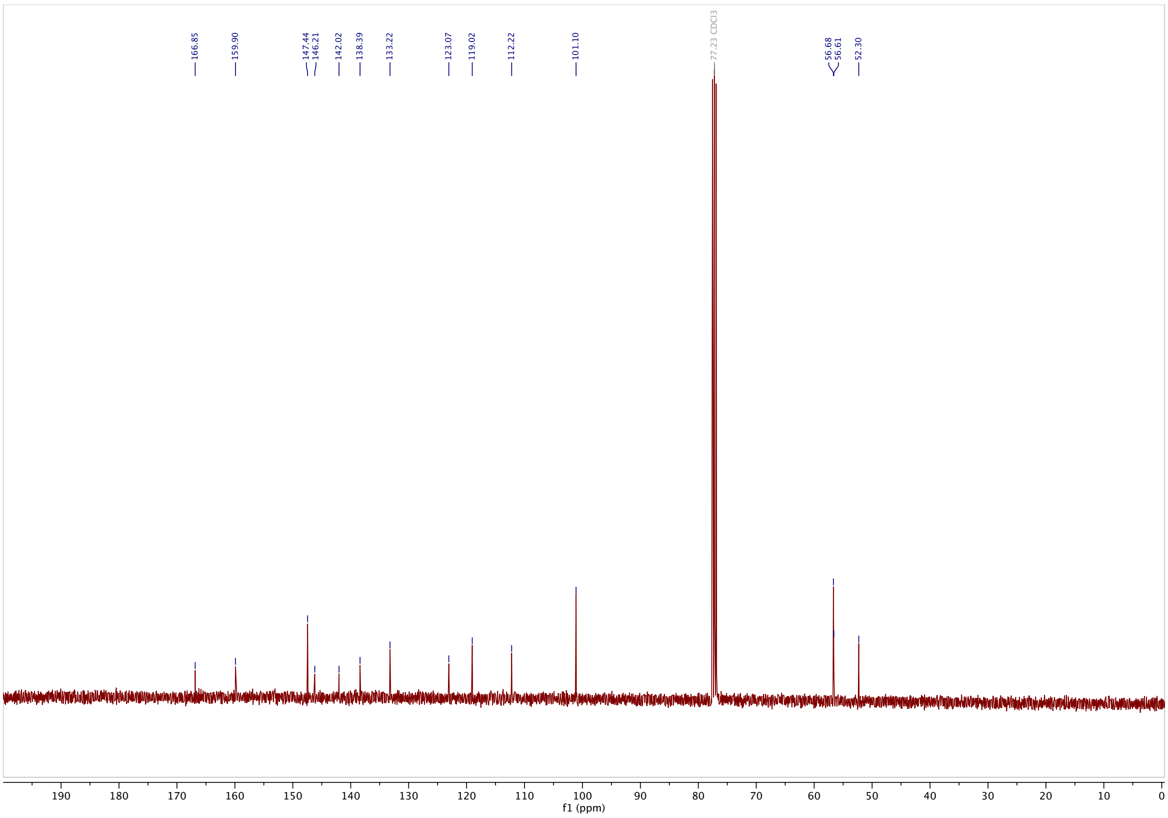

#### 3-((3,4,5-trimethoxyphenyl)diazenyl)benzoic acid (3TM-$CO_2$H): [1]H-NMR

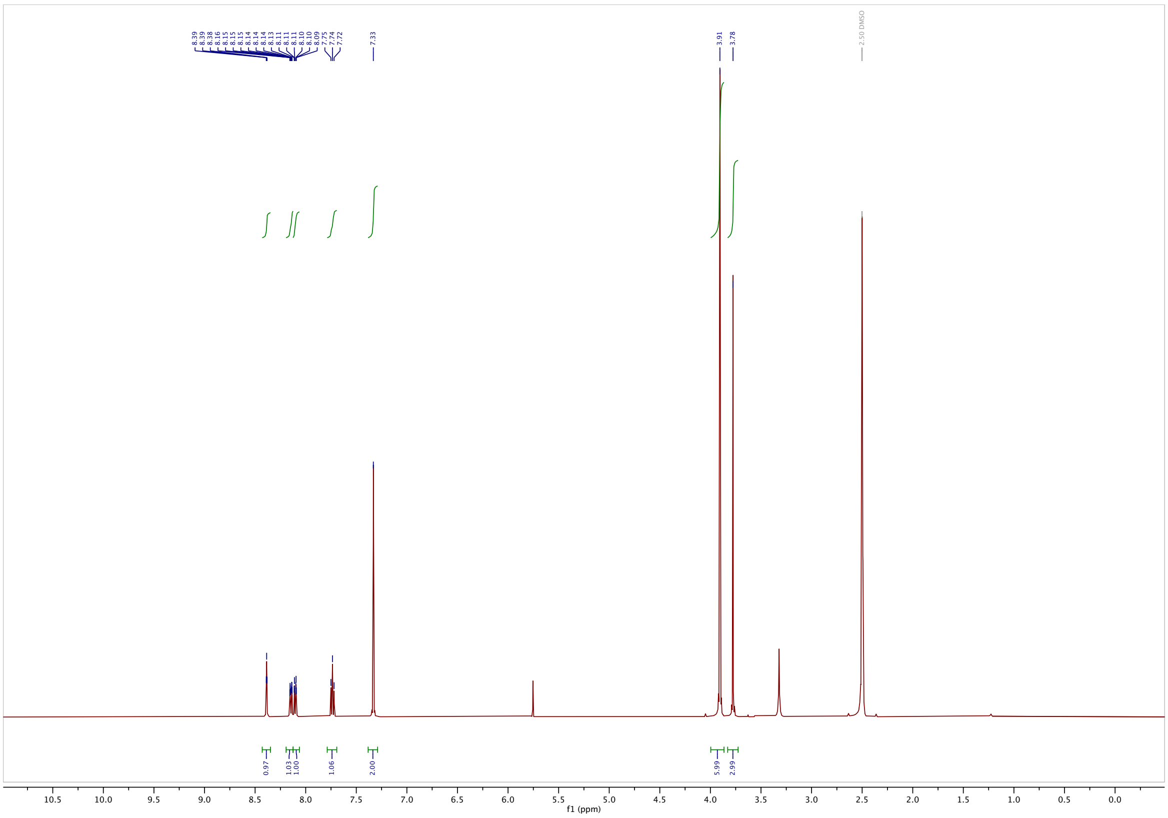

#### [13]C-NMR

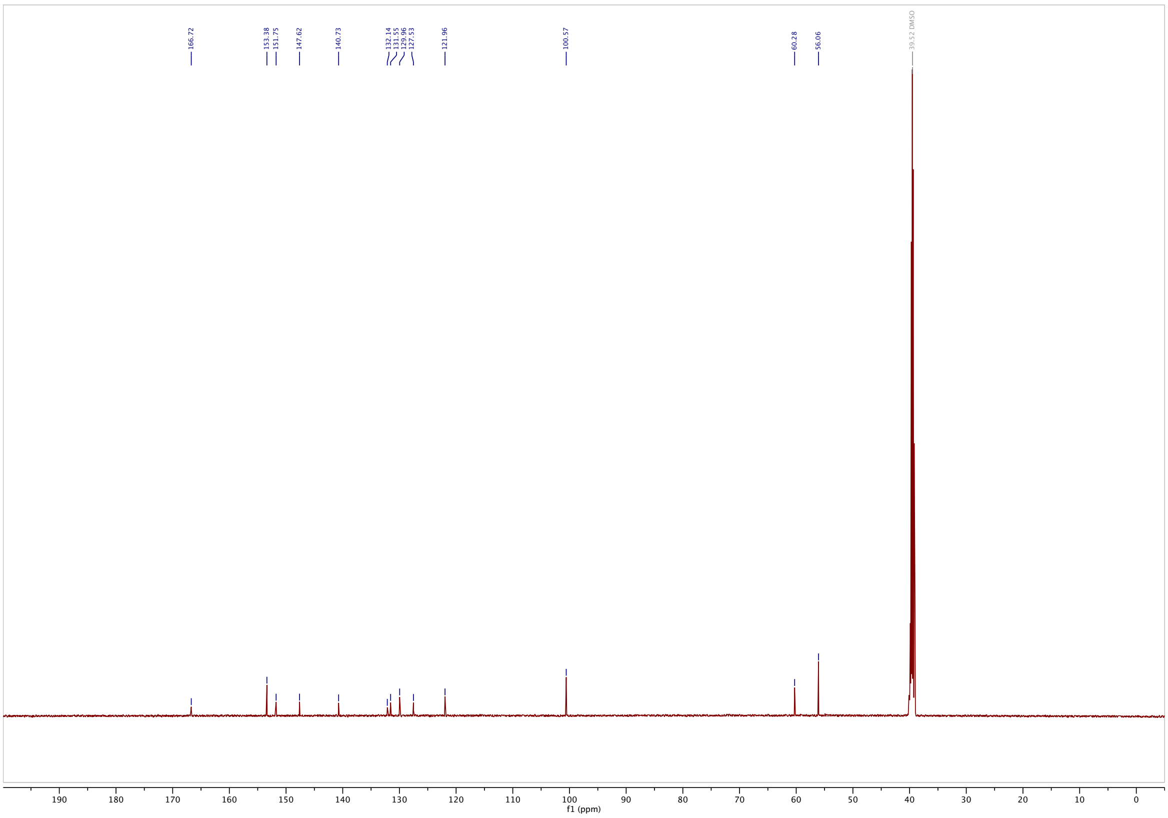

**methyl 3-((4-(bis(2-hydroxyethyl)amino)phenyl)diazenyl)benzoate (S11)**: $^{1}$**H-NMR**

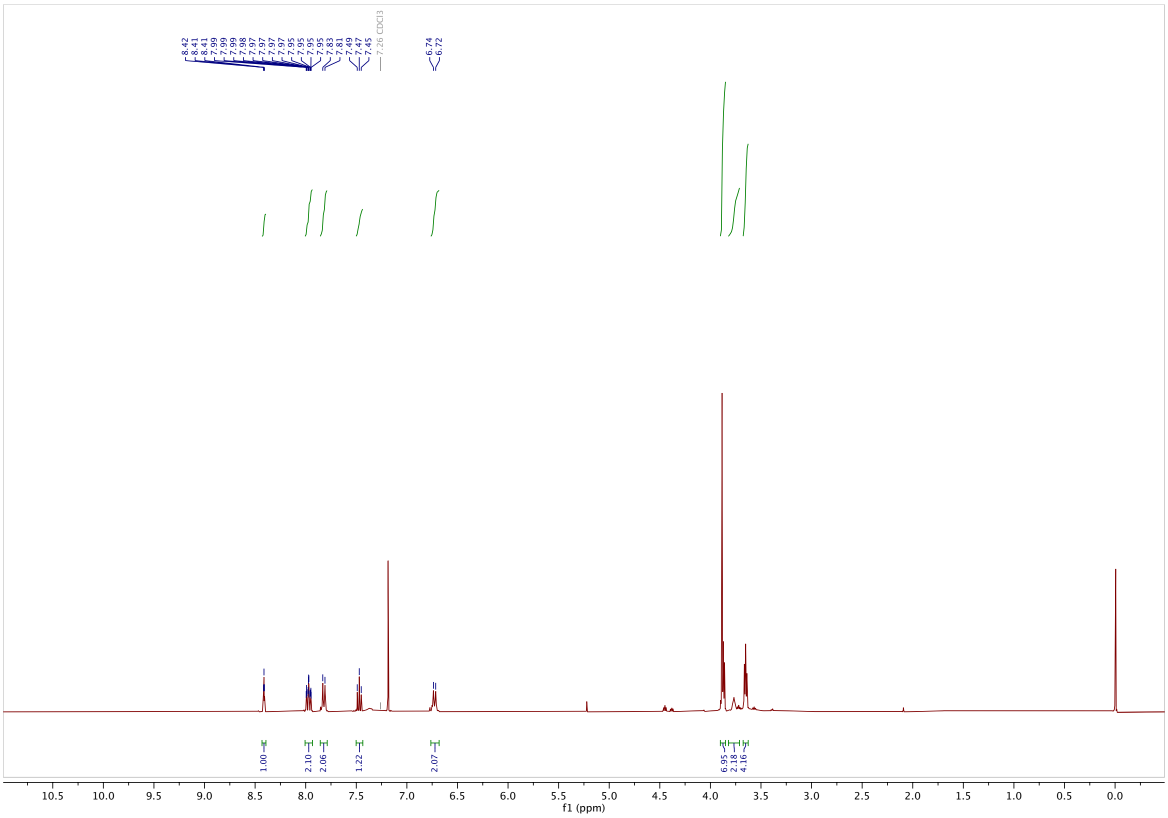

$^{13}$**C-NMR**

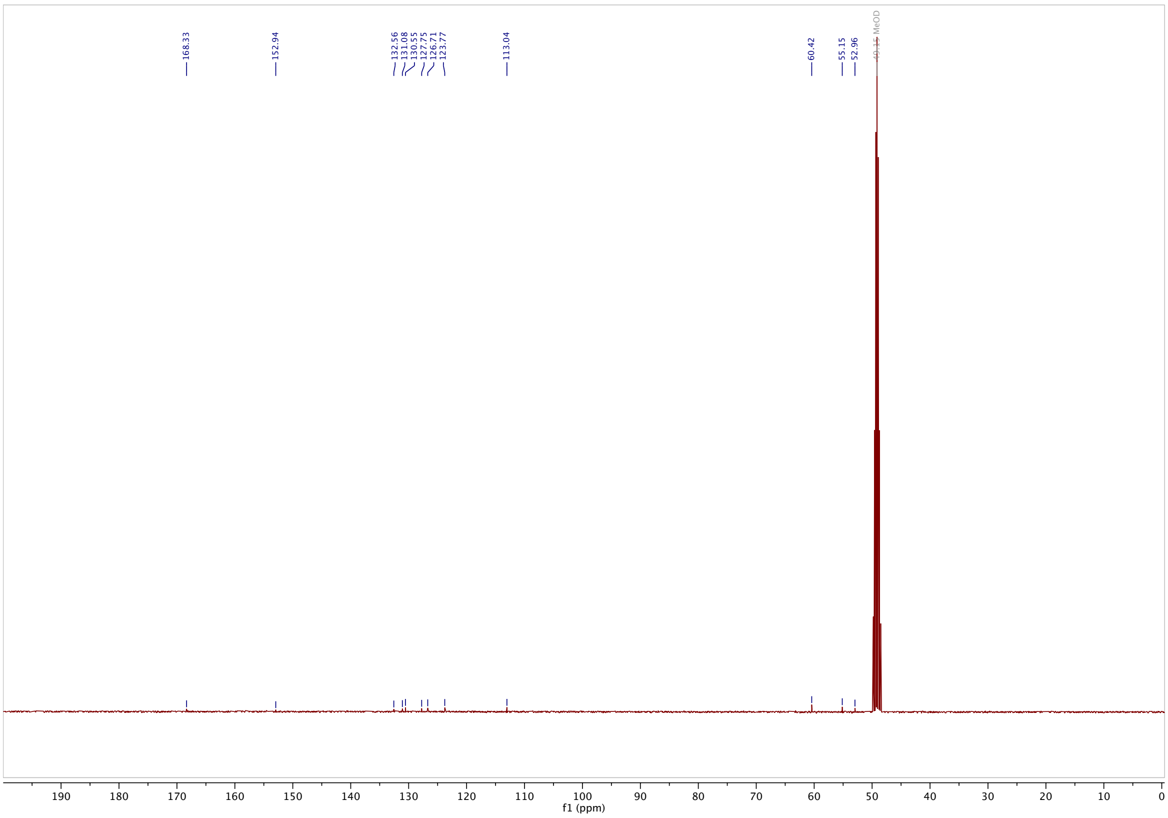

**3-((4-(bis(2-hydroxyethyl)amino)phenyl)diazenyl)benzoic acid (3DEA-$CO_2$H): $^1$H-NMR**

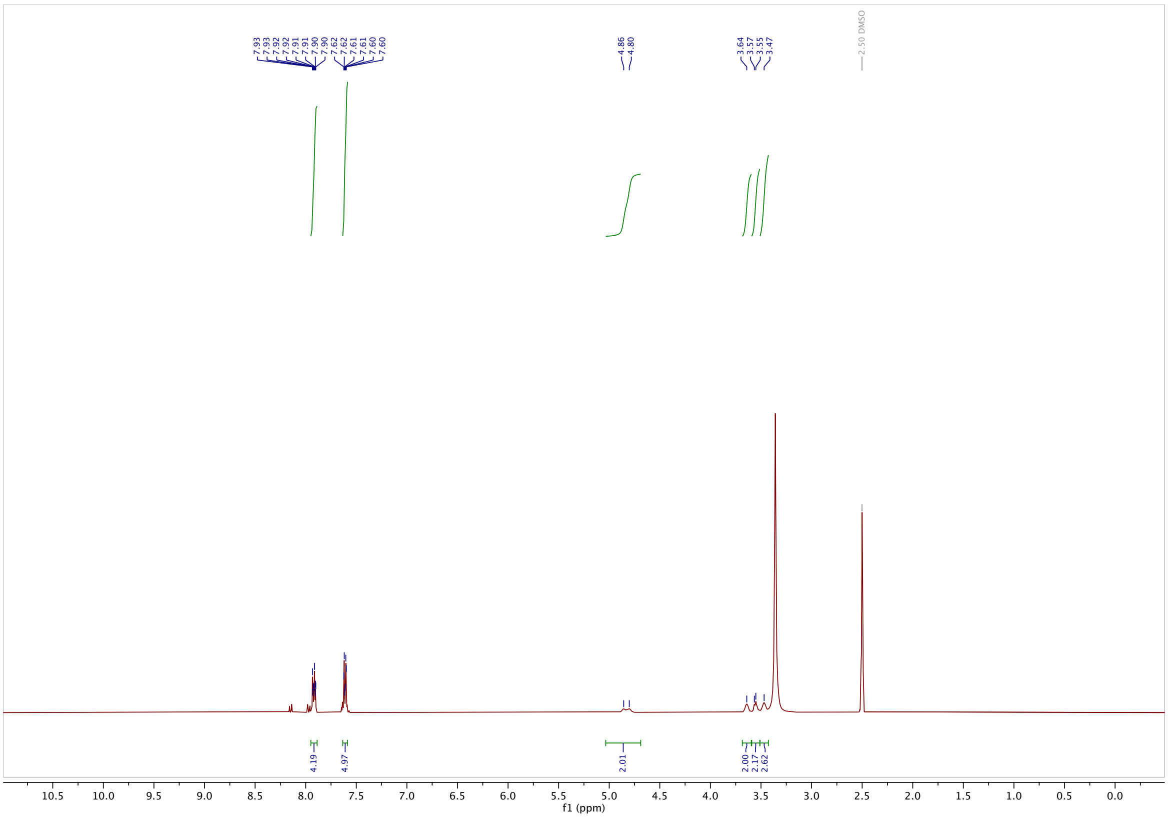

**$^{13}$C-NMR**

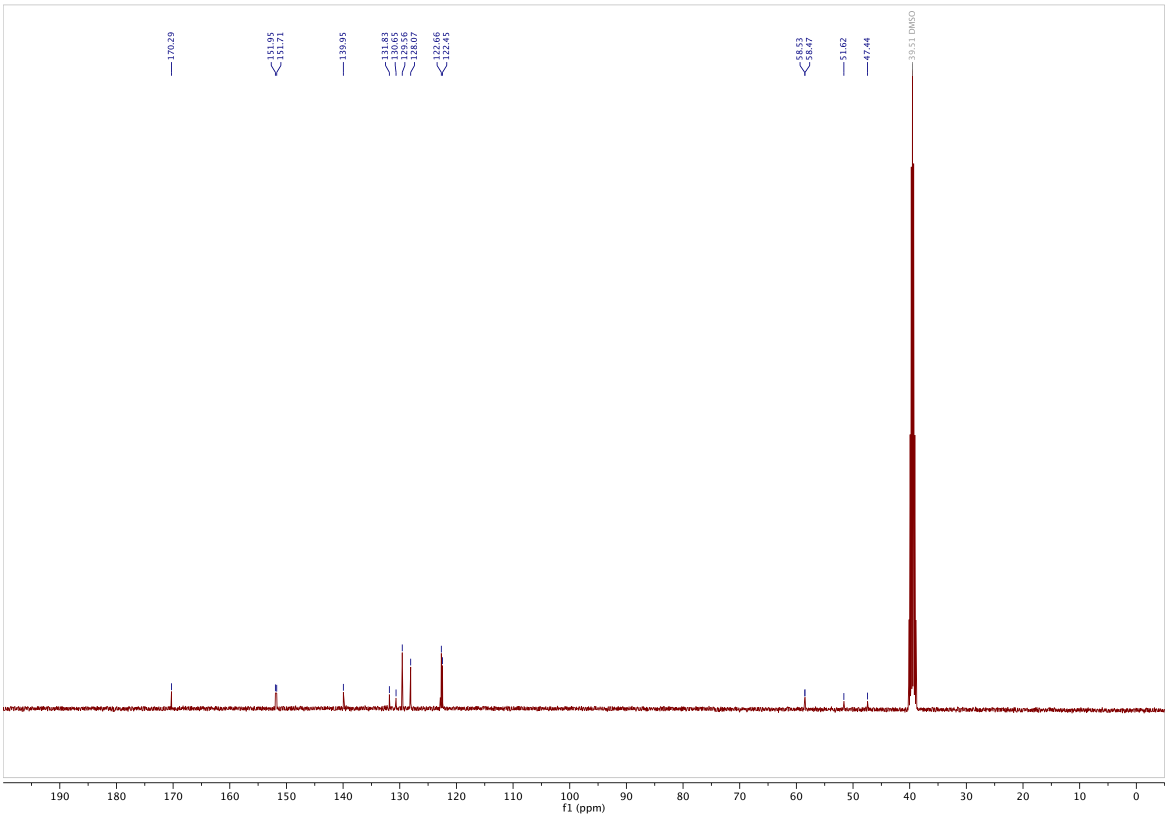

#### methyl 2-((4-hydroxyphenyl)diazenyl)benzoate (S12): $^{1}$H-NMR

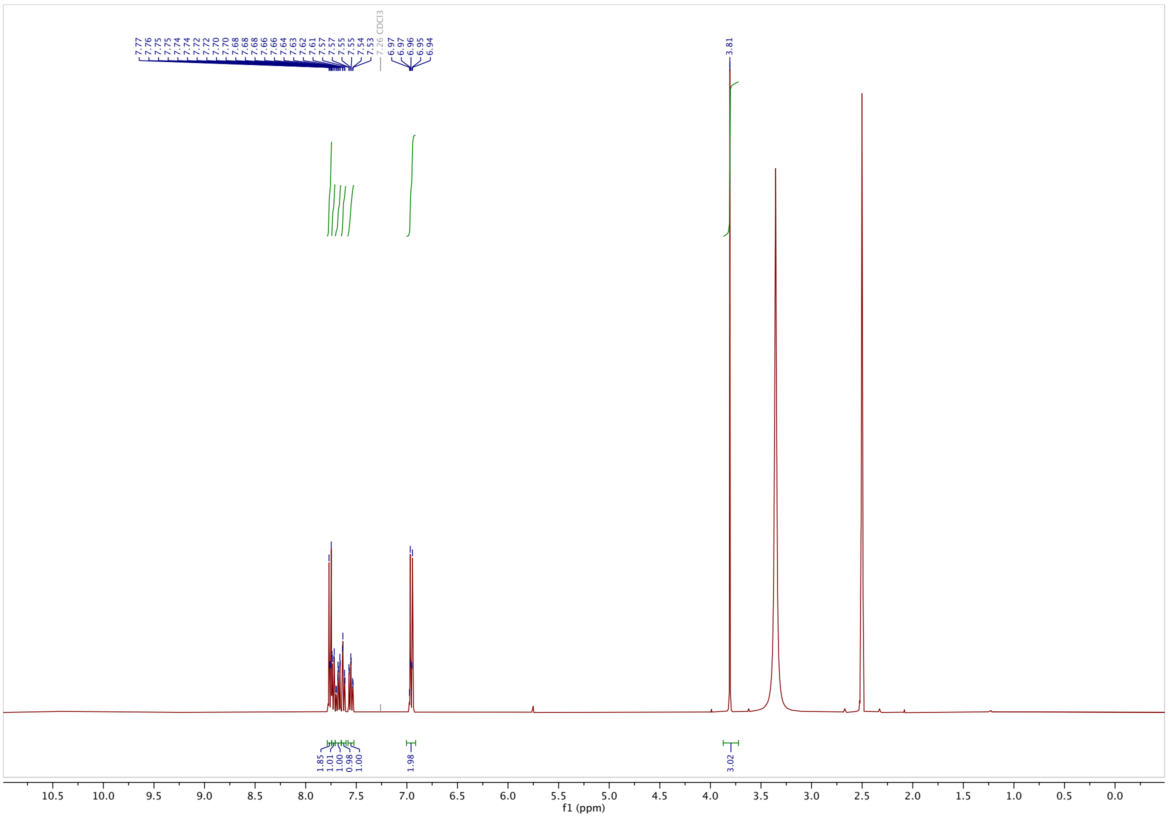

#### $^{13}$C-NMR

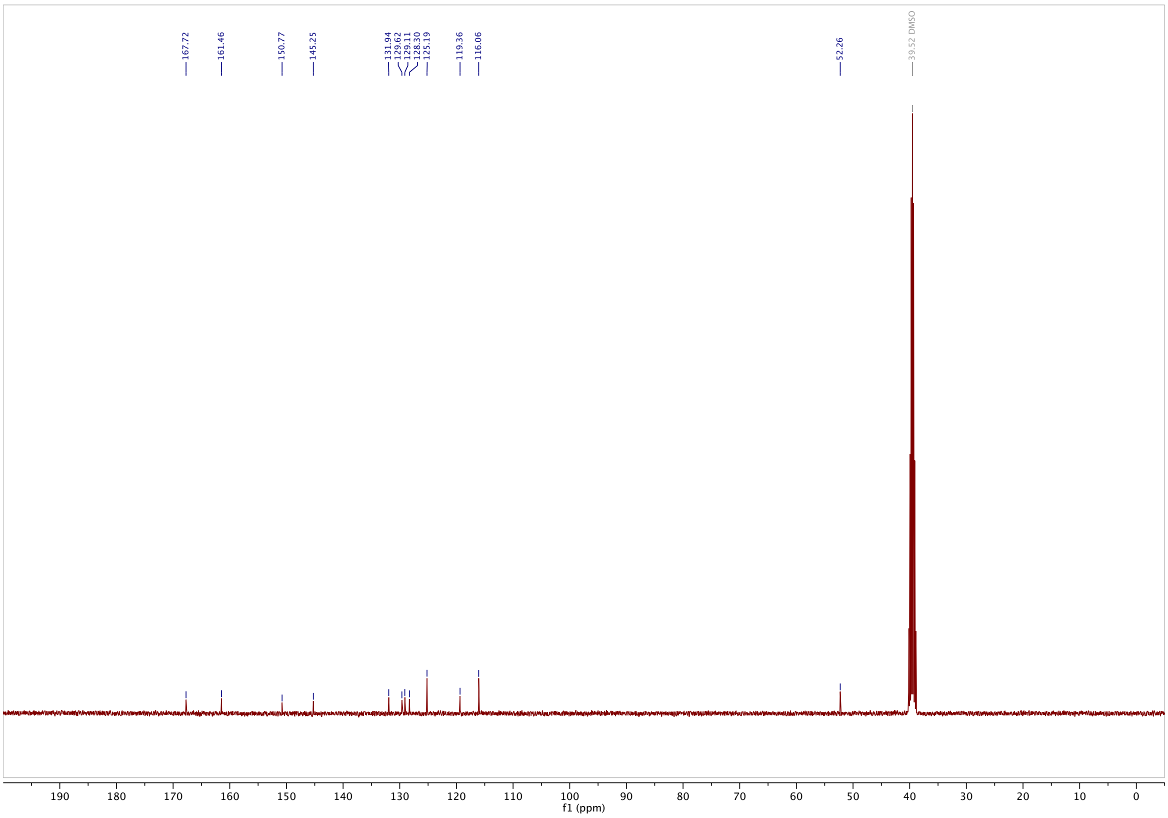

**methyl 2-((4-methoxyphenyl)diazenyl)benzoate (S13)**: **$^1$H-NMR**

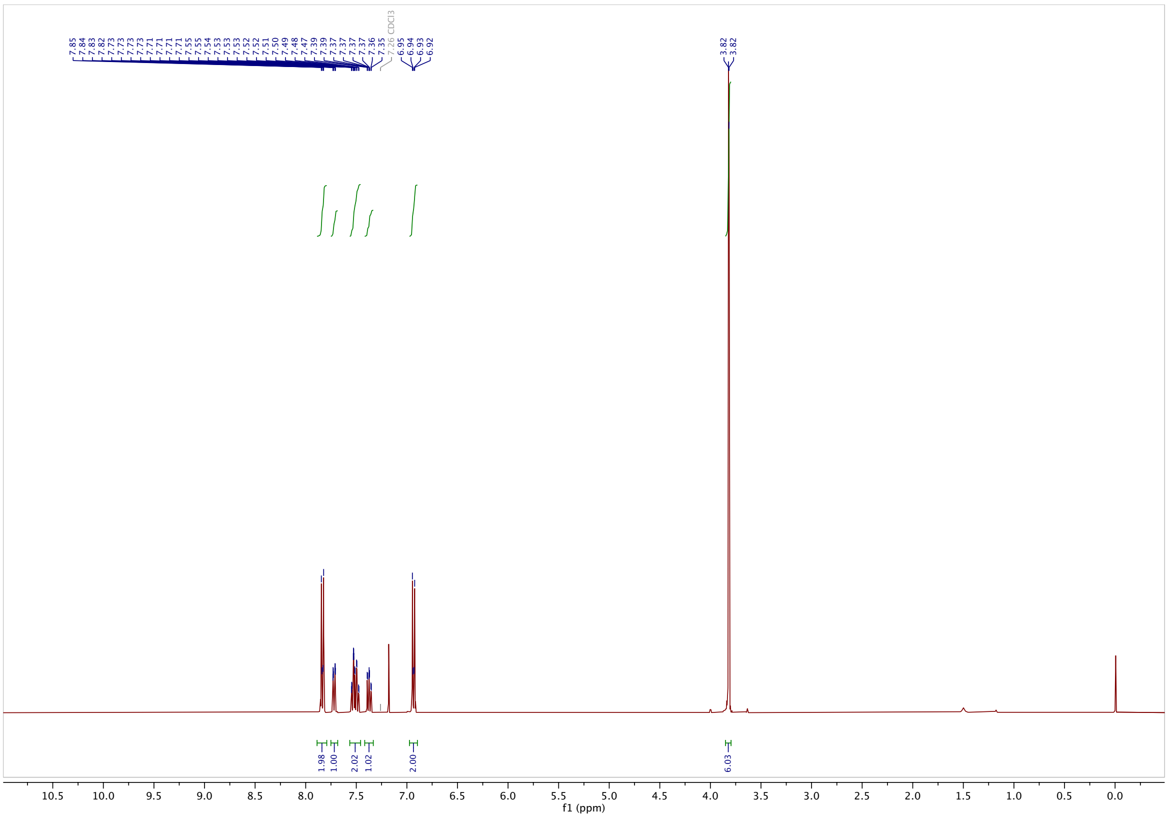

**$^{13}$C-NMR**

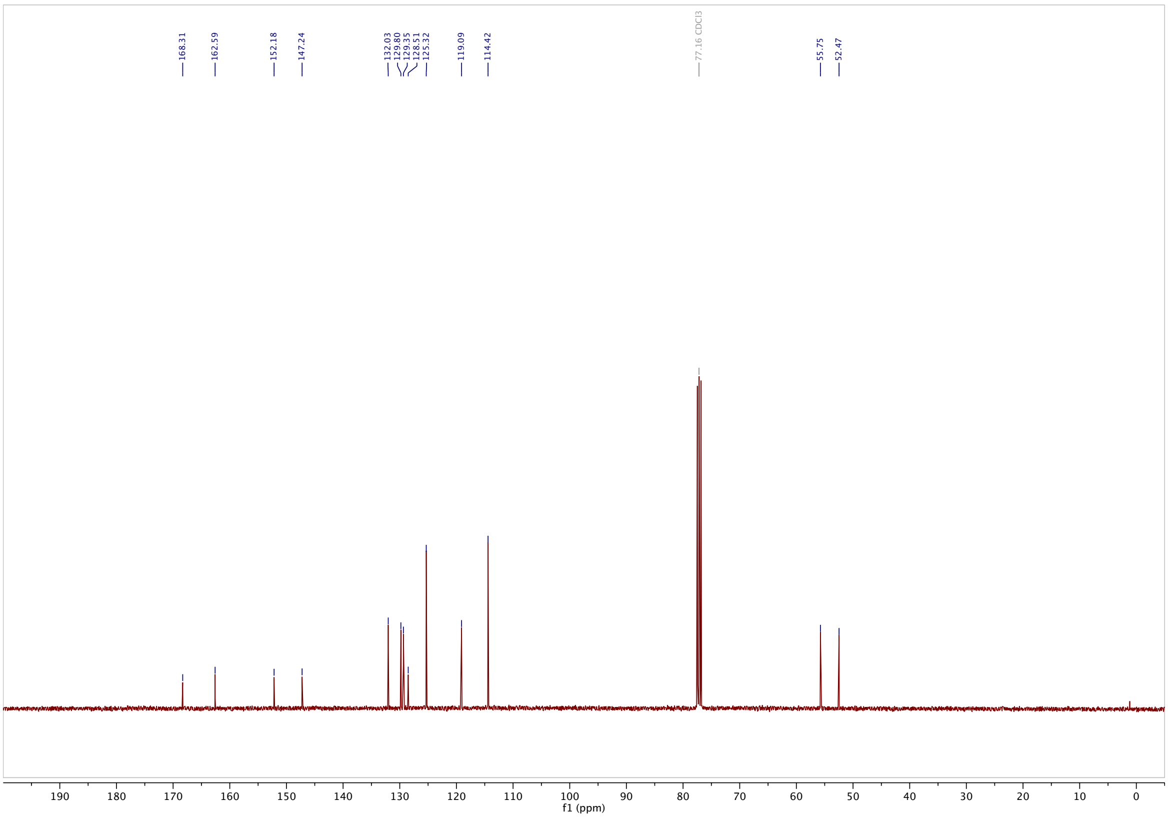

#### 2-((4-methoxyphenyl)diazenyl)benzoic acid (2MP-$CO_2$H): $^1$H-NMR

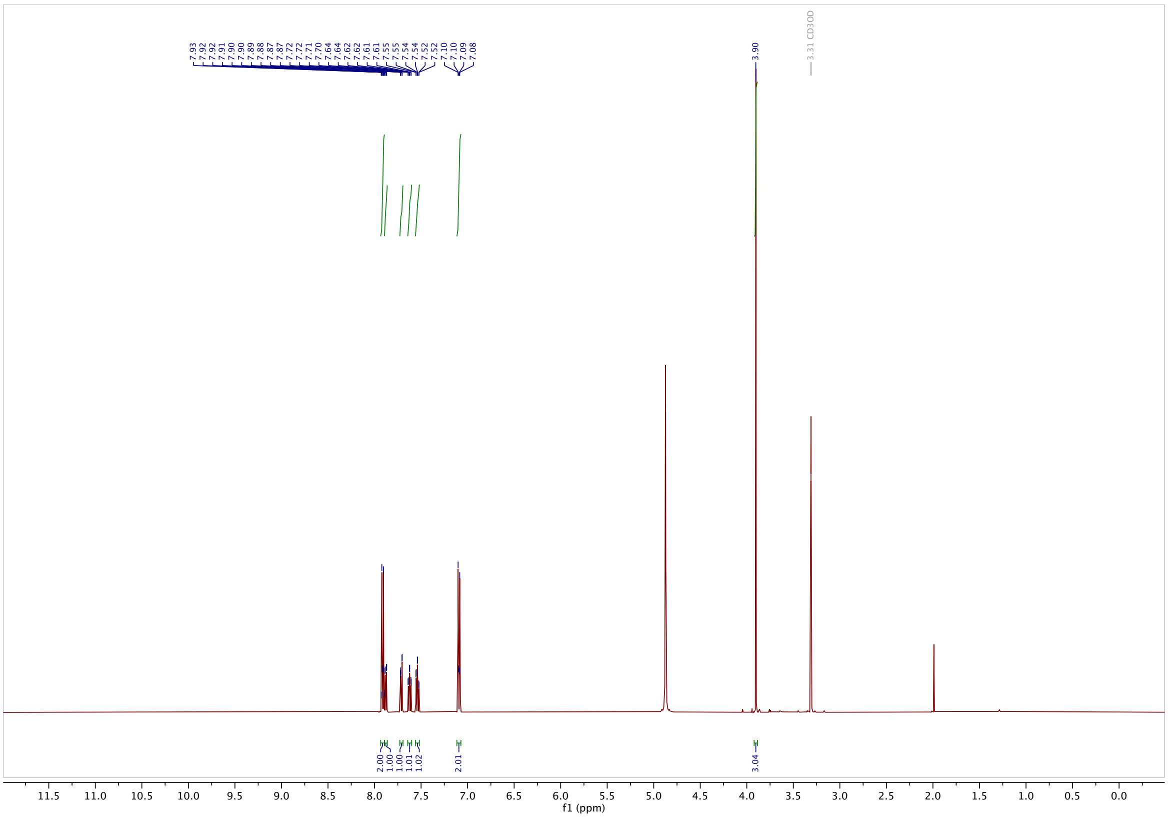

#### $^{13}$C-NMR

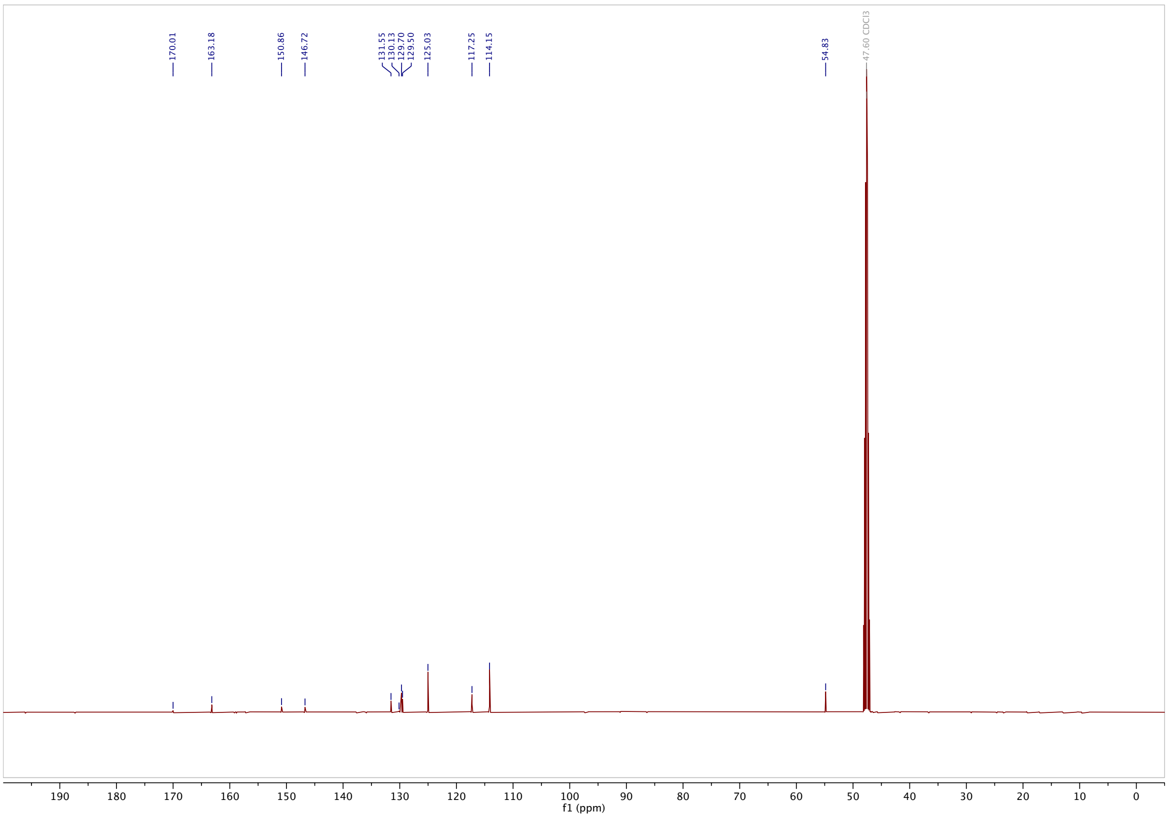

#### AzTax4H: $^{1}$H-NMR

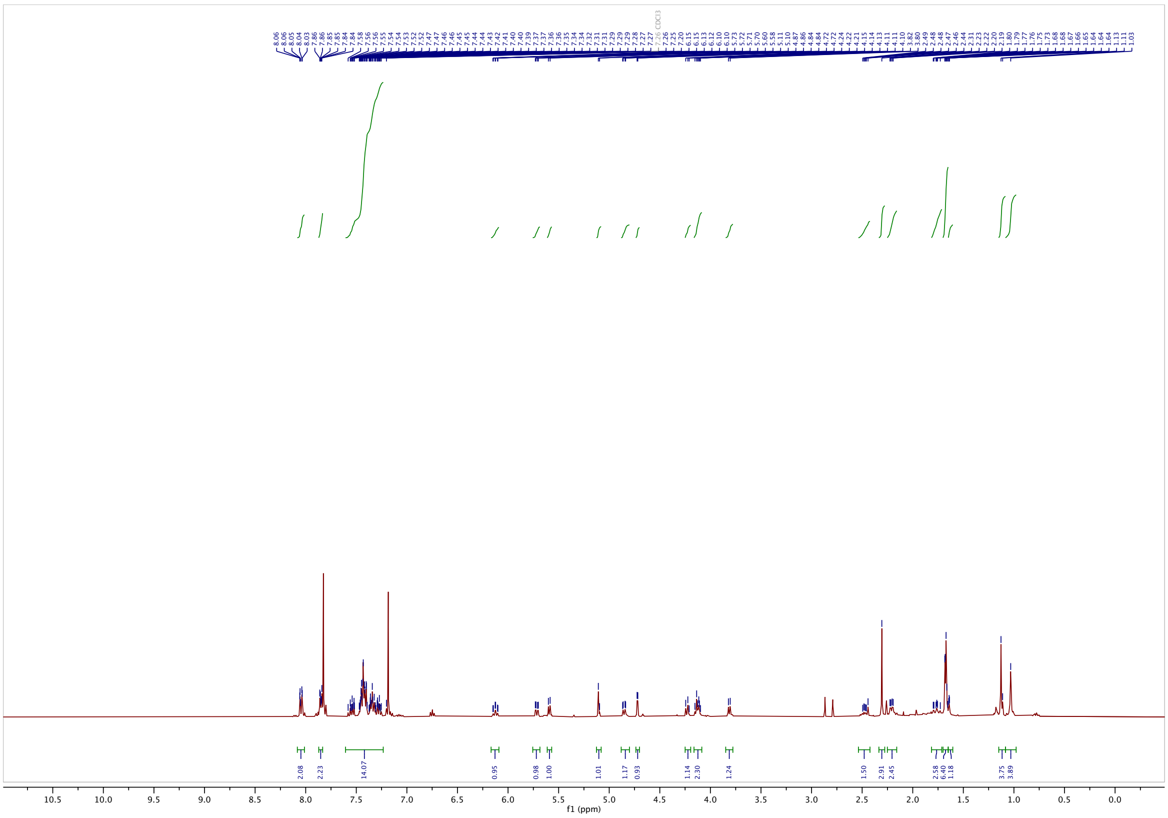

#### $^{13}$C-NMR

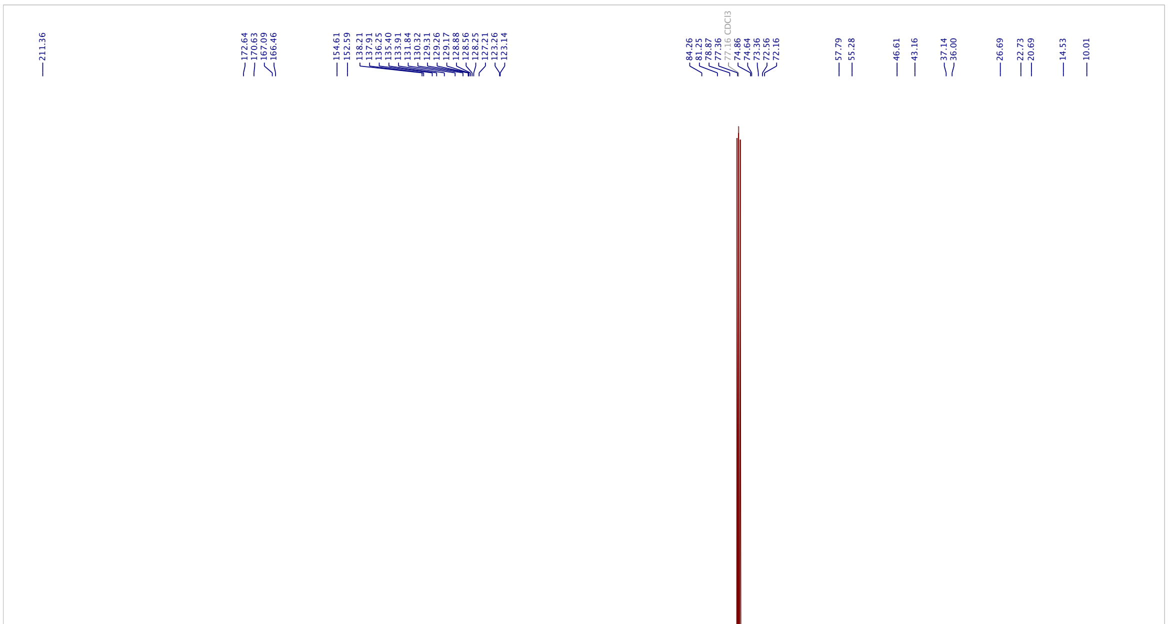

#### AzTax4DMA: $^1$H-NMR

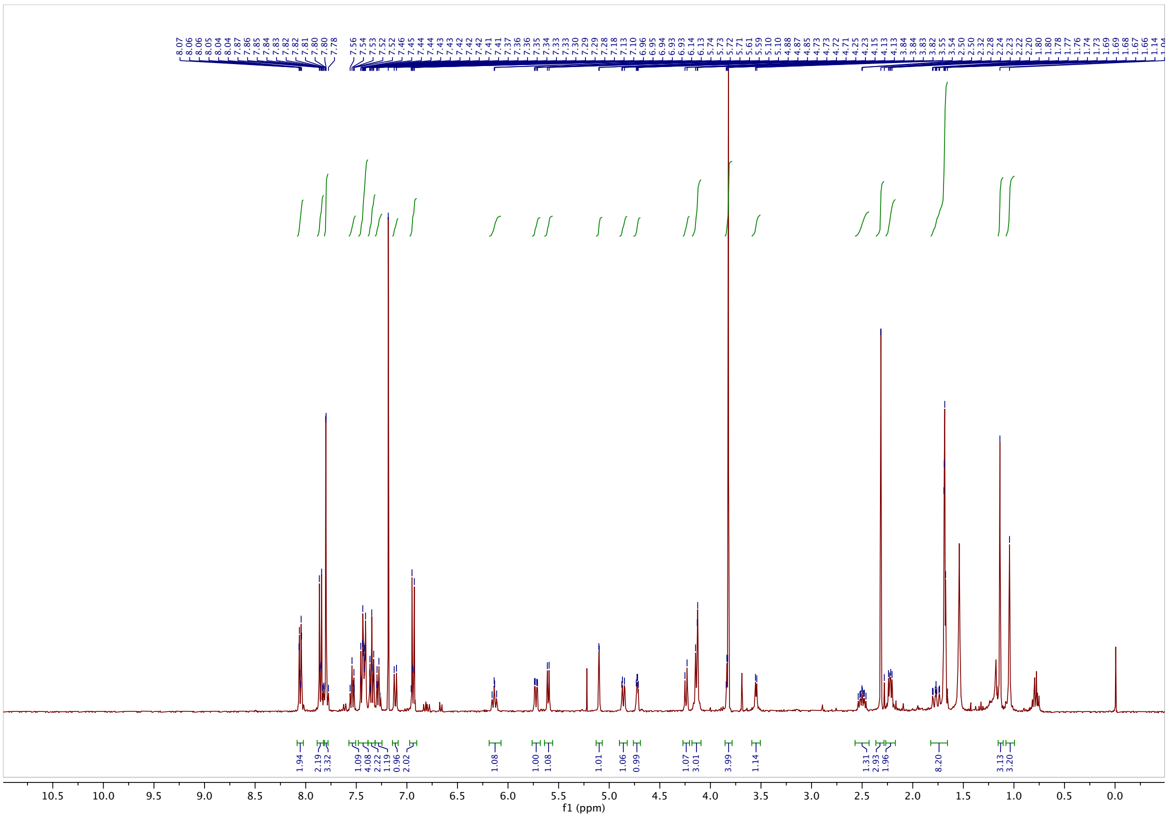

#### $^{13}$C-NMR

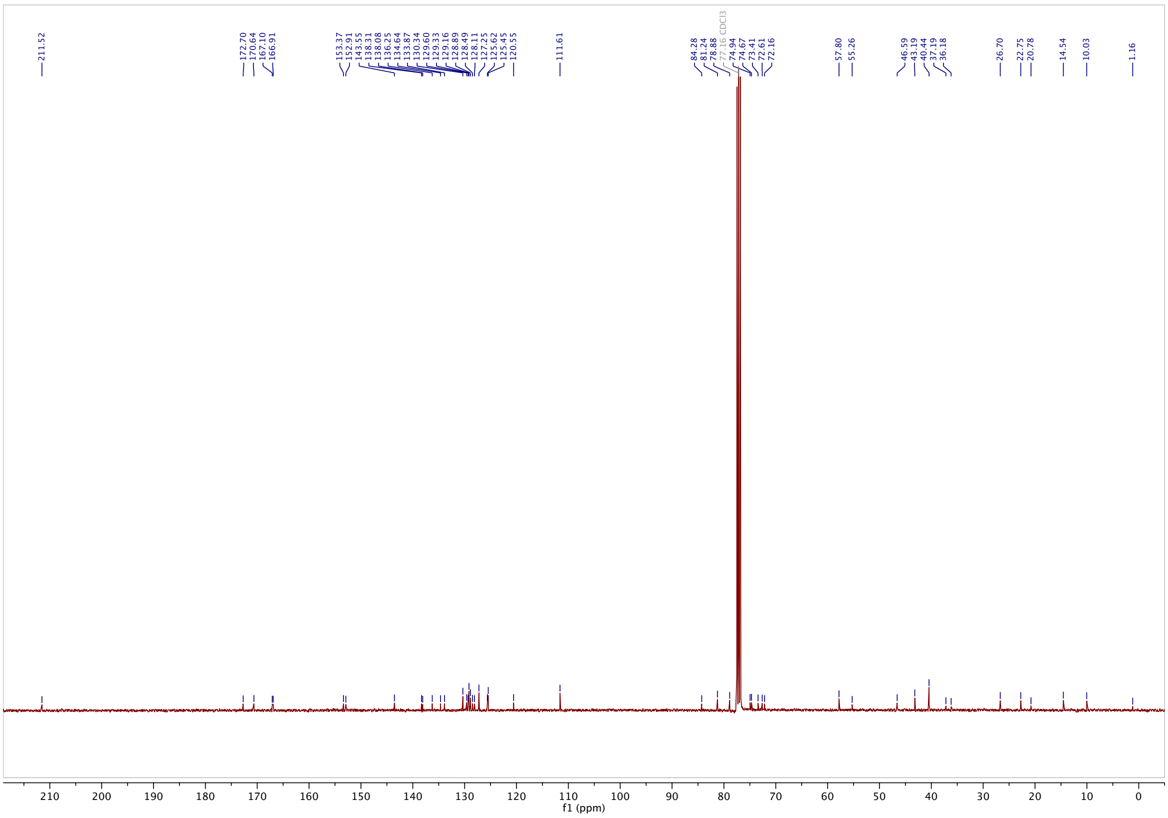

#### AzTax4MP: $^{1}$H-NMR

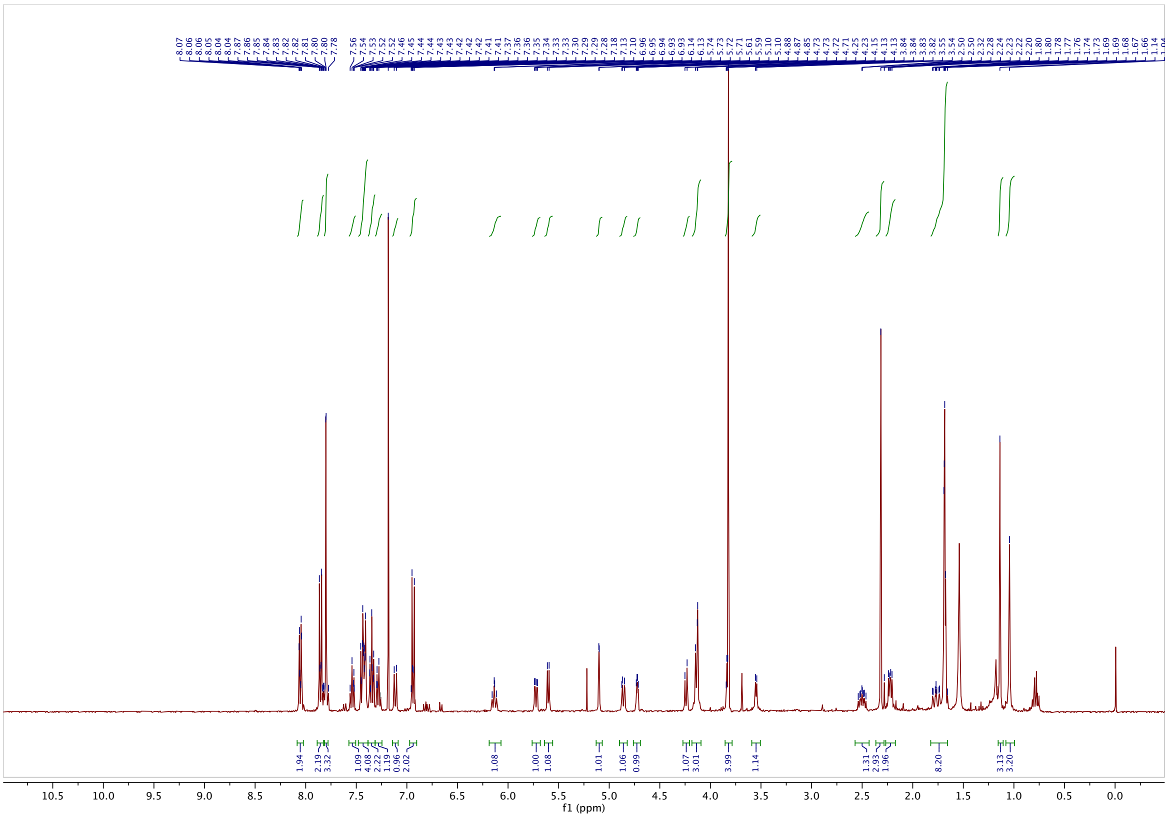

#### $^{13}$C-NMR

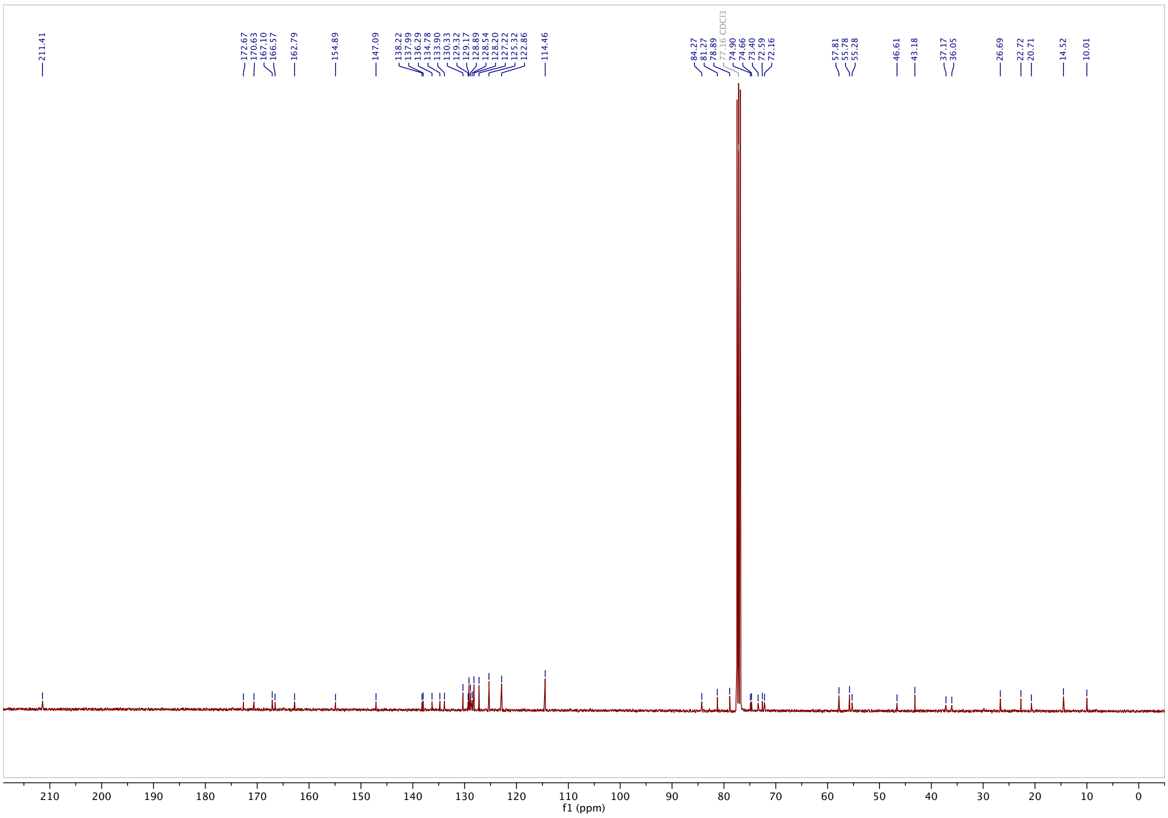

#### **AzTax3DMA**: $^1$H-NMR

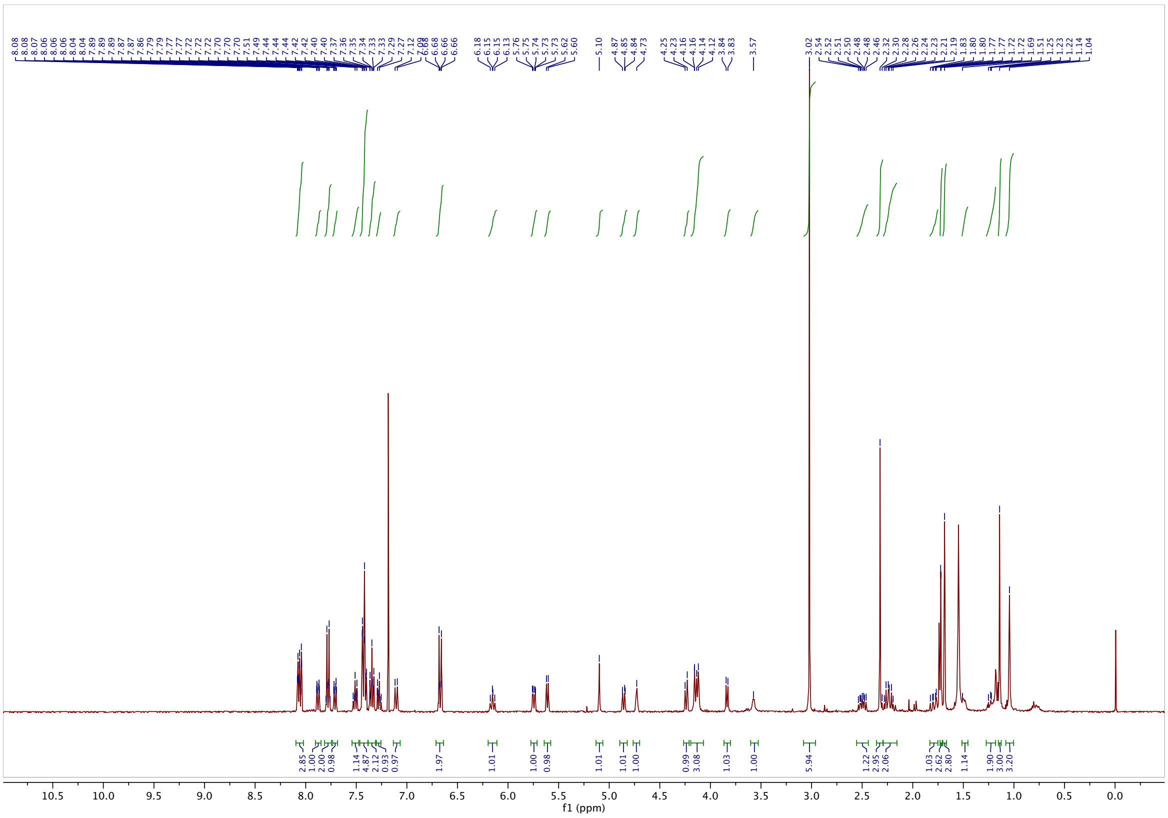

#### $^{13}$C-NMR

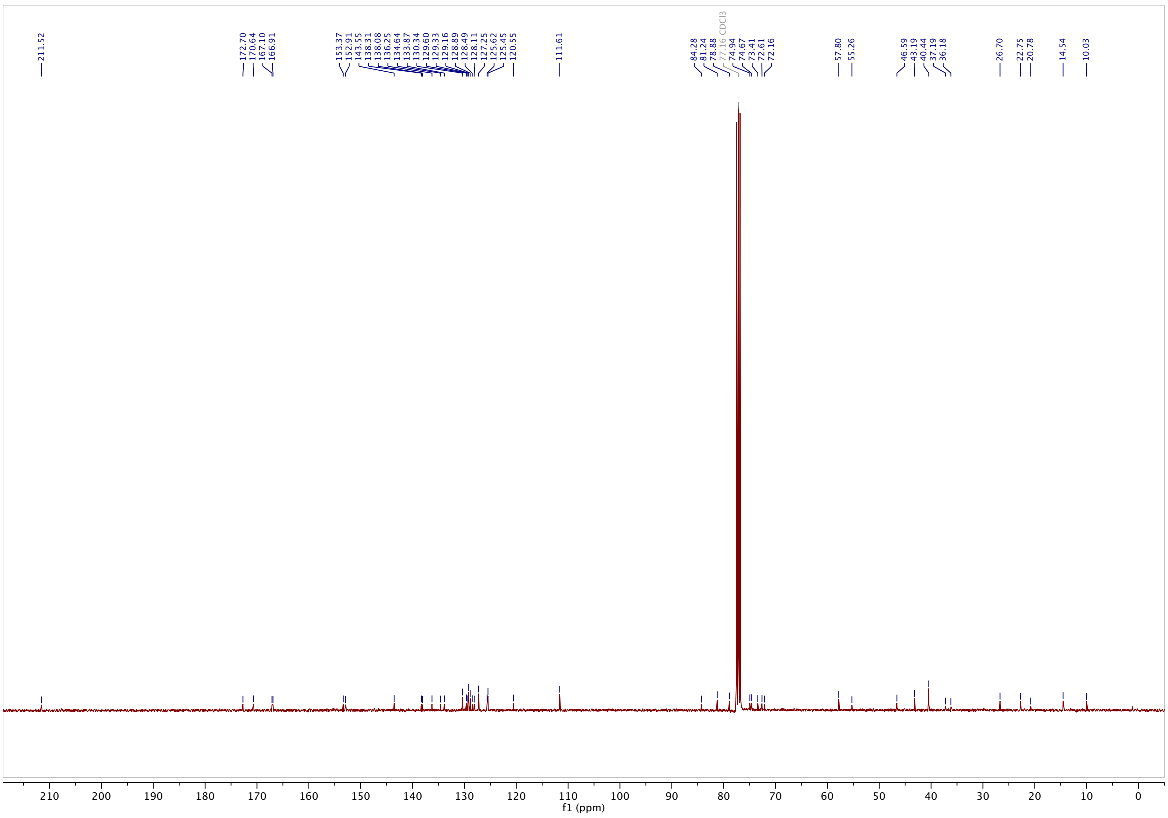

#### AzTax3MP: <sup></sup>1H-NMR

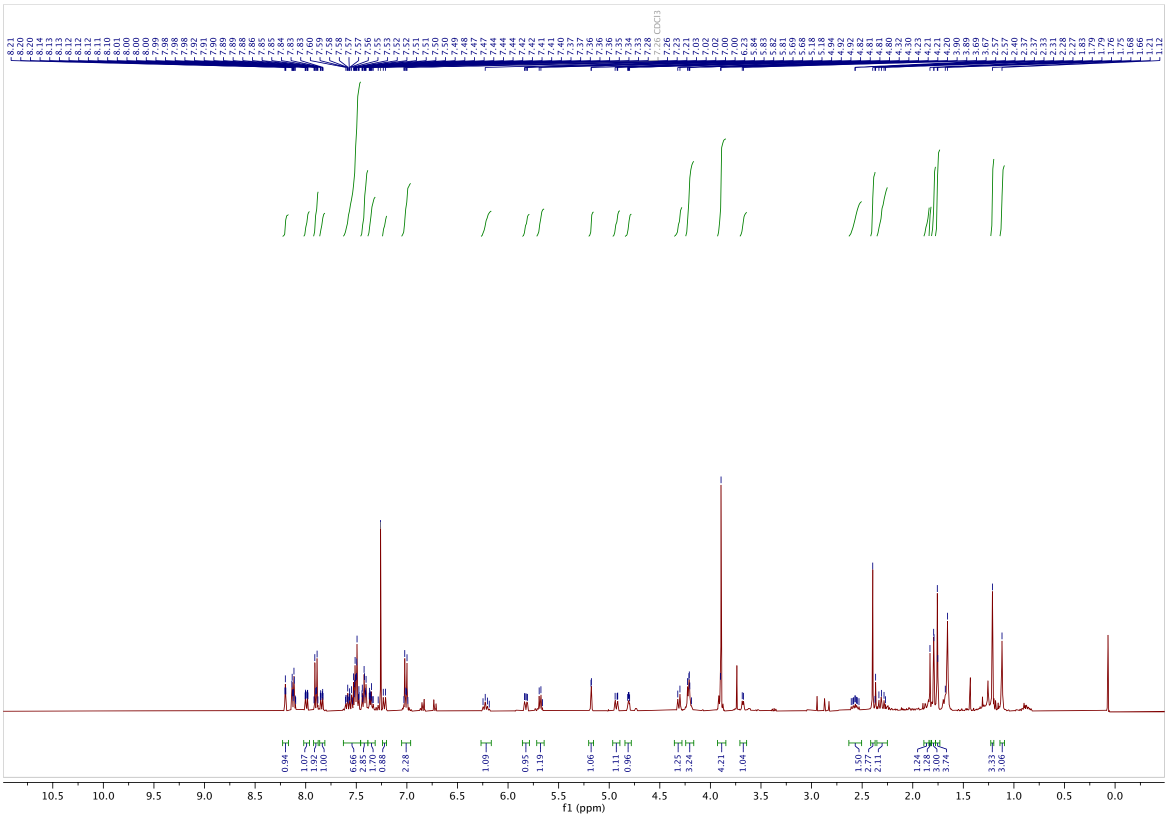

#### 13C-NMR

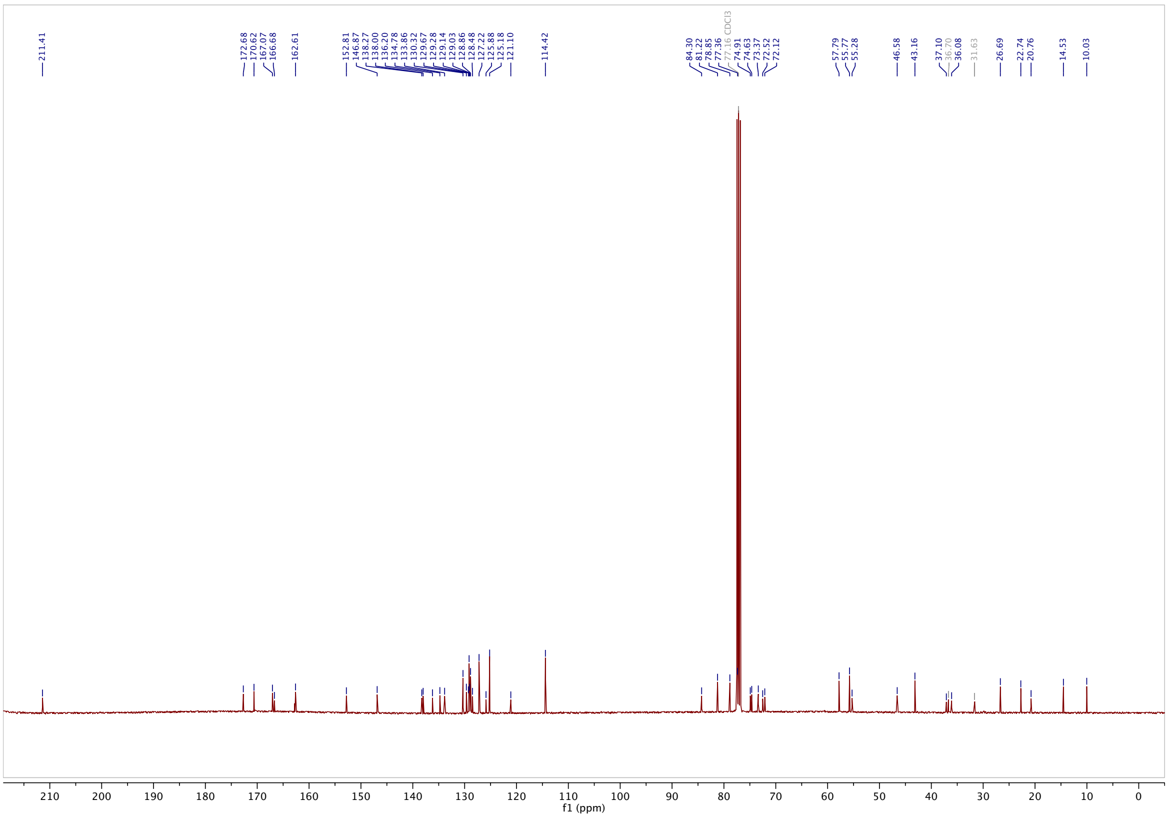

#### AzTax3MTM: $^{1}$H-NMR

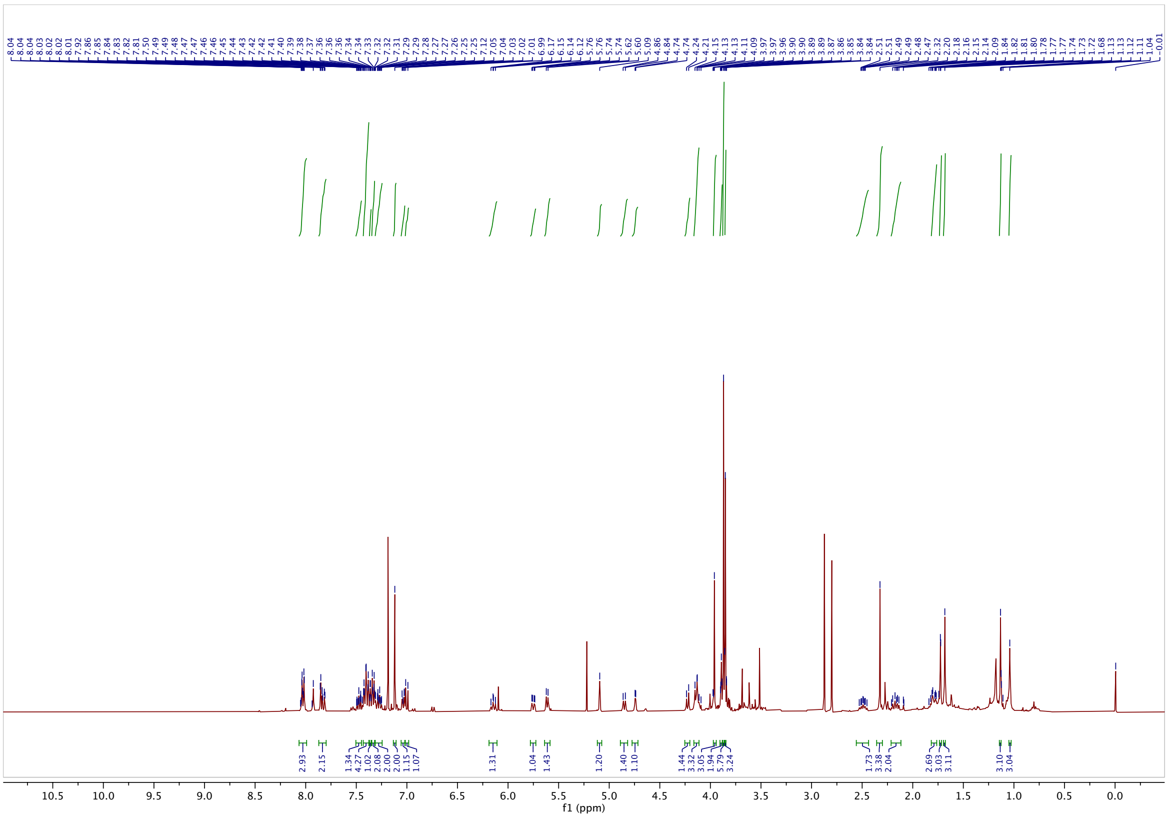

#### $^{13}$C-NMR

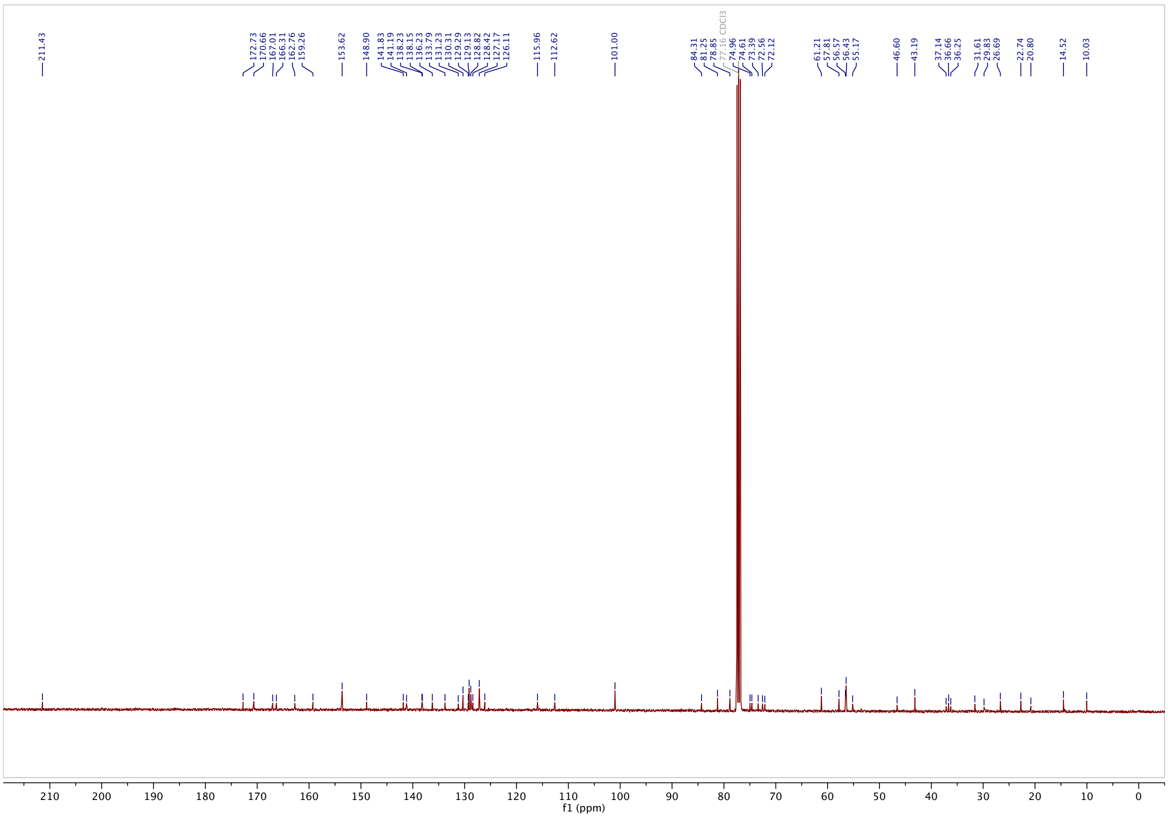

#### AzTax3TM: [1]H-NMR

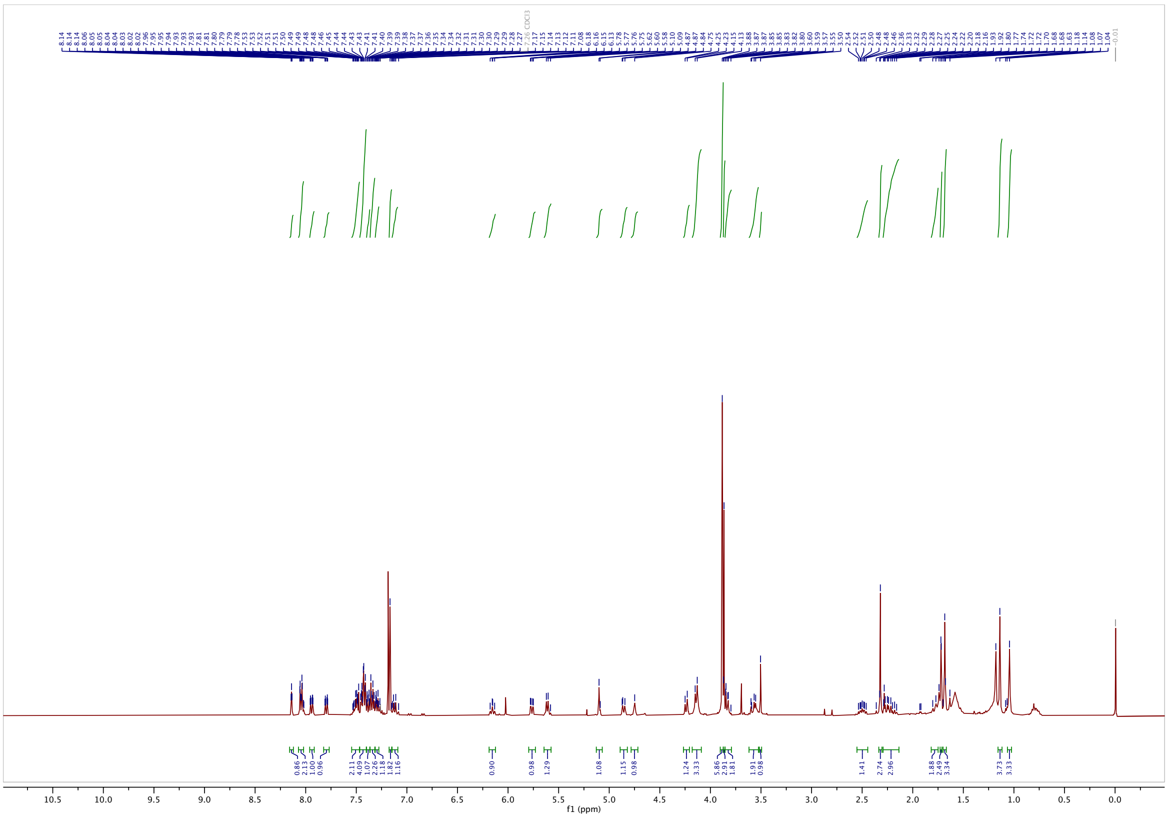

#### [13]C-NMR

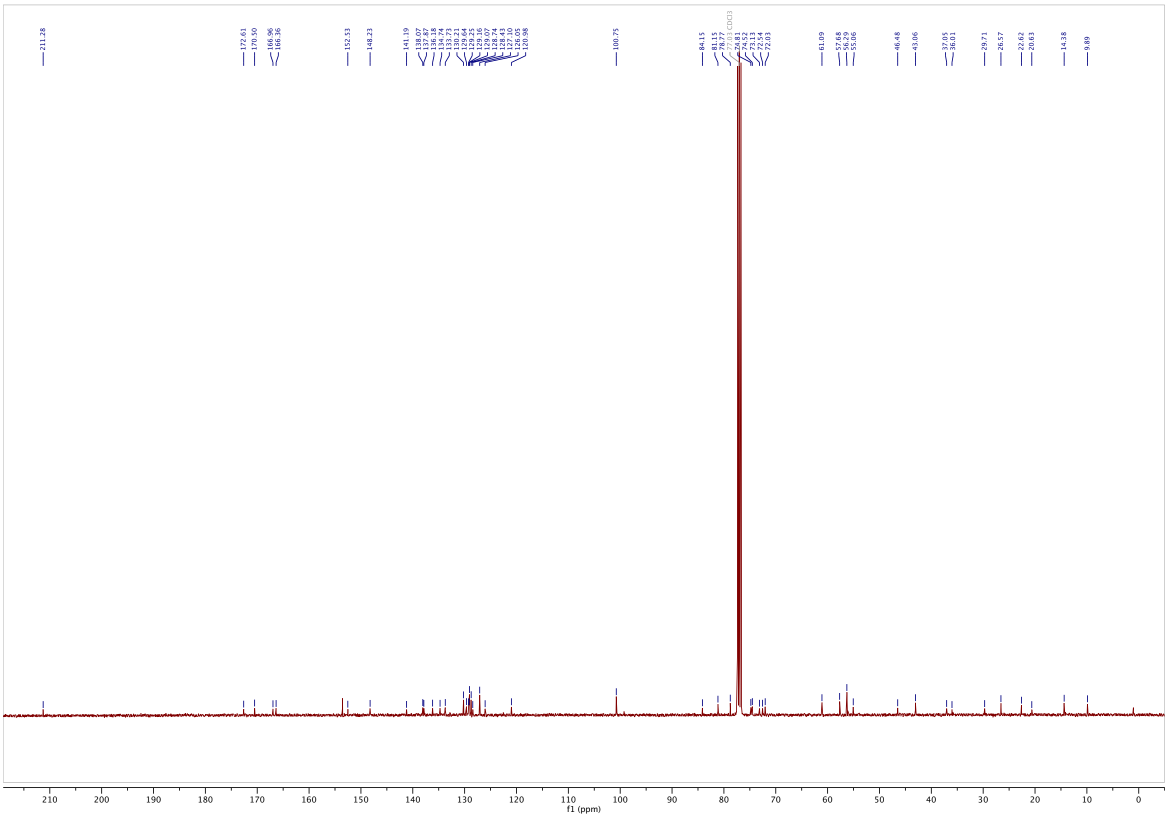

#### AzTax3DEA: [1]H-NMR

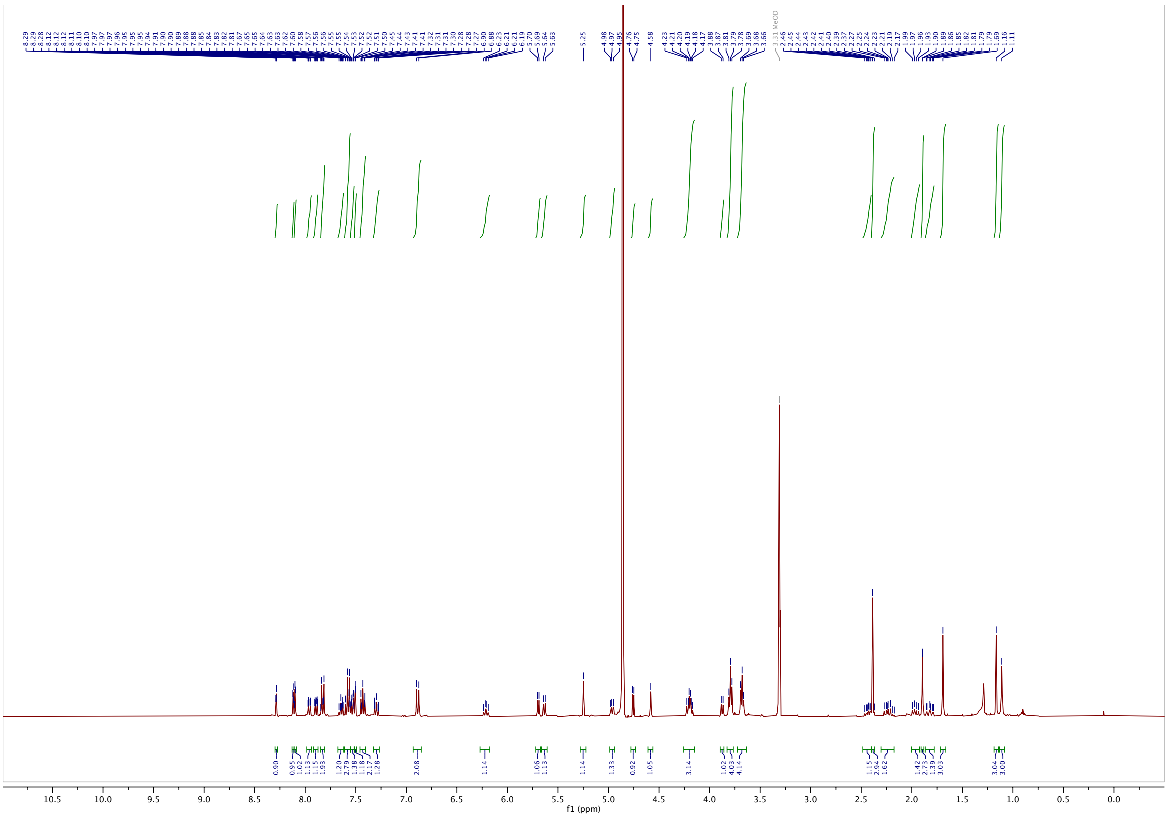

#### [13]C-NMR

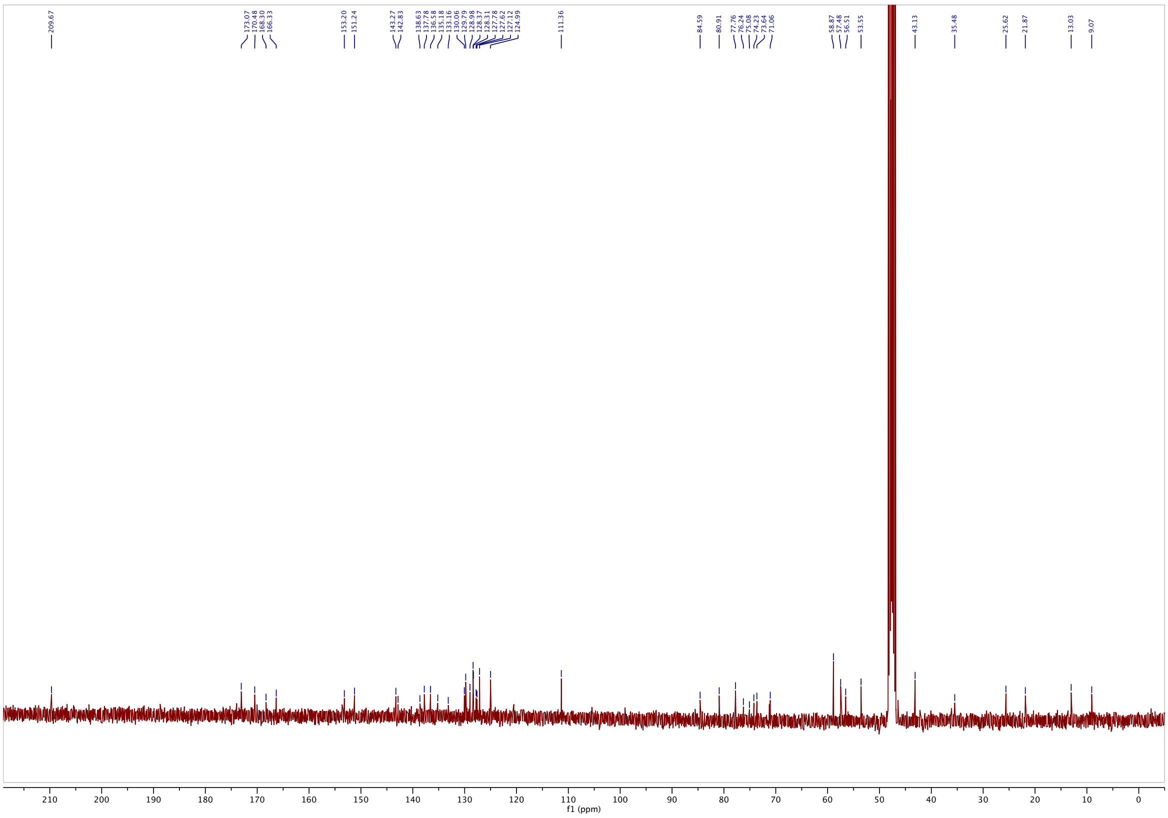

#### **AzTax2MP**: $^{1}$H-NMR

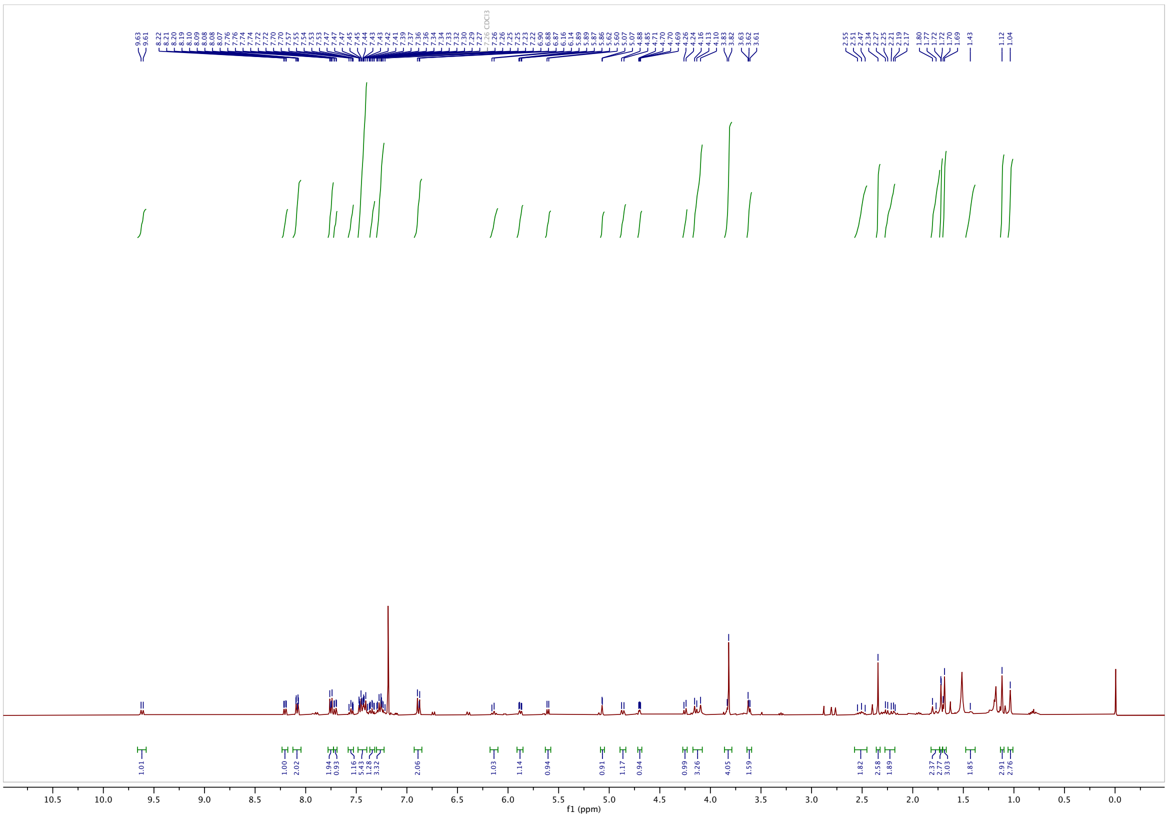

#### $^{13}$C-NMR

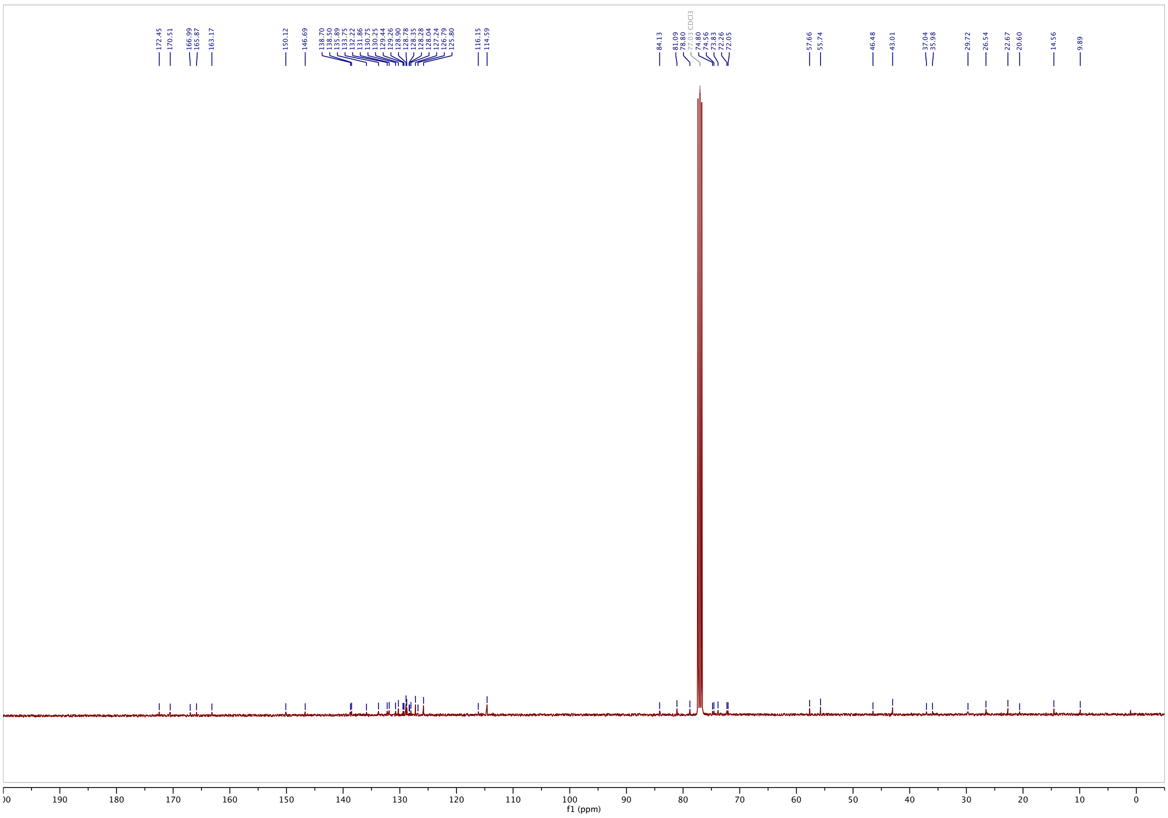

#### *N,N*-bis(2-hydroxyethyl)-4-(phenyldiazenyl)benzamide (4H): $^{1}$H-NMR

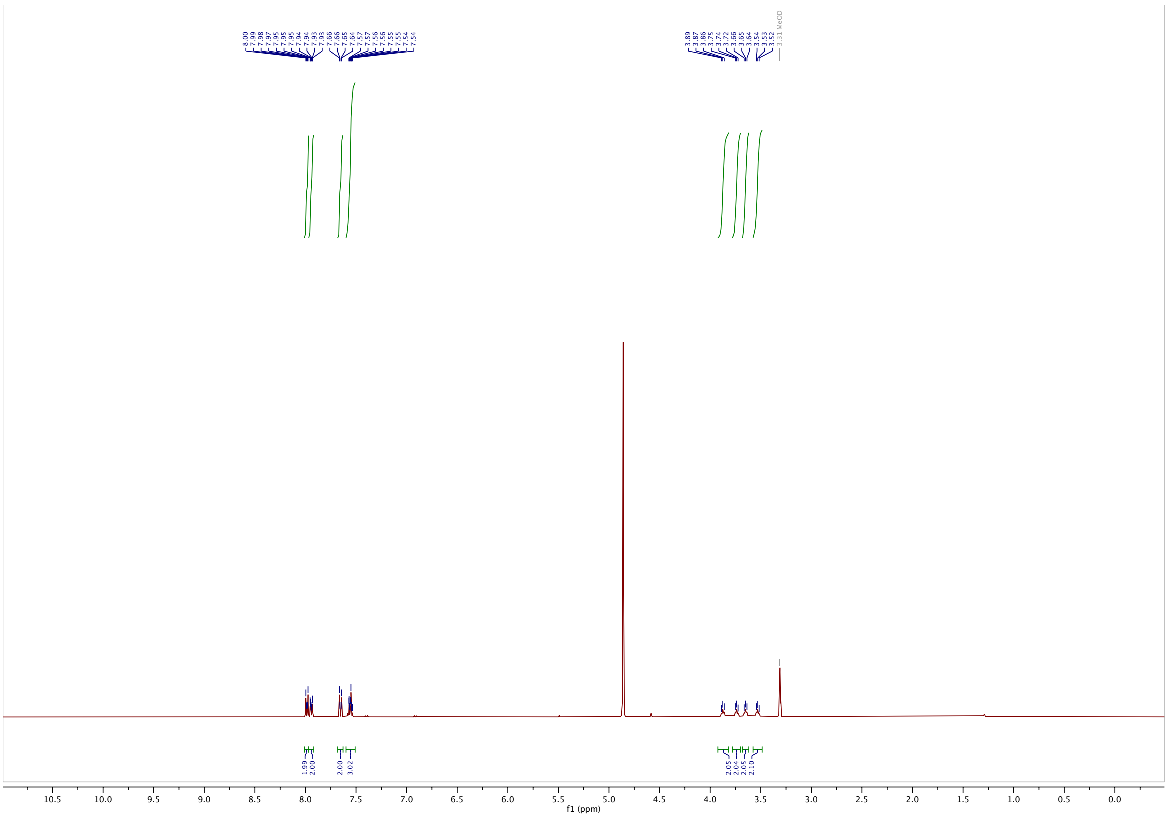

#### $^{13}$C-NMR

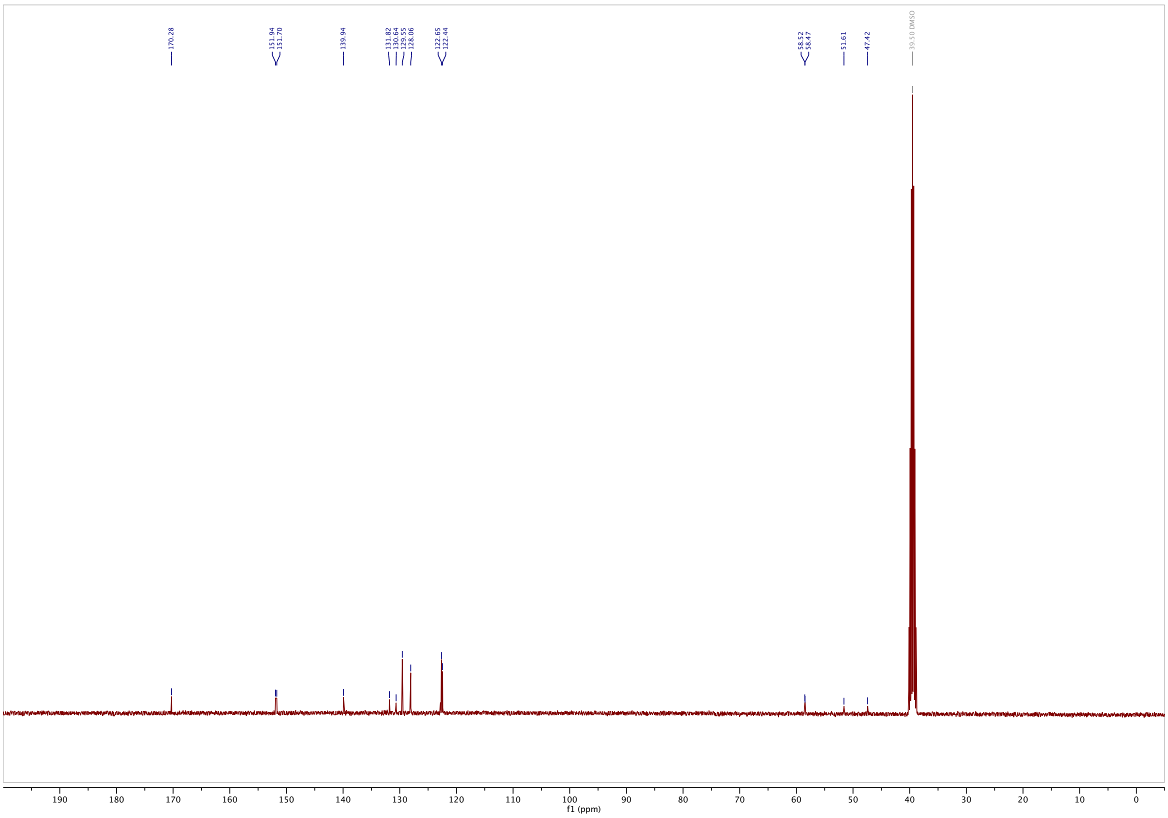

**3-((4-(dimethylamino)phenyl)diazenyl)-*N,N*-bis(2-hydroxyethyl)benzamide (3DMA): $^{1}$H NMR**

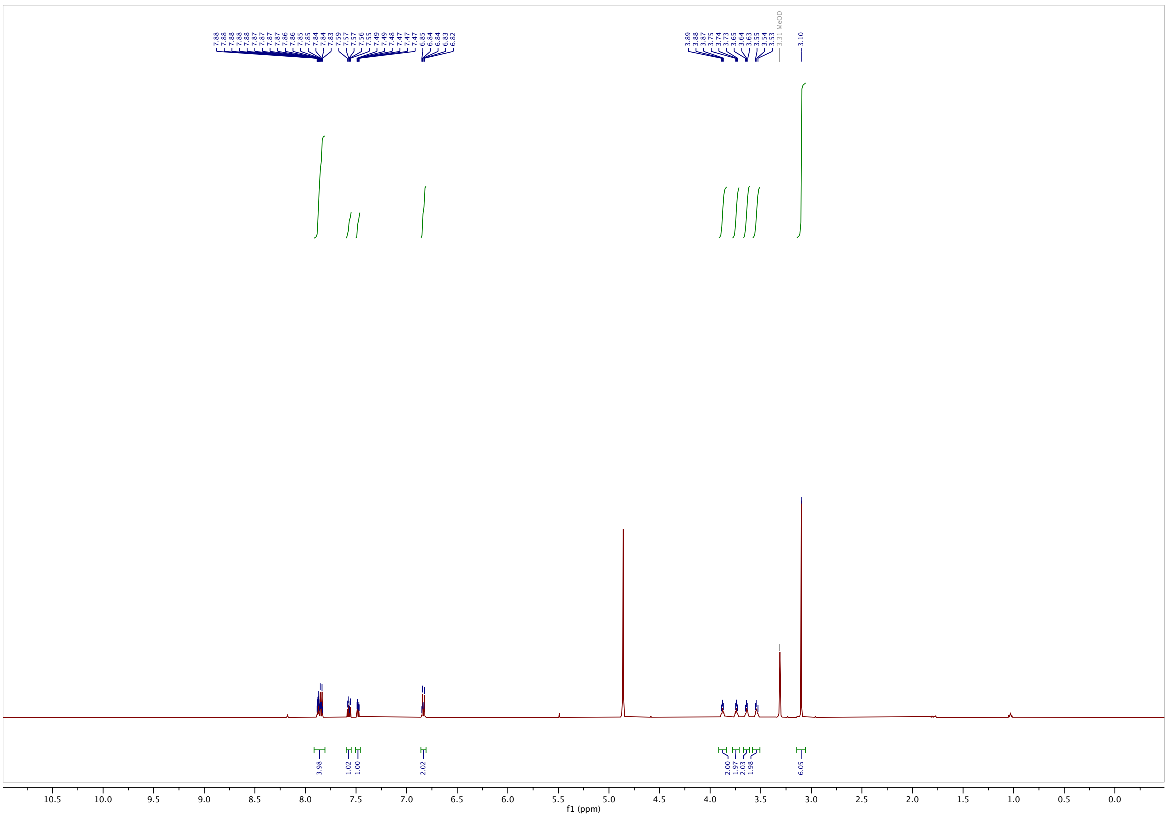

**$^{13}$C-NMR**

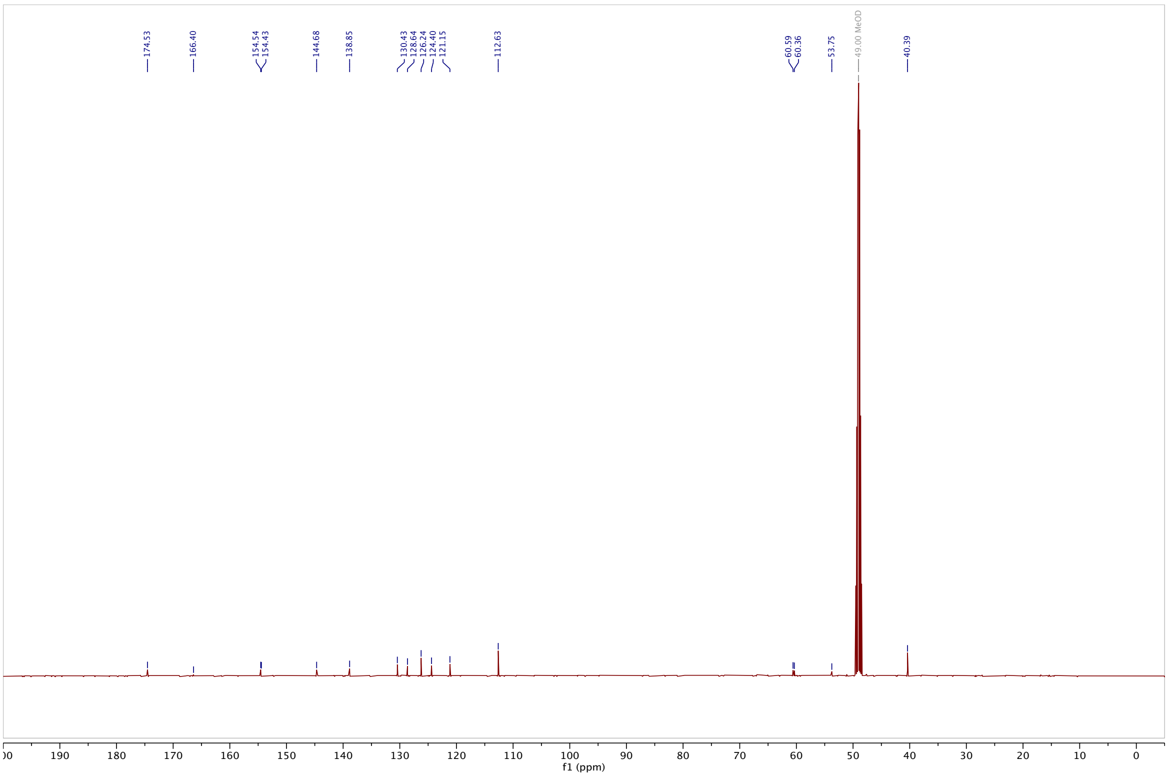

#### *N,N*-bis(2-hydroxyethyl)-3-((4-methoxyphenyl)diazenyl)benzamide (3MP): $^{1}$H-NMR

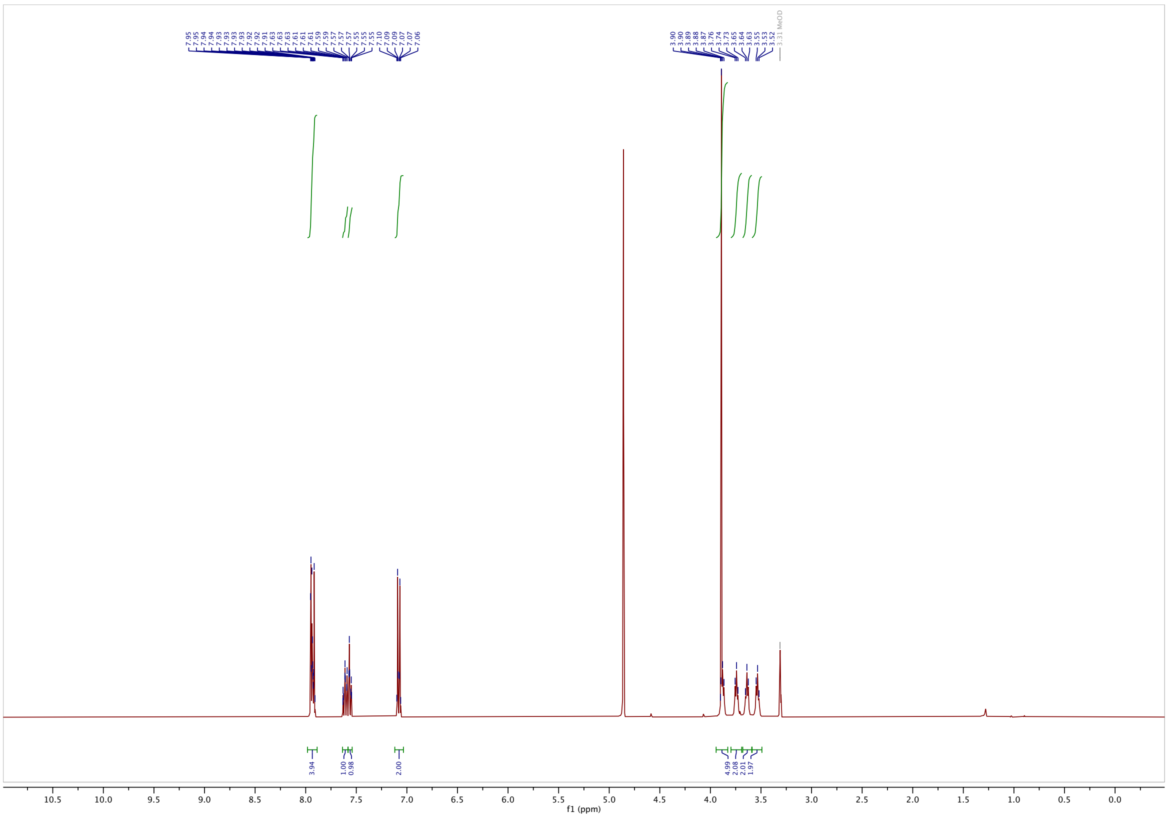

#### $^{13}$C-NMR

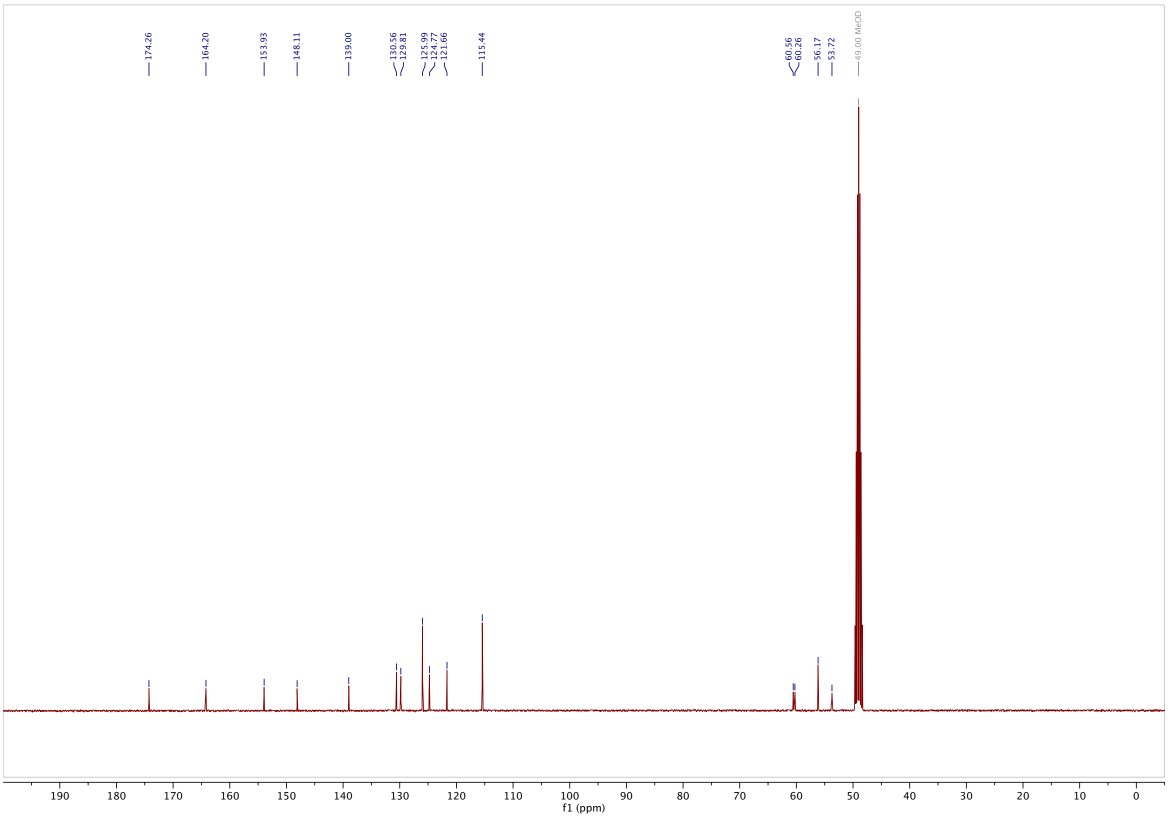
